# Photoreceptor complexity accompanies adaptation to challenging marine environments in Anthozoa

**DOI:** 10.1101/2020.05.28.118018

**Authors:** Sebastian G. Gornik, B. Gideon Bergheim, Nicholas S. Foulkes, Annika Guse

**Affiliations:** Centre for Organismal Studies, Heidelberg University, Heidelberg 69120, Germany; Institute of Biological and Chemical Systems, Karlsruhe Institute of Technology, Hermann-von-Helmholtz Platz 1, Eggenstein-Leopoldshafen 76344, Germany

## Abstract

Light represents a key environmental factor, which shapes the physiology and evolution of most organisms. Notable illustrations of this are reef-building corals (Anthozoa), which adapted to shallow, oligotrophic, tropical oceans by exploiting light from the sun and the moon to regulate various aspects of physiology including sexual reproduction, phototaxis and photosymbiosis. Together with the Medusozoa, (including jellyfish), the Anthozoa constitute the ancestral metazoan phylum cnidaria. While light perception in Medusozoa has received attention, the mechanisms of light sensing in Anthozoa remain largely unknown. Cnidaria express two principle groups of light-sensing proteins: opsins and photolyases/cryptochromes. By inspecting the genomic loci encoding these photoreceptors in over 35 cnidarian species, we reveal that Anthozoa have substantially expanded and diversified their photoreceptor repertoire. We confirm that, in contrast to Medusozoa, which retained one opsin class, anthozoans possess all three urmetazoan opsin classes. We show that anthozoans also evolved an extra sub-group (actinarian ASO-IIs). Strikingly, we reveal that cryptochromes including CRY-IIs are absent in Medusozoa, while the Anthozoa retained these and evolved an additional, novel cryptochrome class (AnthoCRYs), which contain unique tandem duplications of up to 6 copies of the PHR region. We explored the functionality of these photoreceptor groups by structure-function and gene expression analysis in the anthozoan model species *Exaiptasia pallida* (*Aiptasia*), which recapitulates key photo-behaviors of corals. We identified an array of features that we speculate reflect adaptations to shallow aquatic environments, moonlight-induced spawning synchronization and photosymbiosis. We further propose that photoreceptor complexity and diversity in Anthozoa reflects adaptation to challenging habitats.

## Introduction

Light from both the sun and moon dominates the life of many organisms and has had a profound impact on their evolution. While the mechanisms underlying light sensing have been studied in a comparatively small group of animal models, little is known about the impact of light on the physiology and evolution of more ancestral metazoan groups such as the cnidarians. These are basal, non-bilaterian, eumetazoan animals divided into two major groups, the Anthozoa and the Medusozoa (Figure 1A; [1], which both exploit a complexity of sunlight and moonlight-based cues to regulate various aspects of their physiology and behaviour (Figure 1B). Notable examples of highly light-dependent cnidarians are reef-building corals and anemones (both Anthozoa; Figure 1A), many of which live in an evolutionary ancient symbiotic relationship with eukaryotic, photosynthetic dinoflagellates of the Symbiodiniaceae family [2, 3]. The symbionts use sunlight to provide essential photosynthetically-fixed nutrients to their hosts to support host survival in otherwise oligotrophic tropical oceans. In fact, the nutrient transfer from dinoflagellate symbionts to the reef-building corals powers the productivity of reef ecosystems, which are home to more than 25% of all marine species [4]. The majority of these ‘photosynthetic cnidarians’ have a sessile lifestyle in shallow sunlit waters and are mobile only during early development at the larval stage (Figure 1B). Due to this almost ‘plant-like’ lifestyle, sessile cnidarians face similar challenges as true plants such as exposure to intense sunlight, which also bears the risk of temperature stress and UV-induced DNA damage. In addition, most cnidarians harness light from both the sun and moon to orchestrate gamete release during sexual reproduction, including the synchronous mass-spawning events of reef-building corals worldwide [5, 6]. Other important photo-induced behaviours of cnidarians include phototaxis and diurnal migration [7–10]. Given the strong dependence of the cnidarian lifestyle upon environmental lighting conditions, a key question is: which mechanisms mediate these broad ranging effects of light? Given the ecological niches that cnidaria occupy, it is likely that their photoreceptors and light responsive systems would participate in temporally and spatially coordinating behaviour and physiology as well as combating the damaging effects of sunlight. However, with few exceptions including jellyfish eye evolution [11, 12], studies on cryptochrome function in relation to circadian rhythms in *Acropora* and *Nematostella* [13, 14], the light-induced gamete-release in *Clytia hemisphaerica* [15] and a study of opsin evolution in *Hydra* [16], the repertoire and function of cnidarian photoreceptors remains poorly understood.

**Figure 1:**
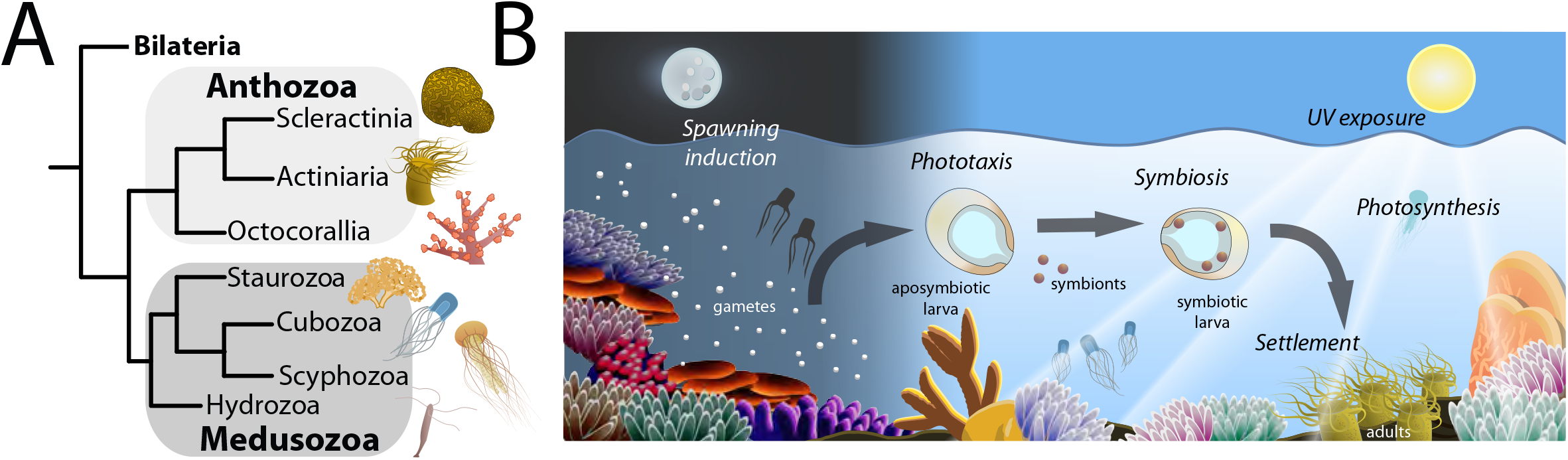
The cnidarian lifestyle is dominated by sun- and moonlight. (A) Cladogram revealing phylogenetic relations within the phylum cnidaria showing the two main classes Anthozoa and Medusozoa and their major subclasses. (B) Schematic overview of how environmental light impacts (symbiotic) cnidarians throughout life from embryo to adult. Light by the moon induces spawning and synchronizes gamete release. Larvae then use environmental light for orientation and during settlement. Adults change behaviour in response to light to modulate photosynthesis rates of symbionts, to seek shelter from UV radiation or for predator avoidance.

Two main groups of light sensing proteins exist in metazoans. Opsins are eumetazoan-specific 7-transmembrane G-protein-coupled receptors, which typically incorporate a retinal chromophore and appear to have evolved from ancestral metazoan hormone-responsive receptors [10, 17–19]. In most animal groups they have been shown to function as membrane-bound photoreceptors that mediate visual as well as non-visual light sensing, and trigger intracellular signalling events upon detection of specific wavelengths. The second group, comprising photolyases (PLs) and cryptochromes (CRYs), is a set of highly conserved flavoproteins involved in harvesting light energy to drive the repair of DNA damage as well as regulating the circadian clock in response to light. Specifically, PLs enzymatically repair pyrimidine-pyrimidone (6-4) and cyclobutane pyrimidine dimer (CPD) DNA lesions generated by UV radiation. They were already present in the common ancestors of Bacteria, Archaea, and Eukarya and are classified according to the type of DNA damage that they repair ((6-4) PLs and CPD-PLs) [20–23]. CRYs, which generally lack photolyase enzyme activity, appear to have evolved independently several times from PLs later during evolution in the Eukarya [23–28]. CRY1s are directly light-sensitive and sometimes also referred to as *Drosophila*-type CRYs [29] while CRY2s (also called vertebrate-type CRYs) exhibit no light-dependent function. They instead regulate clock gene transcription in the negative limb of the circadian clock feedback loop [30, 31]. CRY-DASHs ((*Drosophila*, *Arabidopsis*, *Synechocystis*, Human)-type CRYs) are functionally intermediate between PLs and CRYs and considered photoreceptors with residual DNA repair activity [23, 32]. However, all PLs and CRYs share an amino-terminal <u>ph</u>otolyase-<u>r</u>elated (PHR) region that contains a DNA-binding photolyase domain (also called alpha/beta domain which binds 5,10-methenyltetrahydrofolate (pterin or MTHF)) and a flavin adenine dinucleotide (FAD) domain, which binds to a FAD chromophore [23].

In order to explore the function and evolution of these two photoreceptor protein groups in the cnidarian lineage, we have classified extant cnidarian photoreceptors using a detailed phylogenomics approach. Based on this phylogenetic analysis, we investigate the expression and regulation of cnidarian photoreceptors in the anthozoan symbiosis model *Exaiptasia pallida* (commonly *Aiptasia*) [33]. *Aiptasia* is widely used to investigate the molecular mechanisms underlying cnidarian-dinoflagellate symbiosis establishment and maintenance [34–38] as well as its breakdown, a phenomenon known as ‘coral bleaching’ [39]. Moreover, analogous to most corals, *Aiptasia* also exhibits synchronous, blue (moon) light induced gamete release [40] to produce motile larvae that have to find a suitable niche for their light-dependent lifestyle (Figure 1B). Here we reveal a large and highly diverse photoreceptor repertoire in Anthozoa, and in particular in *Aiptasia*, paving the way to dissecting photoreceptor evolution and function to reveal fundamental principles of how cnidarians adapt to their light-dominated environments.

## Results

### Opsin complexity in Cnidaria

As a first step towards a better understanding of the evolution of the opsin gene family in cnidarians, we generated a detailed molecular phylogeny, based on RNA sequence data and gene structure analysis. Originally, only two types of opsins, the ciliary (c-opsins) and the rhabdomeric (r-opsins) were described [41]. The c-opsins serve as the main visual photoreceptors in vertebrates, while the r-opsins play the same role in invertebrates. Since then, taking advantage of an enormous increase in available sequencing data, new molecular phylogenies and functional studies, more opsin classes have been defined. To date, ten distinct opsin classes have been identified across all Metazoa, three of which, namely the cnidopsins as well as the Anthozoan-specific opsins I (ASO-I) and II (ASO-II) occur in cnidarians (Figure 2A; [42]). The cnidopsins are relatively well studied and are often expressed in a distinct tissue- and stage-specific manner. For example, cnidopsin expression has been studied in the light-sensitive cilia of jellyfish (Medusozoa) eyes, in the hydrozoan battery complex and more ubiquitously in sensory nerve cells [9, 11, 12, 43]. However, far less is known about the anthozoan-specific ASO-I and ASO-IIs; both of which are restricted to Anthozoa (including sea anemones and corals that lack comparable eye-like sensory organs) but are absent in the Medusozoa. Moreover, due to a lack of extensive taxon sampling, a sophisticated assessment of function and diversity for the ASO-I and ASO-II opsins has been lacking. To address this issue, we mined genomic data from 36 cnidarians (7 Anthozoa and 29 Medusozoa). Our new, large-scale phylogeny resolves the ten previously identified distinct opsin classes including the three cnidarian types (Figure 2A). All cnidarians (Anthozoa and Medusozoa) possess cnidopsins, which are monophyletic and sister to the animal xenopsins. The monophyletic ASO-I group appears ancestral to all other opsins and has likely been lost secondarily, similar to the loss of ASO-II opsins in the Medusozoa (Figure 2A;[42]). Moreover, our extended analysis revealed that the ASO-II opsins comprise two distinct, previously not formally described sub-clusters (see Vöcking et al. (2017)). We noticed that one of these two clusters exclusively contains opsins from sea anemones *(Actiniaria*), while the second cluster contains genes from all Anthozoa including sea anemones and corals. This distinct split was confirmed using a second round of reciprocal BLAST searches and phylogenetic inference against a more extensive number of ASO-IIs using data from Picciani et al. (2018) (Figure 2B). Accordingly, we have named the *Actiniaria* (sea anemone)-specific cluster ‘actiniarian ASO-IIs’ while the second cluster remains ASO-IIs. Intron phase analysis corroborates the ASO-II and Actiniarian ASO-II split since we revealed that genes in both clusters have a single intron at distinct positions (Supplementary Figure 1A). Moreover, we observe that the ASO-I and ASO-II intron distribution is distinct from the ctenopsin and c-opsin intron distribution despite their common ancestry since there is a lack of any conserved homologous intron positions (Supplementary Figure 1A). This suggests that the ancestral intron-less ASO-II gene duplicated and subsequently each gene acquired a distinct intron before the ASO-II sub-clusters further diversified.

**Figure 2:**
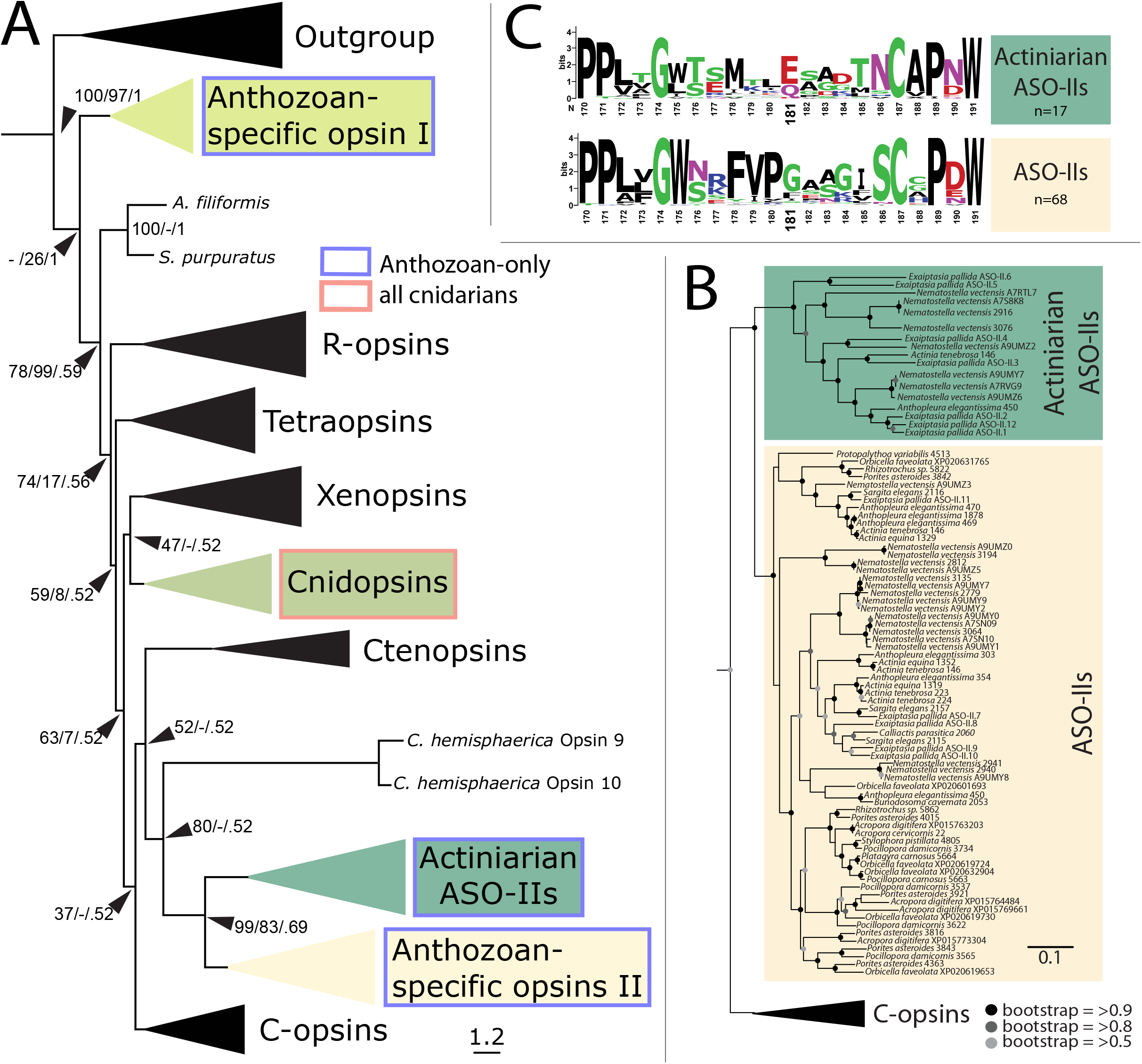
Anthozoa retained the most ancient animal opsin classes and evolved a novel sub-group (actinarian ASO-IIs) (A) Consensus tree topology generated with IQTree reveals that cnidarians have at least 3 opsin paralogs. Anthozoans possess ASO-Is and ASO-IIs, which are monophyletic. ASO-I appears ancestral to all other animal opsins. All cnidarians (Anthozoa and Medusozoa) possess cnidopsins. Cnidopsins are monophyletic also and sister to the animal xenopsins. ASO-IIs contain two distinct sub-clusters. The first cluster comprises solely *Actiniaria* (sea anemone) sequences and we named this grouping ‘Actiniarian ASO-IIs’ while the second cluster contains genes from all Anthozoa families including corals. Intron phase analysis suggests that ASO-IIs and Actiniarian ASO-IIs both are distinct from ctenopsin and c-opsin gene structures despite their common ancestry lacking any conserved homologous intron positions (Supplementary Figure 1A). Note: *C. hemispherica* opsin 9 and opsin 10 resolve at the base of the ASO-IIs, however elsewhere they are included within the cnidopsins albeit with long branches [15]. Support values are (SH-aLRT boostrap percentages/UFBoots boostrap percentages/aBayes Bayesian posterior probabilities). Branch length is proportional to substitutions per site. (B) Bayesian topology tree generated with MrBayes using an expanded ASO-II dataset (Picciani et al. (2018)) confirms the ASO-II/actiniarian ASO-II sub-clustering. Branch length is proportional to substitutions per site. (A) + (B) Full trees can be accessed through Supplementary File 4. (C) Sequence logo showing that most actiniarian ASO-IIs possess a typical primary counter ion (glutamic acid [E]) at position 181; all other ASO-IIs lack this counter ion. The separation of the two ASO-groupings is also reflected in their intron phasing: both ASO-II groupings possess distinct homologous introns (Supplementary Figure 1B).

Changes in amino acid sequence involving key residues is a hallmark of opsin evolution and functional diversification [17, 44–47]. To assess the overall sequence diversity of the opsins occurring in cnidaria, we used pairwise identity mapping of more than 500 opsins (including 237 cnidarian opsins). We found that at the protein level most cnidarian opsins are indeed highly diverse (Supplementary Figure 2). For example, while the ancestral ASO-Is cluster tightly and form one distinct group with a highly similar amino acid composition, the cnidopsins and ASO-IIs are subdivided into various clusters which are interspersed with several classes of opsins associated with higher animals including, for example, vertebrate-specific c-opsins (Supplementary Figure 2). Opsin amino acid composition and photosensory function are highly correlated and specific amino acid residues interact with the chromophore to tune peak spectral sensitivities [44, 48]. Therefore, the high levels of diversity we observe in cnidarian opsins may reflect photosensory diversity rather than being the result of synonymous gene duplications that merely created functionally identical opsin paralogs.

Interestingly, we specifically noted that opsins within the two ASO-II clusters differ in one major functional amino acid. Typically, opsins contain a highly conserved glutamate residue at position 181 (E181), which stabilizes retinal, the light-sensitive chromophore forming a so-called protonated Schiff base (PSB) when bound to opsin. Light absorption triggers retinal *cis*-to-*trans* isomerization, which, in turn, results in opsin conformational changes that reveal a cytoplasmic G-protein binding site and thereby enables the activation of signalling cascades. Free retinal is maximally sensitive to UV light but its absorption maximum is shifted towards visible light when covalently bound to opsins, ensuring its maximal sensitivity lies within the visual spectrum [17]. However, while most actiniarian ASO-IIs, similar to the cnidopsins and ASO-Is, indeed have a glutamate [Q] at the equivalent position, all members of the anthozoan-wide occurring ASO-IIs sub cluster lack this conserved feature (Figure 2C; Supplementary Figure 1B). To date, E181 has been found to be conserved in all opsins [48, 49] with the exception of vertebrate visual opsins where it occurs together with, or is replaced by E113 [17, 47, 50–53], a vertebrate-specific feature associated with even higher fidelity visual photoreception. Interestingly, one *Aiptasia* ASO-II (ASO-II.4) contains both E113 and E181 (Supplementary Figure 1B), suggesting that this presumed vertebrate-specific feature may also have arisen independently in some cnidarians and so may represent an example of convergent evolution conferring higher-fidelity photoreception [53]. Furthermore, these specific amino acid differences are consistent with functional diversification during evolution of the ASO-II opsin group.

### Life-stage, symbiotic state and tissue type-specific opsin expression in Aiptasia

In the model species *Aiptasia,* we identified 18 distinct opsins: 4 cnidopsins XP_020899757.1, XP_020913977.1, XP_020904301.1, XP_020897723.2), 2 ASO-Is (XP_020902074.1, XP_028515325.1) and 5 ASO-IIs (XP_020903100.1, AXN75743.1, XP_020897790.2, XP_028514120.1, XP_020909716.1) and 7 actiniarian ASO-IIs XP_020914799.1, XP_020906239.1, XP_020909580.1, XP_020907384.1, XP_020910007.1, XP_020893775.2, XP_020914907.2). See also: Supplementary File 1. To assess whether this broad opsin repertoire is actively expressed and if so, during which life stages, we compared opsin expression levels using publicly available *Aiptasia* RNA-Seq data [34, 54]. We found that with one exception, all *Aiptasia* opsins are ubiquitously expressed in both larvae and adults with expression levels of some opsins elevated specifically in adults and others during larval stages suggesting the existence of opsins with larval- and adult-specific functions (Figure 3A). Likewise, opsin expression varies depending on the symbiotic state (Figure 3B). For example, one opsin from the actiniarian-specific ASO-II group (ASO-II.12, dark green), and two from the ASO-II group (ASO-II.7 and ASO-II.11, yellow) show significantly higher expression levels in symbiotic anemones when compared to their aposymbiotic (non-symbiotic) counterparts. This is consistent with previous findings that symbiotic association influences photo-movement in *Aiptasia* [8] and suggests that host perception of environmental light by opsin-mediated light-sensing may change in response to symbiosis, for example to adjust the levels of sunlight exposure for optimal photosynthesis rates.

**Figure 3:**
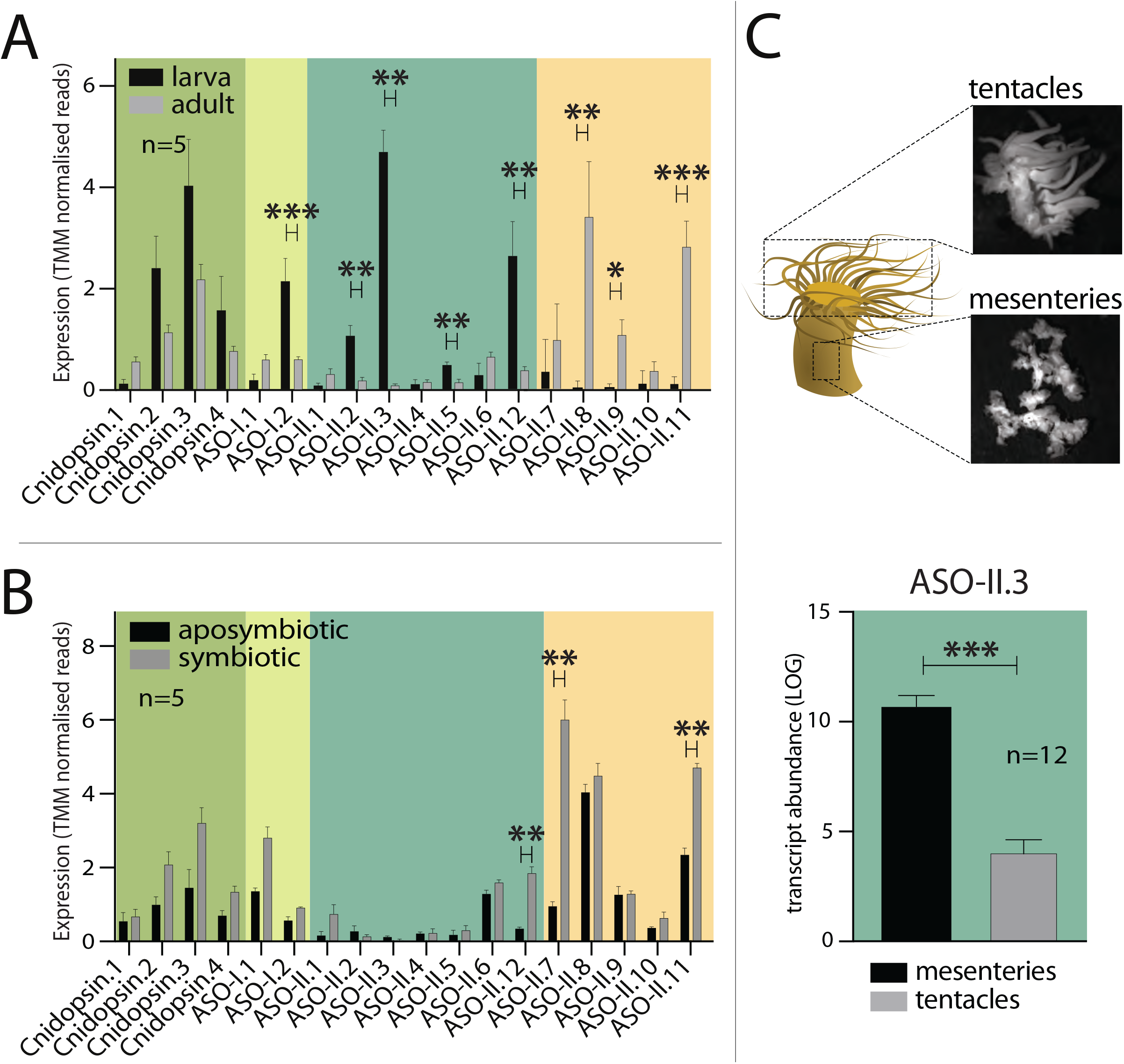
Expression profiles of *Aiptasia* opsins varies between life-stages and symbiotic state. (A) Bar chart comparing the expression (TMM normalised reads) of *Aiptasia* opsins in adults and larva. ASO-I.2, ASO-II.2, ASO-II.3, ASO-II.5 and ASO-II.12 are significantly upregulated in larva. ASO-II.8, ASO-II.9 and ASO-II.11 are significantly upregulated in adults. (B) Bar chart comparing the expression (TMM normalised reads) of *Aiptasia* opsins in symbiotic and aposymbiotic adults. ASO-II.12, ASO-II.7 and ASO-II.11 are significantly upregulated in symbiotic adults. (C) Tissue-specific qPCR after Methacarn fixation of adult *Aiptasia* polyps reveals that ASO-II.3 is significantly upregulated in mesenteries, when compared with tentacle tissue. Data for all other opsins are shown in Supplementary Figure 3. For all charts significant differences are: * P ≤ 0.05, ** P ≤ 0.01, *** P ≤ 0.001; and error bars are: SEM.

Opsins have been implicated in ‘sensing’ moonlight to synchronize gamete release in corals [5, 55]. Specifically, it is predicted that physiologically relevant blue shifts in the irradiance spectrum measurable during twilight on several days before and after the full moon acts as a trigger for a potential opsin-mediated dichromatic visual system where readouts from a blue- and red-light sensitive opsin are integrated to induce spawning [6]. Accordingly, exposure for 5 nights to LED-based blue light has been shown to specifically induce synchronized spawning in *Aiptasia* [40]. By analogy with the jellyfish *Clytia hemisphaerica* in which the opsin relevant for spawning is specifically expressed in the gonadal tissue [15], we therefore asked whether any of the opsin genes present in *Aiptasia* showed a gonad-specific expression pattern (Supplementary Figure 3). By using qPCR analysis, we revealed that *Aiptasia* ASO-II.3 is indeed expressed in a tissue-specific manner and significantly up-regulated in mesenteries when compared to the tentacles (Figure 3C). Thus, *Aiptasia* ASO-II.3 represents a candidate opsin that may be involved in spawning induction in this species.

### Novelty in the cnidarian photolyase and cryptochrome repertoire

Another major group of light-sensing proteins in animals are the CRY and PL flavoproteins, however, to date their phylogeny in cnidarians has not been assessed in detail. To address this issue, we next used phylogenomic analysis and considered the position of conserved introns. We revealed that while they possess CRY-DASH, CPD-II PLs and (6-4) PLs, CPD-Is and CRY-Is are absent from all cnidarians (Figure 4A, Supplementary Figure 4A). Furthermore, while CRY-IIs are encountered only in the subphylum Anthozoa, strikingly CRYs are completely absent from the Medusozoa that are represented by 29 taxa in our analysis (Supplementary File 5).

**Figure 4:**
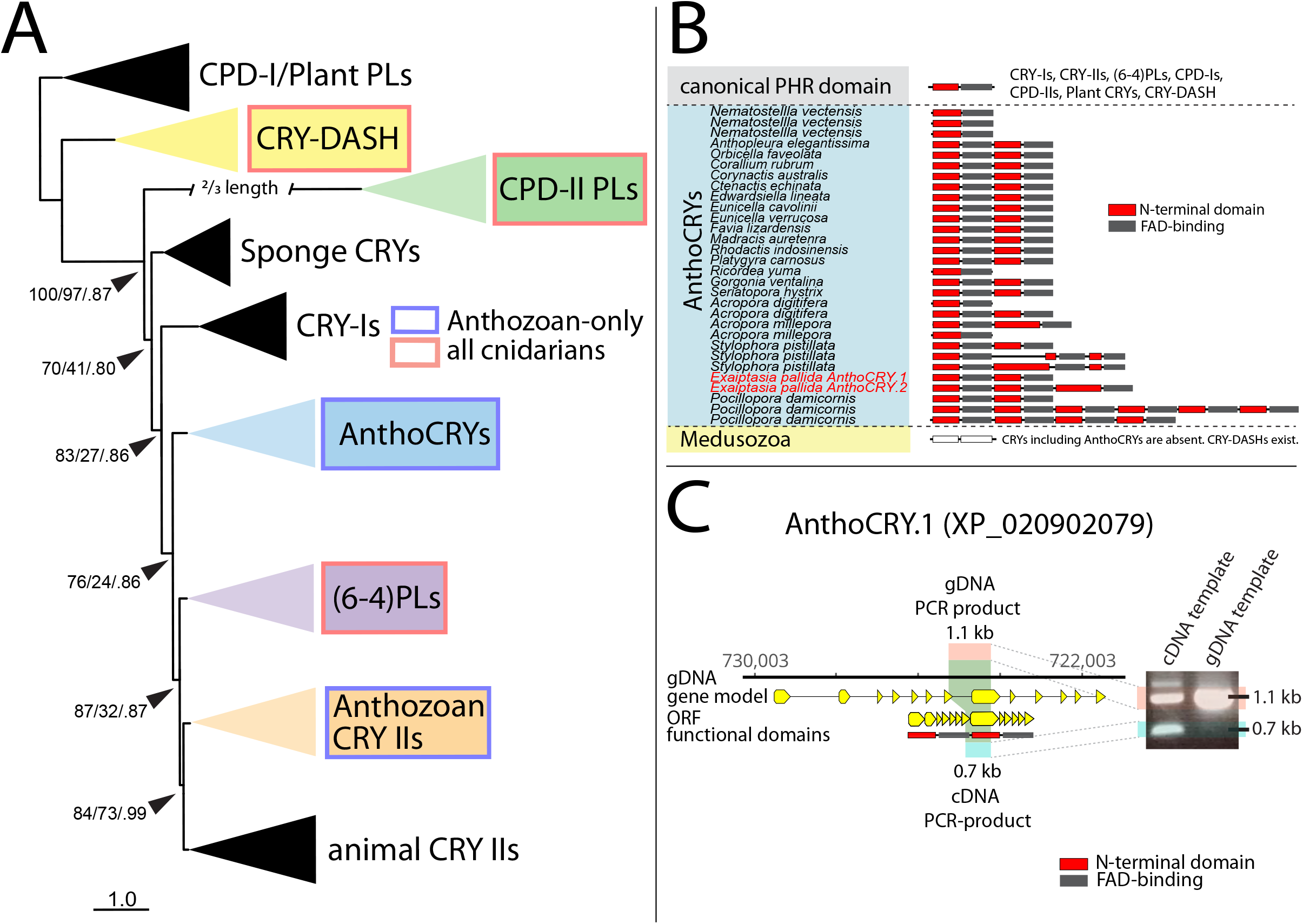
PL and CRY phylogeny reveals a novel, Anthozoan-specific CRY class. (A) Consensus tree topology generated with IQTree reveals that PLs (CPD-II, PLs and (6-4) PLs) and CRY-DASHs are common amongst all Cnidaria. Other CRYs including CRY-II occur only in Anthozoa but are entirely absent from Medusozoa. The present phylogeny is supported by highly conserved and specific exon-intron patterns (Supplementary Figure X) suggesting that CRY-IIs and (6-4) PLs share a common origin. A well supported but previously unidentified, anthozoan-specific CRY family resolves at the base of animal CRY-IIs and (6-4) PLs which we name Anthozoan-specific CRYs (AnthoCRYs). Support values are (SH-aLRT boostrap percentages/UFBoots boostrap percentages/aBayes Bayesian posterior probabilities). A full tree can be accessed through Supplementary File 5. (B) Cryptochromes (except CRY-DASH) are absent in Medusozoa, while the Anthozoa possess a novel class of cryptochromes (AnthoCRYs), which contain unique tandem duplications including up to 6 copies of the PHR region comprising the N-terminal DNA-binding photolyase domain (also called alpha/beta domain, red) and the chromophore-binding FAD domain (grey). (C) A comparison between cDNA- and genomic DNA-derived exon spanning amplicon sequencing confirms PHR region duplication in *Aiptasia* AnthoCRY.1 (XP_020902079).

Interestingly, we have identified two distinct CRY groups in Anthozoa. One is a sister group to ‘animal CRY-IIs’ that we named anthozoan CRY-IIs. Due to their phylogenetic position at the base of animal CRY-IIs we speculate these are likely to be involved in circadian clock function [29, 56, 57]. However, a second, novel CRY group appears to be basal to both (6-4) PLs and ‘animal CRY-IIs’ (but distinct from CRY-Is and sponge CRYs). We thus named this group Anthozoan-specific CRYs (AnthoCRYs) (Figure 4A and 4B). Previous, preliminary analysis of both cryptochrome groups lead to them being classified within the animal CRY-IIs, presumably due to a lack of cnidarian taxa representation in associated phylogenies [13, 58, 59]. Our study now clarifies their phylogenetic position and proposes their name based on identity. Thus, similar to the situation for the opsin genes, the Anthozoa possess a more extensive repertoire of CRYs when compared with the Medusozoa.

A hallmark of PLs and CRYs is the PL-homologous region (PHR region) that contains a N-terminal DNA-binding photolyase domain (also called the alpha/beta domain) and a C-terminal FAD chromophore binding domain [21, 22, 57]. All PLs and CRYs described to date contain a single PHR region. Strikingly, however, here we reveal that AnthoCRYs contain up to six tandemly repeated PHR regions (Figure 4B). We confirmed this PHR region duplication independently in the sea anemone *Aiptasia* by PCR and sequencing (Figure 4C) of the *Aiptasia* AnthoCRY.1 cDNA. Such a PHR region expansion has not been described for any PL or CRY to date. This expansion occurs across all Anthozoa including non-symbiotic and symbiotic members indicating that the tandem duplication of PHR domains in Anthozoan CRYs might serve a common purpose in this animal group. Interestingly, it appears to be absent from *Nematostella*.

### Light regulated CRY gene expression in Cnidaria

We next wished to investigate the functionality of these flavoprotein genes in terms of their regulation following light exposure. In many animal groups the expression of CRY and PL genes is induced upon exposure to light as a key mechanism to regulate the circadian clock or to upregulate DNA repair capacity in response to prolonged sunlight exposure. Consistently, in *Nematostella* three CRYs have been reported, two of which are upregulated in response to light [59]. Similarly, at least two different CRYs (Cry1a (XP_001631029) and Cry1b (XP_001632849)) are expressed in a light dependant manner in the coral *A. millepora* [13]. In the sea anemone *Aiptasia diaphana* two CRYs were identified (without accession numbers) and shown to be expressed rhythmically in the presence of a day-night cycle [60]. We therefore explored to which extent light regulates the 8 different *Aiptasia* PL and CRY genes that we identified (2 CPD_II isoforms (XP_020910442.1, XP_020910516.1), 1 CRY-DASH XP_020903321.1), 2 (6-4) PL isoforms (XP_020915076.1, XP_020915067.1), 1 Anthozoan CRY (XP_020904995.1) and 2 AnthoCRYs (XP_020902079.1, XP_020917737.1; see also: Supplementary File 2). With the exception of AnthoCRY.2, where mRNA levels are undetectable in *Aiptasia* larvae, we showed that all CRY and PL genes are generally ubiquitously expressed in both *Aiptasia* larvae and adults (Supplementary Figure 4B). We next adapted *Aiptasia* for 4 days to constant darkness and then exposed them for a period of 8 hours to light, sampling at 2-hours intervals. Our results revealed that the expression of *Aiptasia* PLs and CRYs was differentially affected by light exposure. Specifically, *Aiptasia* CPD-II, *Aiptasia* DASH-CRY and *Aiptasia* (6-4) PL were largely unresponsive to light treatment (Figure 5A to 5C). In contrast, AnthoCRYs and CRY-II expression was rapidly induced upon exposure to light (Figure 5D to 5F).

**Figure 5:**
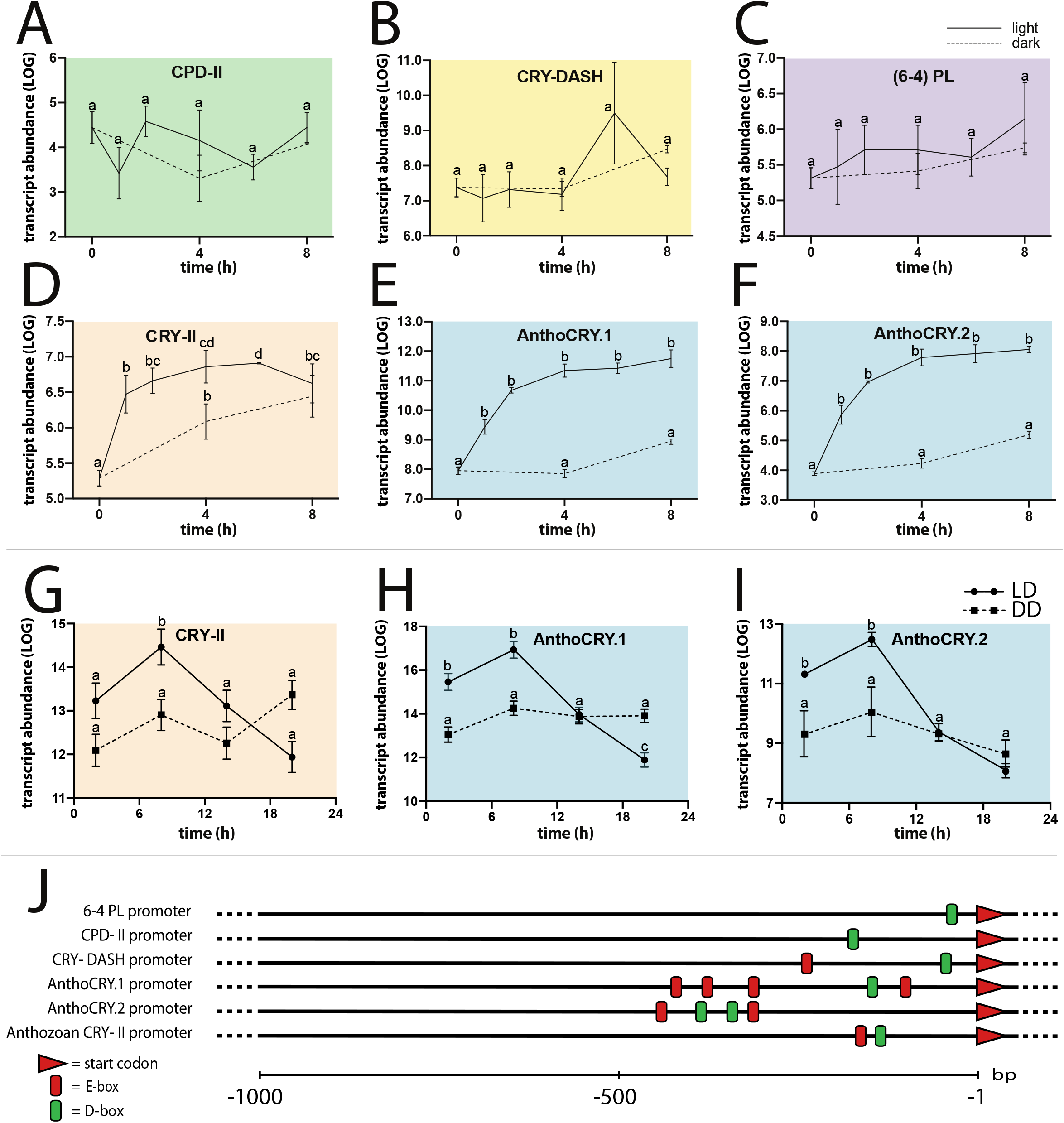
Light-induced expression profiles of *Aiptasia* CRYs and PLs. (A) – (F) qPCR analysis of CRY and PL gene expression in response to light exposure in adult *Aiptasia*. Animals were adapted for 4 days to constant darkness and then exposed to light for a period of 8 hours; sampling every 2 hours. Control animals were kept in constant darkness. *Aiptasia* CPD-II, *Aiptasia* DASH-CRY and *Aiptasia* (6-4) PL are unresponsive to light treatment. AnthoCRY.1, AnthoCRY.1 and CRY-II expression is rapidly induced by light. (G) - (I) qPCR analysis of CRY-II, AnthoCRY.1 and AnthoCRY.2 expression in LD-adapted *Aiptasia* polyps. Animals were exposed to light-dark (LD, 12 hours:12 hours) cycles and then transferred to constant darkness (DD), sampling at 2, 8, 14 and 20 hours (LD) and 26, 32, 38 and 44 hours (DD). Under LD conditions, we observed rhythmic expression with elevated expression, peaking at 8 hours after lights on, and then decreasing during the dark period, with a trough at 8 hours after lights off. In contrast, immediately upon transfer to DD conditions, rhythmic expression was absent. (A) - (I) ANOVA was performed to confirm statistically significant differences at each time point; P < 0.01. (J) Schematic representation of the D- and E-box distribution (green and red boxes respectively) in *Aiptasia* PL and CRY promoter regions extending up to 1000 bp upstream from the ATG start codons.

A light-inducible expression pattern may be the consequence of acutely light-driven gene expression or alternatively of regulation by the circadian clock that is also likely to be synchronized during the period of light exposure. To distinguish between these two possibilities, we tested AnthoCRY and CRY-II expression by exposing *Aiptasia* to light-dark (LD) cycles and then transferring animals to constant darkness (DD). Clock regulation would be revealed by the appearance of rhythmic expression under an LD cycle that would persist following transfer to constant darkness. Therefore, samples were prepared at 6 hours intervals during either exposure to an LD cycle or immediately following transfer from LD to DD conditions and then CRY gene expression was assayed by qPCR. Under LD conditions we observed rhythmic expression with elevated expression during the light period, peaking at 8 hours after lights on, and then decreasing during the dark period, with a trough at 8 hours after lights off (Figure 5G to 5I). In contrast, immediately upon transfer to DD conditions, rhythmic expression was absent, showing that changes in CRY gene expression are indeed light-, rather than clock-driven (Figure 5G to 5I).

We have previously studied the mechanisms underlying light-driven gene expression of cryptochromes and photolyases in vertebrates [61, 62]. We revealed a conserved role for light-induced transcription of these genes, mediated by the D-box element, an enhancer that has also been associated with circadian clock regulation together with the E-box enhancer [63]. We therefore tested whether D-box or E-box enhancer elements might be encountered within the promoter regions of the various *Aiptasia* CRY and PL genes. By scanning the genomic regions 1 kb upstream of the respective gene START codon (ATG), single D-box enhancers were identified in 6-4 PL and CPD-II, two genes which were not induced upon exposure to light. However, in the case of the light-inducible AnthoCRY and CRY-II genes we identified several PAR/bZIP binding sites (D-boxes; Figure 5J), interestingly, located proximally to E-box enhancers. Furthermore, consistent with the existence of functional D-box regulatory pathways in Anthozoa, we show that *Aiptasia* possesses 4 putative orthologs of the PAR/bZIP TFs family (three TFs basal to <u>H</u>epatic <u>l</u>eukemia <u>f</u>actor (HLF)-type PAR/bZIP TFs and one DBP (<u>D</u>-box <u>B</u>inding <u>P</u>AR/bZIP) TF (Supplementary Figure 5) which have been shown to bind to and regulate transcription from the D-box enhancer in vertebrates [64–66].

## Discussion

Reef-building corals and sea anemones represent critically important members of the ecosystems in shallow, oligotrophic, tropical oceans. Their physiology is dominated by light. They exploit sunlight and moonlight to regulate their sexual reproduction, phototaxis and photosymbiosis. Furthermore, there exposure to sustained high levels of sunlight puts them at particular risk from elevated levels of DNA damage. In order to explore the molecular mechanisms linking light with anthozoan biology, we present the first detailed phylogenetic analysis of two major light sensing protein groups: the opsins and the cryptochrome/photolyase flavoproteins. Within the broader context of the ancestral metazoan phylum, the cnidaria, we reveal that the Anthozoa have substantially expanded and diversified their photoreceptor repertoire compared with the Medusozoa. This striking observation raises several fundamental questions concerning how this expanded photoreceptor capacity may be linked with adaptation to their extreme, shallow water environments.

### The origins of opsin diversification in the Cnidaria

The last common ancestor of the cnidaria and bilateria possessed three classes of distinct opsins giving rise to the cnidopsins, ASO-Is and ASO-IIs, yet the Medusozoa only retained the cnidopsins. The extant Anthozoa on the other hand possess multiple ASO-Is and substantially expanded and diversified the ASO-IIs. The ASO-I is the most ancient opsin class in the metazoan lineage and represents phylogenetically and, based on protein sequence, a coherent group, but to date its function is entirely unknown. Interestingly, the ASO-IIs are much more diverse at the sequence level and share a common ancestry with the tetraopsins and r-opsins as well as to the cnidopsins, xenopsins, ctenopsins and the well-studied c-opsins responsible for visual photoreception in vertebrates. This suggests that an urmetazoan animal possessed an ancestral but now extinct opsin that early in opsin evolution gave rise to multiple opsin classes. Frequent lineage-specific gains and losses then shaped the broad repertoire of light-sensing mechanisms and opsin classes that we see to date [12, 42, 67]. This capacity for diversity is still reflected by the novel ASO-II sub-cluster that is restricted to anthozoan anemones. Interestingly, if cnidopsins are indeed early xenopsins as suggested by our study and also Ramirez et al. (2016) and if ASO-IIs are indeed sister to the c-opsins as suggested here and by Ramirez et al. (2016) and Vöcking et al. (2017) the cnidarians may be the only animals where xenopsins and c-opsins (or at least their direct ancestral cousins) co-occur.

In accordance with the notion that ASO-IIs play an important role in the adaptation of Anthozoa, to their environments, we find that the highly conserved E181 amino acid residue is absent in the ASO-IIs suggesting that this key position has been modified, possibly to shift the ASO-IIs wavelength specificity more towards blue light (shorter wavelengths) absorption which may be a specific adaptation to aquatic marine environments where the penetration of longer wavelengths is reduced with increasing water depth [68, 69]. Indeed, by using computational modelling based on a vertebrate opsin crystal structure, it was shown that loss of E181 causes a light absorption shift of more than 100 nm towards blue light [70]. In the future, a functional analysis of the wavelength specificities of anthozoan opsins will provide fundamental new insight into the ability of basal animals to exploit various light cues.

### Adaptations to light-induced sexual reproduction

Coral sexual reproduction is a vital process for species viability. It is key for genetic diversity, dispersal by motile larvae and affects the abundance of juvenile corals to replenish aging coral communities. However, for sessile Anthozoa such as reef-building corals, achieving efficient fertilization rates is challenging because it occurs effectively only a few hours after gamete release by the parental colonies and gametes are easily diluted within the open space of the ocean. Accordingly, corals have evolved a precise spawning synchrony within populations integrating various environmental cues including temperature and solar irradiance to set the exact month, lunar cycles to set the exact night and circadian light cues to set the exact hour (see [71] and references therein). In this context it is worth noting that *Aiptasia* recapitulates key aspects of light-induced synchronized spawning observed in corals. Under laboratory conditions, delivery of LED-based blue-light for 5 consecutive nights, simulating full moon, triggers gamete release peaking 9-10 days after the last exposure to blue light. This effect is wavelength-specific as it works effectively with light at 400-460nm, while white light is ineffective [40]. Gamete release occurs ~5.5h after ‘sunset’ and after release, fertilization efficiency drops from 100% to ~25% within the first hour [72] underlining the need for precise timing of sexual reproduction within the Anthozoa.

Opsins have been implicated as light-sensors to trigger spawning in corals [5, 6, 73]. Moreover, a functional relationship between a light-induced, opsin-dependent spawning mechanism has been shown in a medusozoan, the jellyfish *Clytia hemispherica*. This elegant study found that a gonadal opsin senses blue light to trigger spawning upon dark-light transitions [15]. Thus, it is tempting to speculate that *Aiptasia* ASO-II.3, which is also expressed at elevated levels in gonads (Figure 3C) may serve a similar function for synchronous gamete release in this anthozoan species. *Aiptasia* represents a tractable model to experimentally dissect the mechanisms of light-induced sexual reproduction in Anthozoa including the integration of lunar cycles as well circadian periodicities to trigger gamete maturation and synchronous release. A mechanistic understanding of how Anthozoa have adapted the timing of sexual reproduction to their environments, together with analysing how environmental changes affect the spawning synchronicity within the ecosystem, is key to direct future research and conservation efforts [71].

### Adaptations to increased UV-induced DNA damage

Our results have also revealed significantly more diversity of the CRY/PL flavoproteins in the Anthozoa compared with the Medusozoa. While both contain CRY-DASH, CPD-II PLs and (6-4) PLs, Anthozoa also have two extra CRY classes (AnthoCRYs and Anthozoan CRYIIs), which are absent from Medusozoa. Thus, strikingly, the Medusozoa completely lack CRY genes. The AnthoCRYs represent a phylogenetically well-supported but previously unidentified, anthozoan-specific CRY family, which resolves at the base of animal CRY-IIs and (6-4) PLs. A hallmark of all CRY/PL proteins analysed to date is the highly conserved structure of the PHR domain consisting of a N-terminal domain and a FAD domain. However, here we reveal that the anthozoan AnthoCRYs exhibit an extensive and unprecedented PHR region duplication. Tandemly repeated PHR regions have never been observed before in any other eukaryotic or prokaryotic species. While the structure and function of the PHR has been studied in great deal in the context of PLs revealing its light-dependant DNA-repair function, the role of the PHR region in the CRYs remains rather unclear [21]. Nevertheless, until this current report, only single PHR regions have been scrutinized in PL or CRY proteins. The identification of tandem duplication of the PHR region in Anthozoa provides some tantalizing clues as to the functional significance of this domain. It is tempting to speculate that AnthoCRYs, which are clearly distinct from vertebrate CRY-IIs and CRY-Is and sister also to (6-4) PLs, have not yet lost their light-dependant DNA repair activity and in fact have evolved in the Anthozoa to provide higher UV-damage repair capacities reflected by their domain duplications. Furthermore, the observation that both AnthoCRY.1 and AnthoCRY.2 exhibit light-inducible expression would also be consistent with these genes playing a key role in the response to intensive sunlight exposure, a threat that sessile corals likely face on a daily basis in their sunlit tropical habitats.

### Adaptations of the circadian clock during cnidarian evolution

Transcriptional control of gene expression in response to light serves as a central regulatory element within the circadian clock core mechanism. This enables regular adjustment of the phase of the circadian clock to match that of the environmental day-night cycle. In non-mammalian vertebrates D-boxes mediate light-inducible gene expression, alone or in combination with other enhancers such as the E-box [61–63]. Furthermore, in fish, D-boxes also activate transcription in response to oxidative stress and UV exposure [64]. Thus, in the majority of vertebrates, D-boxes coordinate the transcription of a set of genes that includes certain clock genes as well as genes involved in the repair of UV-damaged DNA [61, 64] to constitute a cellular response to sunlight exposure. This contrasts with the situation of mammals where D-boxes exclusively direct clock-controlled rhythms of gene expression [74]. Therefore, the discovery of an enrichment of proximally spaced E and D-box enhancer elements in the promoters of the light inducible AnthoCRY genes supports the view that the D-box plays an ancestral sunlight-responsive role. Furthermore, this may also predict a function for AnthoCRYs in the complex cellular response to the damaging effects of sunlight which may involve responses to visible and UV light as well as oxidative stress.

CRYs are key regulators of the circadian clock in animals. Together with the Period proteins they serve as negative regulators within the core transcription-translation feedback loop mechanism [30]. Based on previous studies of plant and animal circadian clocks, it can be predicted that clock function is of fundamental importance for many anthozoan species. For example, the adaptation of the host cell physiology to the daily cycles of photosynthetic activity of the dinoflagellate symbionts as well as the sessile lifestyle of Anthozoa is likely to rely heavily on this endogenous timing mechanism that anticipates the course of the day-night cycle. This may well account for the conservation of CRY function in the Anthozoa. The absence of CRY in Medusozoa suggests that in these species, circadian clocks may be based upon fundamentally different mechanisms. Interestingly, canonical circadian clock genes were previously reported to be absent in *Hydra* and possibly all Medusozoa [7, 73, 75, 76]. However, whether medusozoan species have evolved alternative mechanisms to control rhythmic behaviour and physiology, or whether they may have actually lost circadian clock function represents a fascinating topic for future investigation.

### The origins of photoreceptor diversification

One key difference between the Anthozoa and Medusozoa is that Medusozoa have a free-swimming medusa phase during their lifecycle, while Anthozoa do not. Instead, Anthozoa are typically sessile animals, which are only motile during larval stages (Figure 1A). Possibly connected with this fundamental difference is that the Anthozoa lack eyes. Thus, we speculate that the evolution of relatively sophisticated eye-like structures based on light-sensitive cilia expressing cnidopsins allows for the integration of various light cues simultaneously in Medusozoa. Instead, in the Anthozoa the repertoire of non-visual opsins expanded in order to perceive light in different photic environments and during distinct life stages. This expansion could allow fine-tuning of animal physiology and behaviour including gamete release and phototaxis as well as optimizing conditions for their photosynthetic symbionts. Another striking example of how the expansion and sequence diversification of opsins allows adaptation to specific environments has recently been elucidated in deep-sea fish. While classically all vertebrates rely on only a single rod opsin rhodopsin 1 (RH1) for obtaining visual information in dim light conditions, some deep-sea fish have independently expanded their single RH1 gene to generate multiple RH1-like opsins that are tuned to different wavelengths of light by modulating key functional residues [44].

Our demonstration that eighteen distinct opsins and eight distinct PLs and CRYs are expressed in *Aiptasia*, in either larval or adult stages, in symbiotic or aposymbiotic animals, in a tissue-specific manner and in some cases, in a light inducible manner suggests that symbiotic anthozoans possess a remarkable, functional diversity in their photoreception mechanisms. The augmented complexity of photoreceptors in the Anthozoa is likely due to gene loss in the Medusozoa as well as to continued gene expansion and diversification within the Anthozoa and is indicative of distinct light-sensing mechanisms associated with different lifestyles. It may well be that the increased photoreceptor diversity of Anthozoa including corals and sea anemones represents an essential adaptation to their predominantly sessile lifestyle. Ultimately, complex light sensing mechanisms may permit the integration of sun and moon light to regulate physiology and behaviour, and to facilitate adaptation to their challenging environments: shallow, highly sunlit, tropical oceans where food is scarce and there is an enhanced risk of UV-induced DNA damage (Figure 1B). However, to date, no anthozoan-specific photoreceptor has been functionally characterized. Here we have generated an essential framework to experimentally analyse this diverse repertoire of non-visual photoreceptors using *Aiptasia* as a tractable model. A functional characterization of photoreceptors to uncover the mechanisms of light-sensing of cnidarians will provide profound new insight into the basic principles whereby metazoans adapt to light-dominated environments and how distinct lifestyles shape their photoreceptor repertoires.

## Materials and Methods

### *Aiptasia* culture and spawning

*Aiptasia* stocks were cultured as described [40]. Briefly, animals were reared from pedal lacerates for at least 6 months. For the spawning experiments, animals with a pedal disc diameter of 1 cm were separated into individual, small-sized, food-grade translucent polycarbonate tanks (GN 1/4-100 cm height, #44 CW; Cambro, Huntington Beach, USA) filled with artificial seawater (ASW) (Coral Pro Salt; Red Sea Aquatics Ltd, Houston, USA or REEF PRO; Tropic Marin, Switzerland) at 31-34 ppt salinity at 26°C. They were fed with *Artemia salina* nauplius larvae 5 times a week during the entire experimental period. ASW was exchanged twice per week and the tanks were cleaned using cotton tipped swabs as required.

### Sampling regimes

#### Circadian rhythmicity of CRY/PL expression

Firstly, *Aiptasia* polyps were adapted for 4 days to constant darkness and then exposed to light from white fluorescent bulbs with an intensity of ~20-25 μmol m^−2^ s^−1^ of photosynthetically active radiation (PAR), as measured with an Apogee PAR quantum meter (MQ-200; Apogee, Logan, USA) for a period of 8 hours, sampling at 2-hour intervals. Additionally, *Aiptasia* polyps were exposed to light-dark (LD, 12 h:12 h) cycles and then transferred to constant darkness (DD), sampling at 2, 8, 14 and 20 hours (LD) and 26, 32, 38 and 44 hours (DD).

### Computational methods

#### Identification of CRY, PL and opsin photoreceptors and PAR/bZIP transcription factor homologs

Potential CRY, PL and opsin sequences were recovered from *Aiptasia* genomic data (NCBI Bioproject PRJNA261862; [54]) by searching for annotation keywords and BLAST search using specific query sets. For opsins we used bovine (P51490), *Acropora palmata* (L0ATA4), *Carybdea rastonii* (B6F0Y5), honeybee (B7X752) and *Clytia hemisphaerica* (A0A2I6SFS3) opsins. For PLs and CRYs queries we used a set off previously published well-defined CRY/PL proteins, which we expanded to include known sponge and anthozoan CRYs [58, 77]. A set of known vertebrate PAR/bZIP transcription factors (TFs) were used as query sets to identify *Aiptasia* PAR/bZIP homologues using BLAST searches. The longest ORFs from recovered putative opsin genes were translated and aligned with the query sequences using ClustalW (GONNET, goc = 3, gec = 1.8). For opsins, sequences, which did not contain lysine K296, which is essential and indicative of chromophore binding, were excluded. For *Aiptasia* genes, we noticed that some of the associated gene models contained Ns (resulting in Xs in their amino acid sequences). Thus, we verify existing gene models by mapping reads from published short read RNA-seq libraries (Adult-apo: SRR1648359, SRR1648361, SRR1648362; Adult-intermediate: SRR1648363, SRR1648365, SRR1648367, SRR1648368; Adult-sym: SRR1648369, SRR1648370, SRR1648371, SRR1648372; Larvae-apo: SRR1648373, SRR1648374; Larvae-sym: SRR1648375, SRR1648376) to these gene models using HiSAT2 (https://ccb.jhu.edu/software/hisat2/index.shtml) at standard settings [78]. Uniquely mapped reads were extracted using samtools 1.2 and sequences then manually curated prior to alignment and phylogenies (Supplementary Files 1-3).

#### Phylogenetic analyses

A custom in-house BLAST database comprising more than 70 eukaryotic genera including *Nematostalla vectensis, Pocillopora, Stylophora, Orbicella, Acropora millepora, A. digitifera, Exaiptasia pallida (Cnidaria, Anthozoa), Abylopsis tetragona, Aegina citrea, Agalma elegans, Alatina alata, Atolla vanhoeffeni, Aurelia aurita, Calvadosia cruxmelitensis, Cassiopea xamachana, Chironex fleckeri, Chrysaora fuscescens, Clytia hemisphaerica, Craseoa lathetica, Craspedacusta sowerbyi, Craterolophus convolvulus, Cyanea capillata, Ectopleura larynx, Haliclystus sanjuanensis, Hydractinia echinata, H. polyclina, Hydra oligactis, H. viridissima, H. vulgaris, Leucernaria quadricornis, Nanomia bijuga, Physalia physalis, Podocoryna carnea, Stomolophus meleagris, Tripedalia cystophora, Turritopsis sp SK-2016 (Cnidaria, Medusozoa), Aplysia californica (Mollusca), Amphimedon queenslandica (Porifera), Caenorhabditis elegans (Nematoda), Drosophila melanogaster (Athropoda), Homo sapiens, Mus musculus, Danio rerio, Xenopus laevis* (all *Vertebrata*), *Monosiga brevicollis (Choanoflagellate), Pleurobrachia bachei (Ctenophora), Saccoglossus kowalevskii (Hemichordata), Strongylocentrotus purpuratus (Echinodermata), Saccharomyces cerevisiae (Fungi), Toxoplasma gondii, Plasmodium falciparum, Perkinsus marinus, Tetrahymena thermophila* (all *Alveolata*) and *Trichoplax adherens (Placozoa)* were used to generate a CRY/PL dataset for phylogenetic analysis. The majority of cnidarian sequences were obtained from published transcriptomes [79], but manually curated and translated in KNIME using in-house workflows comprising EMBOSS *getorf*. All other sequences were obtained from NCBI. For opsin phylogenies existing datasets were modified replacing *Exaiptasia* sequences with our newly verified *Aiptasia* opsin set [15, 42]. Outgroups were defined according to Vöcking et al. (2017) comprising several GPCR receptor family members (melatonin, octopamine, serotonin and adrenergic receptors) and Trichoplax opsin-like sequences, which all belong to class α rhodopsin-like GPCRs. For the detailed analysis of the ASO-II subtypes, additional ASO-II candidates were identified from the same database used for CRY/PL phylogenies using the previously identified *Aiptasia* ASO-IIs as query. Longest ORFs from recovered putative *Aiptasia* PAR/bZIP genes were translated and aligned to query sequences comprising multiple metazoan PAR/bZIPs and a CEBP (CCAAT-enhancer-binding proteins) outgroup. In all cases sequences were aligned using ClustalW (GONNET, goc: 3, gec: 1.8). Automated trimming was performed using trimAI using standard parameters [80]. Unstable leaves were identified and excluded using phyutilities with “-tt 100” settings [81]. Best-fitting amino acid substitution models were determined using PROTTEST3 (-JTT -LG -DCMut -Dayhoff -WAG -G -I -F -AIC -BIC; https://github.com/ddarriba/prottest3; [82]) and iqTree’s ModelFinder (-m MF -msub nuclear -nt AUTO; [83]). Maximum-likelihood trees were generated using iqTree (opsins: -m LG+G - bb 10000 -bnni -nt AUTO -alrt 10000 -abayes; CRY/PLs: -m LG+R6 -bb 10000 -bnni -nt AUTO -alrt 10000 -abayes; [84]). Bayesian inference trees were calculated using MrBayes (lset rates=gamma ngammacat=5; prset brlenspr=unconstrained:gammadir(1.0,0.1,1.0,1.0) aamodelpr=fixed(lg);mcmc ngen=1100000 samplefreq=200 printfreq=1000 nchains=4 temp=0.2 savebrlens=yes; starttree=random;set seed=518; sumt burnin=500; sump burnin=500; [85]). For opsin and CRY/PL phylogenies support values of resulting ML and Baysian analyses were combined using Treegraph2.14.0-771 [86]. Resulting trees were finalized using FigTree 1.4.4 (http://tree.bio.ed.ac.uk/software/figtree/) and Adobe Illustrator CC 2018. All alignments, tree files and accession numbers are provided in nexus format (Supplementary File 4-6).

#### Domain structure analysis, sequence logos and duplicate domain confirmation

The domain structures of the different CRYs and opsins were predicted using InterProScan v5.44 in Geneious R10 (Biomatters). Sequence logos were generated using Weblogo [87]. Sequencing a region spanning exon 7 and 8, which encode the C-terminal end of the FAD-binding domain of the AnthoCRY.1 PHR region 1 and the start of the N-terminal sequence of the AnthoCRY.1 PHR region 2 confirmed the presence of the PHR region tandem duplication. Here, RNA (as cDNA) and gDNA were PCR amplified using exon-specific primers (Supplementary Table 1), cloned into pJET2.1 and then Sanger sequenced. Alignment of sequenced fragments confirmed that the genomic sequence contains an intron and that the genomic and transcript sequences of AnthoCRY.1 span two individual PHR regions and that both are expressed.

#### Conserved intron structure analysis

Reference sequences were chosen at random to represent the canonical exon-intron structure of the respective Opsin/CRY/PL types. Reference and *Aiptasia* opsin gene models were generated using WebScipio [88] and conserved introns were identified using GenePainter 2.0 [89].

#### Expression quantification of PLs, CRYs and opsins

We analysed expression of the CRY, PL and opsin genes using the same published short read RNA-seq libraries that we used for gene model verification comprising data for adult and larval life stages from aposymbiotic and symbiotic states including 2-4 biological replicates per sample treatment [90]. The ultra-fast, bias-aware short read mapper Salmon [91] and in-house R scripts were used to generate a TMM normalised expression quantitation matrix across all conditions and samples. Average expression data for adult and larval *Aiptaisa* was used irrespective of their symbiotic state to analyse the differential developmental expression of CRYs and opsins. To compare the effect of symbiosis, the average expression in symbiotic and aposymbiotic adults was compared. Graphs were drawn and significance levels were determined using a multiple t-test in Prism 8.1.1 (GraphPad).

#### D-box and E-box searches

Potential PAR/bZIP binding sites in the genomic region 1kb upstream of the CRY/PL TSS were identified using MATCH 1.0 Public (http://gene-regulation.com/pub/programs.html) (Binding sites for Hlf (TransFac ID: T01071) and VBP (TransFac ID: T00881) were considered D-Boxes). The identified potential sites were aligned to the canonical D-Box elements identified in zebrafish using ClustalW (GONNET, goc: 3, gec: 1.8). Statistically overrepresented E-box motifs were identified in the same genomic regions using Clover [92] and a library of 9 TF binding motifs including several know (MITF and USF TF binding motifs) and one manually generated E-box motif. An approx. 6Mbp *Aiptasia* genomic scaffold (Genbank Accession NW_018384103.1) was used as a background sequence.

### Gene expression

#### RNA extraction and qPCR

For circadian rhythmicity qPCR analysis, polyps were macerated in Trizol at a concentration of 500mg per ml, snap frozen in liquid nitrogen and stored at −80 °C. Total RNA was extracted as described, but replacing phenol-chloroform with Trizol [37]. For determination of opsin expression levels in mesentery and tentacle tissues, a number of adult male *Aiptasia* were anesthetized in 7% MgCl_2_ (w/v) in ASW (1:1) for 1 hour and then transferred into Methacarn fixative (6:3:1 methanol:chloroform:acetic acid). Following two Methacarn changes in the first hour the samples were then incubated for 48 hours at RT. Individual polyps were then transferred into PBS and dissected to separate tentacle and mesentery tissues. Total RNA was extracted from dissected tissue samples as described in Hambleton et al. (2020) replacing phenol-chloroform with Trizol. In all cases cDNA was synthesised with 1 μg of total RNA per sample using a ReadyScript cDNA Synthesis Mix (Sigma-Aldrich). Primers for qPCR were determined using NCBI Primer BLAST (standard settings optimised for 100 bp exon-spanning amplicons) or designed manually using the same exon spanning rules when NCBI gene models were not available (For qPCR primers see: Supplementary Table 1). All qPCRs were run on a BioSystems StepOne Real-Time PCR System (ThermoFisher) using a Luna Universal qPCR Master Mix (NEB) at the fast setting following the manufacturer’s instructions to determine dCT levels in triplicate. Genes encoding 40S Ribosomal Proteins S7 and L11 (RPS7 and RPL11) and actin were chosen as comparison/baseline genes. Primers were validated in triplicate by amplicon sequencing of qPCR products. Melt curves were generated after each run confirming only a single product per reaction. Amplification efficiencies of each primer pair were determined through dilution series. Results were analysed according to a standard protocol (https://matzlab.weebly.com/data--code.html) using in-house KNIME (www.knime.com) workflows comprising an R integration of the Bayesian analysis pipeline MCMC.qPCR [93].

## Acknowledgments

This work was supported by the Deutsche Forschungsgemeinschaft (DFG) (Emmy Noether Program Grant GU 1128/3-1 to A.G.); the H2020 European Research Council (ERC Consolidator Grant 724715 to A.G.) and the Helmholtz Association (BioInterfaces in Technology and Medicine (BIFTM) programme to N.S.F.).

---

**Supplementary Figure 1: Intron phase analysis of opsin genes and summary table showing conserved structural and functional opsin motifs in *Aiptasia***

(A) Intron phase analysis of all *Aiptasia* opsins showing that the type-specific introns in Actiniarian ASO-IIs and ASO-IIs are conserved not only by position but also by intron phase (red boxes). (B) Summary table showing conserved structural and functional opsin motifs in *Aiptasia* in comparison to bovine rhodopsin and *Xenopus* melanopsin. These include: (i) two conserved cysteine (C) residues at positions 110 and 187, which are involved in disulphide-bond formation; (ii) two conserved glutamate [E] at position 113 and 181, which act as negative counterion to the proton of the Schiff base and may also affect spectral tuning; (iii) a glutamate [E] at position 134 located within a conserved motif (134-136; ERY in rhodopsin) that provides a negative charge to stabilise the inactive opsin molecule; (iv) a conserved lysine [K] at position 296 that is covalently linked to the 11-cis retinal chromophore via a Schiff base; (v) a conserved NPxxYx motif (302-313), which in rhodopsin contains a NKQ motif (310-312) that assists in maintaining structural integrity upon photopigment activation. The approximate position of the transmembrane domains is also depicted; modified from [94].

**Supplementary Figure 2: Distance matrix**

Distance matrix reflecting an alignment of 577 animal opsins. Note that opsins are clustered by amino acid similarity and are not sorted based on phylogenetic distance only but overall sequence similarity. Cnidarian opsin paralog clusters are highlighted and their names colour coded.

**Supplementary Figure 3: Extended expression profiles of *Aiptasia* opsins in mesentery and tentacle tissue**

qPCR analysis of mesentery and tentacle tissue after Methacarn fixation of adult *Aiptasia* polyps. ASO-I.2, ASO-II.1, ASO-II.4, Cnidopsin.3 and ASO_II.7 are significantly upregulated in tentacle tissue. ASO-II.3 is significantly upregulated in mesenteries. Cnidopsin.4 expression was not detected; ** P ≤ 0.01; error bars are: SEM.

**Supplementary Figure 4: Intron phase analysis of PL and CRY genes and expression profiles of *Aiptasia* PLs and CRYs in larva and adults**

(A) Intron phase analysis of all *Aiptasia* PLs and CRYs showing that select introns in CPDs, CRY-DASHs, CRY-Is, AnthoCRYs, animal CRY-IIs including Anthozoan CRY-IIs, and (6-4) PLs are conserved not only by position but also by intron phase (red boxes). (B) Bar chart comparing the expression (TMM normalised reads) of *Aiptasia* PLs and CRYs in adults and larva. AnthoCRY.2 is significantly upregulated in adults; ** P ≤ 0.01.

**Supplementary Figure 5: *Aiptasia* PAR-bZIP TF phylogeny**

Maximum-likelihood tree generated with iqTree reveals that *Aiptasia* (*E. pallida* in the tree) possesses four PAR-bZIP TFs: three TFs basal to HLF-type PAR/bZIP TFs and one DBP TF. The alignments and tree file including accession numbers are provided in nexus format (Supplementary File 6).

## Notes

### Competing Interest Statement

The authors have declared no competing interest.

