## Supplementary Figure 1-5 for "Photoreceptor complexity accompanies adaptation to challenging marine environments in Anthozoa"

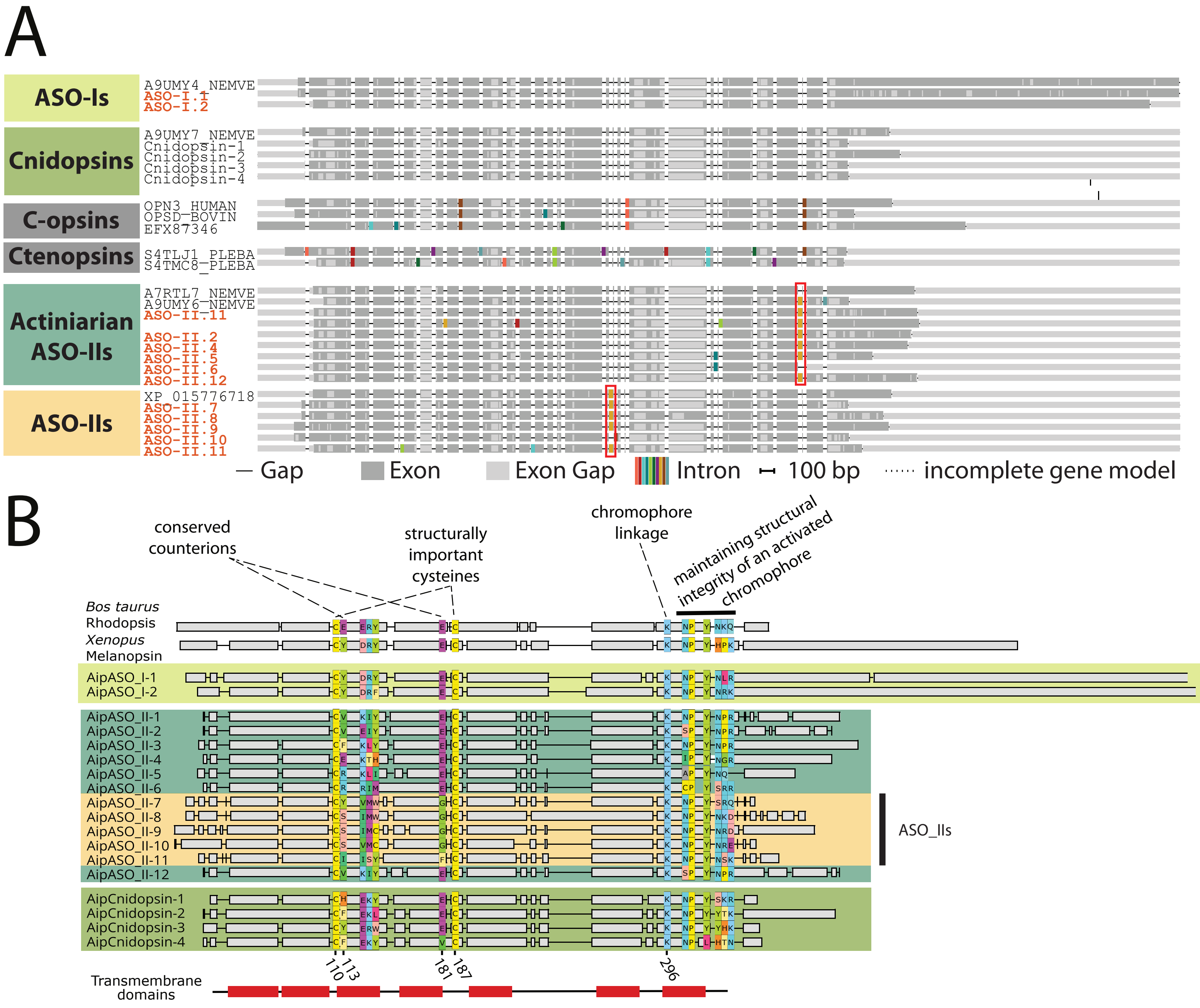

### Supplementary Figure 2

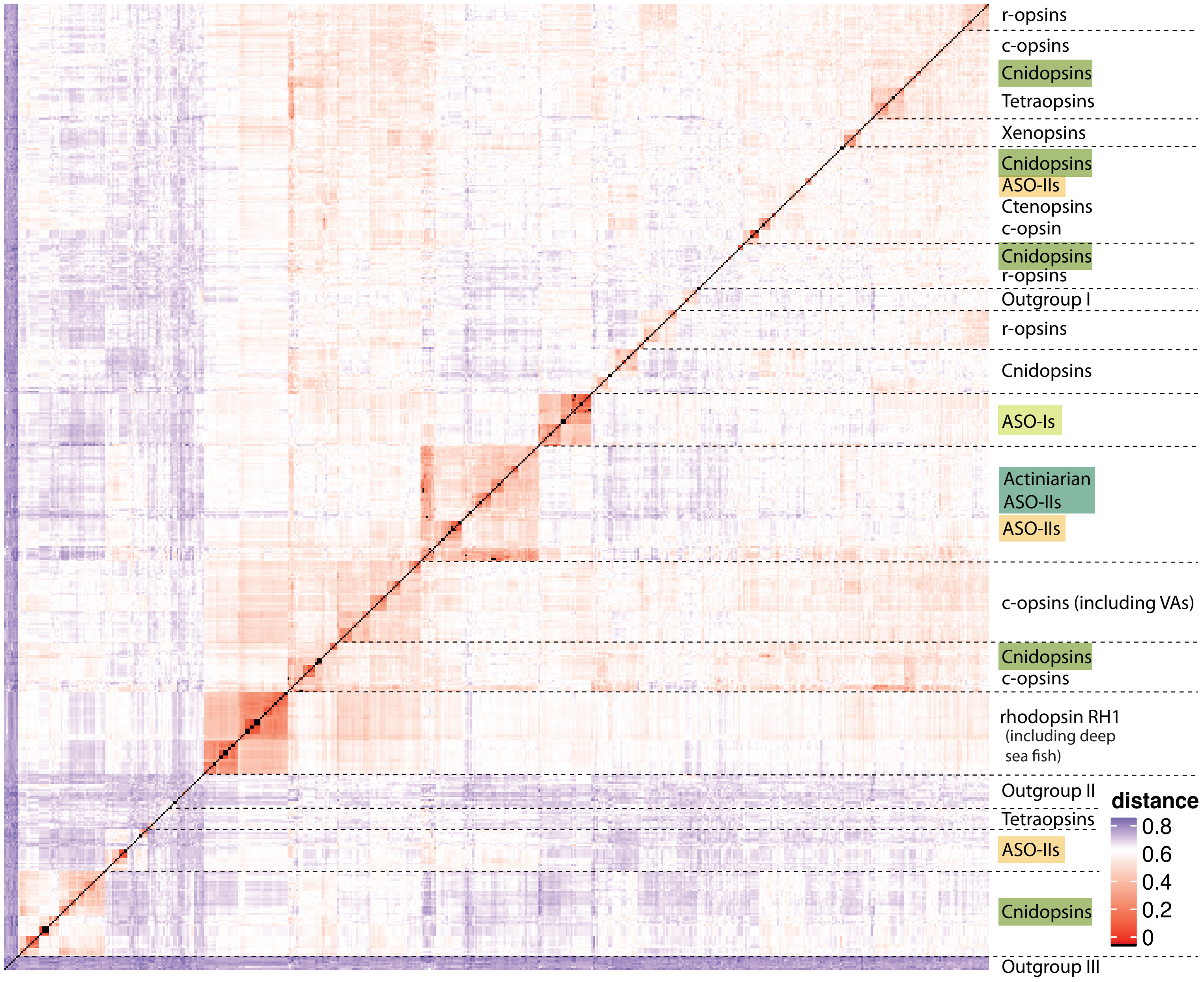

Supplementary Figure 3

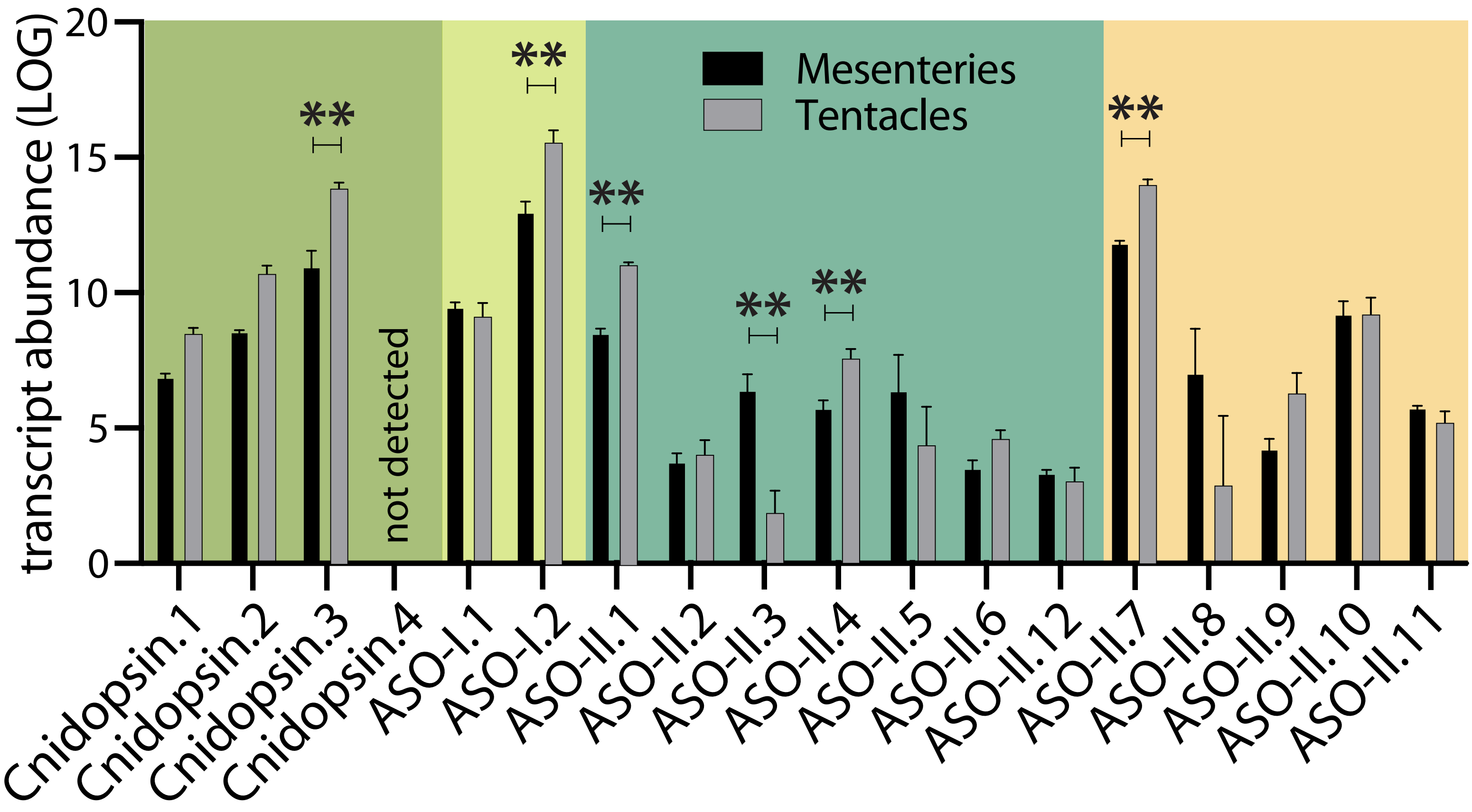

Supplementary Figure 4

A

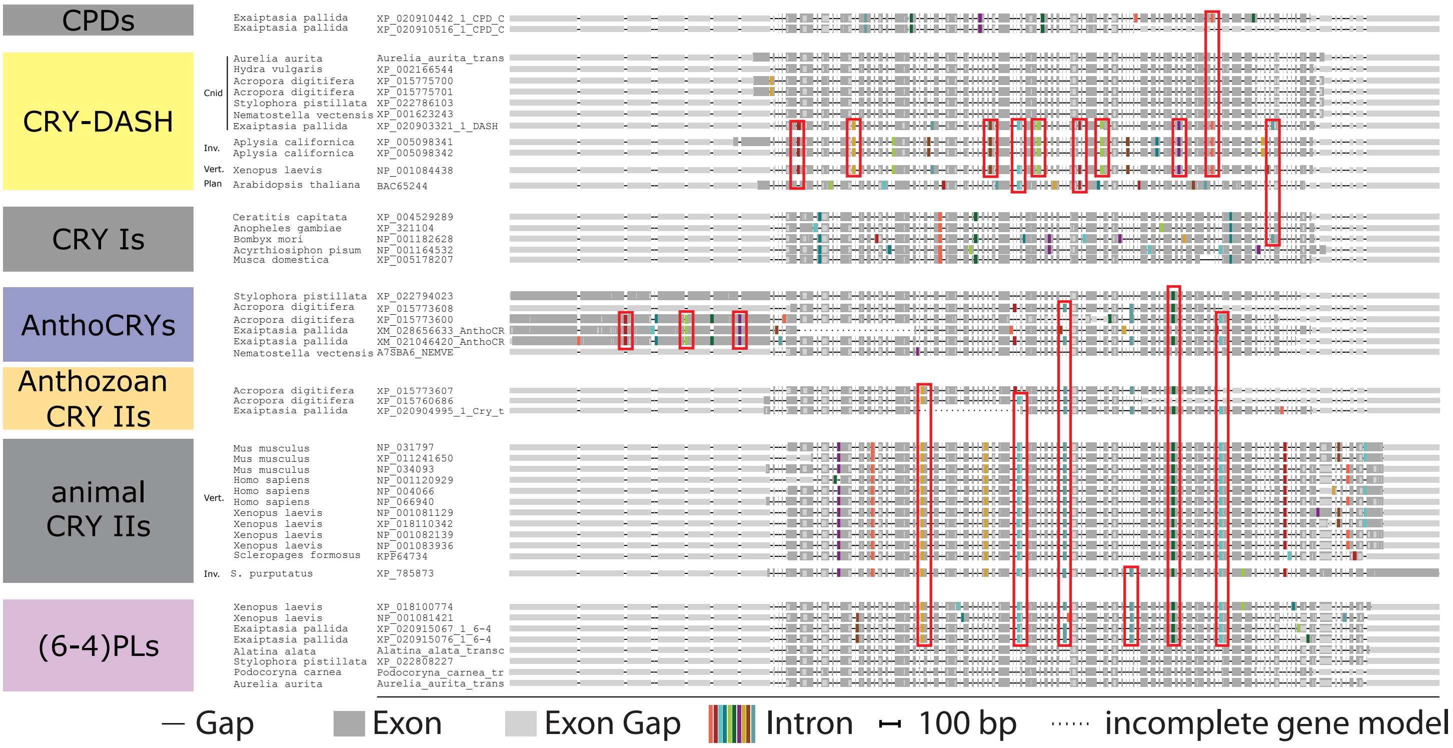


B

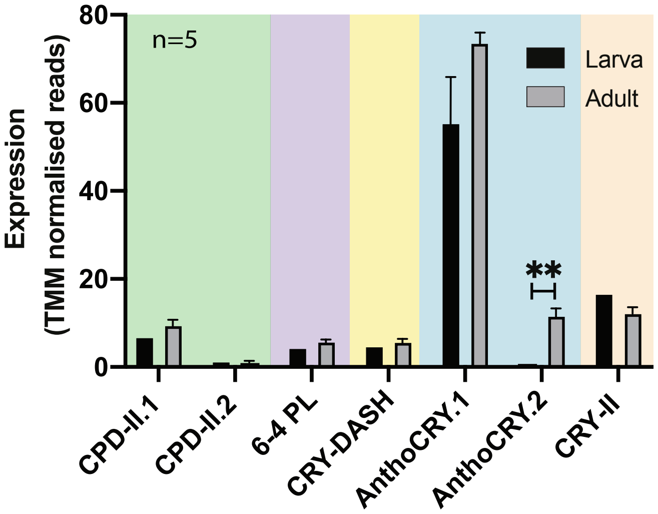


Note: all samples were taken at the mid-point of the 12-h light period

Supplementary Figure 5

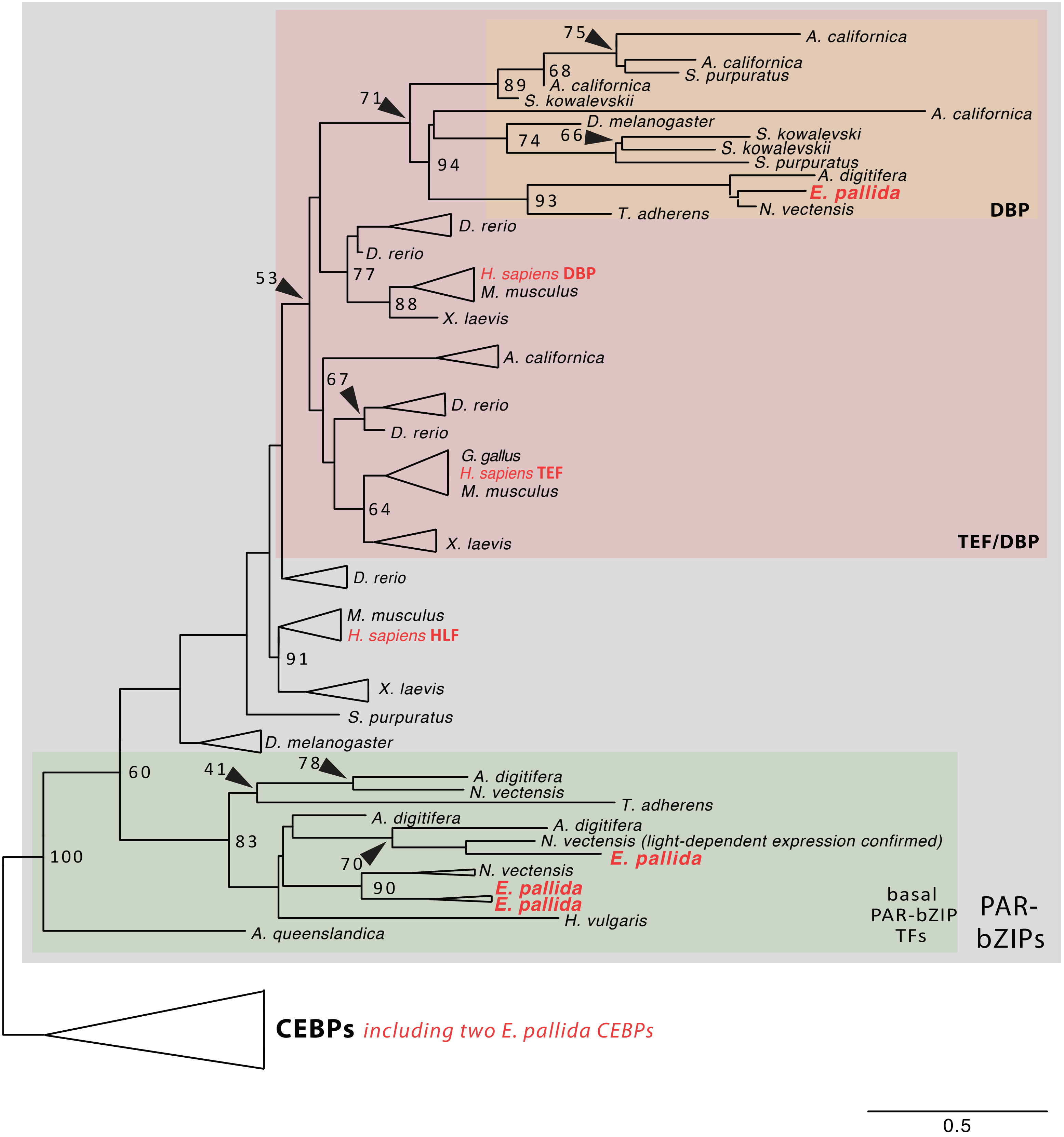

### Supplementary Figure legends

**Supplementary Figure 1: Intron phase analysis of opsin genes and summary table showing conserved structural and functional opsin motifs in *Aiptasia***

(A) Intron phase analysis of all *Aiptasia* opsins showing that the type-specific introns in Actiniarian ASO-IIs and ASO-IIs are conserved not only by position but also by intron phase (red boxes). (B) Summary table showing conserved structural and functional opsin motifs in *Aiptasia* in comparison to bovine rhodopsin and *Xenopus* melanopsin. These include: (i) two conserved cysteine (C) residues at positions 110 and 187, which are involved in disulphide-bond formation; (ii) two conserved glutamate [E] at position 113 and 181, which act as negative counterion to the proton of the Schiff base and may also affect spectral tuning; (iii) a glutamate [E] at position 134 located within a conserved motif (134–136; ERY in rhodopsin) that provides a negative charge to stabilise the inactive opsin molecule; (iv) a conserved lysine [K] at position 296 that is covalently linked to the 11-cis retinal chromophore via a Schiff base; (v) a conserved NPxxYx motif (302–313), which in rhodopsin contains a NKQ motif (310–312) that assists in maintaining structural integrity upon photopigment activation. The approximate position of the transmembrane domains is also depicted; modified from (Davies et al. 2010).

**Supplementary Figure 2: Distance matrix**

Distance matrix reflecting an alignment of 577 animal opsins. Note that opsins are clustered by amino acid similarity and are not sorted based on phylogenetic distance only but overall sequence similarity. Cnidarian opsin paralog clusters are highlighted and their names colour coded.

**Supplementary Figure 3: Extended expression profiles of *Aiptasia* opsins in mesentery and tentacle tissue**

qPCR analysis of mesentery and tentacle tissue after Methacarn fixation of adult *Aiptasia* polyps. ASO-I.2, ASO-II.1, ASO-II.4, Cnidopsin.3 and ASO_II.7 are significantly upregulated in tentacle tissue. ASO-II.3 is significantly upregulated in mesenteries. Cnidopsin.4 expression was not detected; ** $P \leq 0.01$; error bars are: SEM.

**Supplementary Figure 4: Intron phase analysis of PL and CRY genes and expression profiles of *Aiptasia* PLs and CRYs in larva and adults** (A) Intron phase analysis of all *Aiptasia* PLs and CRYs showing that select introns in CPDs, CRY-DASHs, CRY-Is, AnthoCRYs, animal CRY-IIs including Anthozoan CRY-IIs, and (6-4) PLs are conserved not only by position but also by intron phase (red boxes). (B) Bar chart comparing the expression

(TMM normalised reads) of *Aiptasia* PLs and CRYs in adults and larva. AnthoCRY.2 is significantly upregulated in adults; ** $P \leq 0.01$.

**Supplementary Figure 5: *Aiptasia* PAR-bZIP TF phylogeny**

Maximum-likelihood tree generated with iqTree reveals that *Aiptasia* (*E. pallida* in the tree) possesses four PAR-bZIP TFs: three TFs basal to HLF-type PAR/bZIP TFs and one DBP TF. The alignments and tree file including accession numbers are provided in nexus format (Supplementary File 6).
