## Supplementary File 6 for "Photoreceptor complexity accompanies adaptation to challenging marine environments in Anthozoa"

```
#NEXUS
begin taxa;
	dimensions ntax=137;
	taxlabels
	'S._kowalevskii_gi_585693090_ref_XP_002738815.2_'[&Created=M
on May 25 14:16:31 CEST 2020]
	'S._purpuratus_gi_72006198_ref_XP_787318.1_'[&Created=Mon
May 25 14:16:31 CEST 2020]
	'S._kowalevskii_gi_291227743_ref_XP_002733842.1_'[&Created=M
on May 25 14:16:31 CEST 2020]
	'T._adherens_lcl_XM_002109162.1_prot_XP_002109198.1_1'[&Crea
ted=Mon May 25 14:16:31 CEST 2020]
	'S._kowalevskii_gi_291222488_ref_XP_002731253.1_'[&Created=M
on May 25 14:16:31 CEST 2020]
	'S._purpuratus_gi_390352986_ref_XP_785519.3_'[&Created=Mon
May 25 14:16:31 CEST 2020]
	'A._californica_gi_524899363_ref_XP_005106114.1_'[&Created=M
on May 25 14:16:31 CEST 2020]
	'D._melanogaster_gi_24654082_ref_NP_611101.1_'[&Created=Mon
May 25 14:16:31 CEST 2020]
	'S._kowalevskii_gi_291234179_ref_XP_002737021.1_'[&Created=M
on May 25 14:16:31 CEST 2020]
	'N._vectensis_gi_156408121_ref_XP_001641705.1_'[&Created=Mon
May 25 14:16:31 CEST 2020]

thyrotroph_embryonic_factor_like_2__Exaiptasia_pallida_[&Description
="thyrotroph embryonic factor-like [Exaiptasia pallida]",Created=Mon
May 25 14:16:31 CEST 2020]
	'A._digitifera_lcl_XM_015908039.1_prot_XP_015763525.1_1'[&Cr
eated=Mon May 25 14:16:31 CEST 2020]
	'A._californica_gi_524915872_ref_XP_005112723.1_'[&Created=M
on May 25 14:16:31 CEST 2020]
	'A._californica_gi_524895551_ref_XP_005104258.1_'[&Created=M
on May 25 14:16:31 CEST 2020]
	'A._californica_gi_524895679_ref_XP_005104321.1_'[&Created=M
on May 25 14:16:31 CEST 2020]
	'A._californica_gi_524895681_ref_XP_005104322.1_'[&Created=M
on May 25 14:16:31 CEST 2020]
	'A._californica_gi_524895683_ref_XP_005104323.1_'[&Created=M
on May 25 14:16:31 CEST 2020]
	'D._rerio_lcl_NM_131400.1_prot_NP_571475.1_1'[&Created=Mon
May 25 14:16:31 CEST 2020]
	'D._rerio_lcl_XM_005156135.3_prot_XP_005156192.1_1'[&Created
=Mon May 25 14:16:31 CEST 2020]
	'D._rerio_lcl_XM_005156136.3_prot_XP_005156193.1_1'[&Created
=Mon May 25 14:16:31 CEST 2020]
	'D._rerio_lcl_NM_001020661.1_prot_NP_001018497.1_1'[&Created
=Mon May 25 14:16:31 CEST 2020]
	'M._musculus_gi_226958409_ref_NP_705617.2_'[&Created=Mon May
25 14:16:31 CEST 2020]
	'M._musculus_gi_568991979_ref_XP_006520806.1_'[&Created=Mon
May 25 14:16:31 CEST 2020]
	'H._sapiens_gi_223972670_ref_NP_001138870.1_'[&Created=Mon
May 25 14:16:31 CEST 2020]
```

'M._musculus_gi_8394435_ref_NP_059072.1_'[&Created=Mon May 25 14:16:31 CEST 2020]
'Thyrotroph_embryonic_factor_OS_Homo_sapiens_OX_9606_GN_TEF_PE_1_SV_3'[&Created=Mon May 25 14:16:31 CEST 2020]
TEF_CHICK[&Description="RecName: Full=Transcription factor VBP; AltName: Full=Thyrotroph embryonic factor homolog; AltName: Full=Vitellogenin gene-binding protein;",Created=Mon May 25 14:16:31 CEST 2020]
'X._laevis_lcl_NM_001086779.1_prot_NP_001080248.1_1'[&Created=Mon May 25 14:16:31 CEST 2020]
'X._laevis_lcl_XM_018256478.1_prot_XP_018111967.1_1'[&Created=Mon May 25 14:16:31 CEST 2020]
'X._laevis_lcl_XM_018256479.1_prot_XP_018111968.1_1'[&Created=Mon May 25 14:16:31 CEST 2020]
'X._laevis_lcl_NM_001094595.1_prot_NP_001088064.1_1'[&Created=Mon May 25 14:16:31 CEST 2020]
'X._laevis_lcl_XM_018259674.1_prot_XP_018115163.1_1'[&Created=Mon May 25 14:16:31 CEST 2020]
'X._laevis_lcl_XM_018259675.1_prot_XP_018115164.1_1'[&Created=Mon May 25 14:16:31 CEST 2020]
'X._laevis_lcl_XM_018259676.1_prot_XP_018115165.1_1'[&Created=Mon May 25 14:16:31 CEST 2020]
'H._sapiens_gi_530412024_ref_XP_005257326.1_'[&Created=Mon May 25 14:16:31 CEST 2020]
HLF_HUMAN[&Description="RecName: Full=Hepatic leukemia factor;",Created=Mon May 25 14:16:31 CEST 2020]
'M._musculus_gi_31982951_ref_NP_766151.1_'[&Created=Mon May 25 14:16:31 CEST 2020]
'M._musculus_gi_568973277_ref_XP_006533067.1_'[&Created=Mon May 25 14:16:31 CEST 2020]
'M._musculus_gi_568973279_ref_XP_006533068.1_'[&Created=Mon May 25 14:16:31 CEST 2020]
'M._musculus_gi_568973283_ref_XP_006533070.1_'[&Created=Mon May 25 14:16:31 CEST 2020]
'X._laevis_lcl_XM_018235125.1_prot_XP_018090614.1_1'[&Created=Mon May 25 14:16:31 CEST 2020]
'X._laevis_lcl_XM_018235126.1_prot_XP_018090615.1_1'[&Created=Mon May 25 14:16:31 CEST 2020]
'X._laevis_lcl_XM_018240540.1_prot_XP_018096029.1_1'[&Created=Mon May 25 14:16:31 CEST 2020]
'X._laevis_lcl_XM_018240541.1_prot_XP_018096030.1_1'[&Created=Mon May 25 14:16:31 CEST 2020]
'D._rerio_lcl_NM_001077334.2_prot_NP_001070802.1_1'[&Created=Mon May 25 14:16:31 CEST 2020]
'D._rerio_lcl_NM_001197059.1_prot_NP_001183988.1_1'[&Created=Mon May 25 14:16:31 CEST 2020]
'D._rerio_lcl_XM_005156334.4_prot_XP_005156391.1_1'[&Created=Mon May 25 14:16:31 CEST 2020]
'D._rerio_lcl_NM_001197062.1_prot_NP_001183991.1_1'[&Created=Mon May 25 14:16:31 CEST 2020]
'D._rerio_lcl_XM_005157897.4_prot_XP_005157954.1_1'[&Created=Mon May 25 14:16:31 CEST 2020]
'D._rerio_lcl_NM_001197060.1_prot_NP_001183989.1_1'[&Created=Mon May 25 14:16:31 CEST 2020]

'M._musculus_gi_170650717_ref_NP_058670.2_'[&Created=Mon May 25 14:16:31 CEST 2020]
'M._musculus_gi_568947265_ref_XP_006540659.1_'[&Created=Mon May 25 14:16:31 CEST 2020]
DBP_HUMAN[&Description="RecName: Full=D site-binding protein; AltName: Full=Albumin D box-binding protein; AltName: Full=Albumin D-element-binding protein; AltName: Full=Tax-responsive enhancer element-binding protein 302; Short=TaxREB302;",Created=Mon May 25 14:16:31 CEST 2020]
'M._musculus_gi_568947267_ref_XP_006540660.1_'[&Created=Mon May 25 14:16:31 CEST 2020]
'X._laevis_lcl_XM_018226265.1_prot_XP_018081754.1_1'[&Created=Mon May 25 14:16:31 CEST 2020]
'X._laevis_lcl_XM_018228082.1_prot_XP_018083571.1_1'[&Created=Mon May 25 14:16:31 CEST 2020]
'S._purpuratus_gi_193788685_ref_NP_001123286.1_'[&Created=Mon May 25 14:16:31 CEST 2020]
'D._melanogaster_gi_24660427_ref_NP_729297.1_'[&Created=Mon May 25 14:16:31 CEST 2020]
'D._melanogaster_gi_442630879_ref_NP_001261547.1_'[&Created=Mon May 25 14:16:31 CEST 2020]
'D._melanogaster_gi_24660458_ref_NP_729301.1_'[&Created=Mon May 25 14:16:31 CEST 2020]
'D._melanogaster_gi_281365802_ref_NP_729302.2_'[&Created=Mon May 25 14:16:31 CEST 2020]
'D._melanogaster_gi_24660450_ref_NP_729300.1_'[&Created=Mon May 25 14:16:31 CEST 2020]
'D._melanogaster_gi_281365800_ref_NP_729298.2_'[&Created=Mon May 25 14:16:31 CEST 2020]
'D._melanogaster_gi_24660438_ref_NP_729299.1_'[&Created=Mon May 25 14:16:31 CEST 2020]
'D._melanogaster_gi_442630877_ref_NP_001261546.1_'[&Created=Mon May 25 14:16:31 CEST 2020]
'D._melanogaster_gi_442630875_ref_NP_001261545.1_'[&Created=Mon May 25 14:16:31 CEST 2020]
'D._melanogaster_gi_24660469_ref_NP_729303.1_'[&Created=Mon May 25 14:16:31 CEST 2020]
'D._melanogaster_gi_442630873_ref_NP_001261544.1_'[&Created=Mon May 25 14:16:31 CEST 2020]
'N._vectensis_gi_156336982_ref_XP_001619763.1_'[&Created=Mon May 25 14:16:31 CEST 2020]
'N._vectensis_gi_156400985_ref_XP_001639072.1_'[&Created=Mon May 25 14:16:31 CEST 2020]
'A._digitifera_lcl_XM_015892501.1_prot_XP_015747987.1_1'[&Created=Mon May 25 14:16:31 CEST 2020]

thyrotroph_embryonic_factor_like_1__Exaiptasia_pallida_[&Description="thyrotroph embryonic factor-like [Exaiptasia pallida]",Created=Mon May 25 14:16:31 CEST 2020]
Transcription_factor_VBP__Exaiptasia_pallida_[&Created=Mon May 25 14:16:31 CEST 2020]
'H._vulgaris_gi_221106316_ref_XP_002167159.1_'[&Created=Mon May 25 14:16:31 CEST 2020]
'S._kowalevskii_gi_585674208_ref_XP_002736397.2_'[&Created=M

```
on May 25 14:16:31 CEST 2020]
        'A._digitifera_lcl_XM_015892500.1_prot_XP_015747986.1_1'[&Cr
eated=Mon May 25 14:16:31 CEST 2020]
        'N._vectensis_gi_156408099_ref_XP_001641694.1_'[&Created=Mon
May 25 14:16:31 CEST 2020]
        'A._queenslandica_gi_340372715_ref_XP_003384889.1_'[&Created
=Mon May 25 14:16:31 CEST 2020]

hepatic_leukemia_factor_like__Exaiptasia_pallida_[&Created=Mon May
25 14:16:31 CEST 2020]
        'N._vectensis_gi_156388041_ref_XP_001634510.1_'[&Created=Mon
May 25 14:16:31 CEST 2020]
        'A._digitifera_lcl_XM_015913431.1_prot_XP_015768917.1_1'[&Cr
eated=Mon May 25 14:16:31 CEST 2020]
        'T._adherens_lcl_XM_002117855.1_prot_XP_002117891.1_1'[&Crea
ted=Mon May 25 14:16:31 CEST 2020]
        'A._californica_gi_524907652_ref_XP_005108942.1_'[&Created=M
on May 25 14:16:31 CEST 2020]
        'H._vulgaris_gi_221116695_ref_XP_002159994.1_'[&Created=Mon
May 25 14:16:31 CEST 2020]
        A0A1D5NSQ7_DANRE[&Description="SubName: Full=CCAAT/enhancer-
binding protein (C/EBP) 1 {ECO:0000313|
Ensembl:ENSDARP00000142714};",Created=Mon May 25 14:16:31 CEST 2020]
        'D._rerio_lcl_NM_131837.1_prot_NP_571912.1_1'[&Created=Mon
May 25 14:16:31 CEST 2020]
        'CCAAT_enhancer_binding_protein_delta_OS_Homo_sapiens_OX_960
6_GN_CEBPD_PE_1_SV_2'[&Created=Mon May 25 14:16:31 CEST 2020]
        'M._musculus_gi_110347410_ref_NP_031705.3_'[&Created=Mon May
25 14:16:31 CEST 2020]
        'X._laevis_lcl_NM_001089607.2_prot_NP_001083076.1_1'[&Create
d=Mon May 25 14:16:31 CEST 2020]
        'X._laevis_lcl_NM_001089609.1_prot_NP_001083078.1_1'[&Create
d=Mon May 25 14:16:31 CEST 2020]
        'D._rerio_lcl_NM_131887.1_prot_NP_571962.1_1'[&Created=Mon
May 25 14:16:31 CEST 2020]
        'CCAAT_enhancer_binding_protein_epsilon_OS_Homo_sapiens_OX_9
606_GN_CEBPE_PE_1_SV_2'[&Created=Mon May 25 14:16:31 CEST 2020]
        'M._musculus_gi_46369479_ref_NP_997014.1_'[&Created=Mon May
25 14:16:31 CEST 2020]
        'X._laevis_lcl_XM_018243908.1_prot_XP_018099397.1_1'[&Create
d=Mon May 25 14:16:31 CEST 2020]
        'X._laevis_lcl_XM_018259384.1_prot_XP_018114873.1_1'[&Create
d=Mon May 25 14:16:31 CEST 2020]
        'X._laevis_lcl_NM_001086806.1_prot_NP_001080275.1_1'[&Create
d=Mon May 25 14:16:31 CEST 2020]
        'X._laevis_lcl_NM_001091687.1_prot_NP_001085156.1_1'[&Create
d=Mon May 25 14:16:31 CEST 2020]
        'D._rerio_lcl_NM_131885.2_prot_NP_571960.1_1'[&Created=Mon
May 25 14:16:31 CEST 2020]
        'H._sapiens_gi_28872794_ref_NP_004355.2_'[&Created=Mon May
25 14:16:31 CEST 2020]
        'H._sapiens_gi_566559994_ref_NP_001274364.1_'[&Created=Mon
May 25 14:16:31 CEST 2020]
        'H._sapiens_gi_566559992_ref_NP_001274353.1_'[&Created=Mon
```

```
May 25 14:16:31 CEST 2020]
        'H._sapiens_gi_551894998_ref_NP_001272758.1_'[&Created=Mon
May 25 14:16:31 CEST 2020]
        'M._musculus_gi_566559971_ref_NP_001274444.1_'[&Created=Mon
May 25 14:16:31 CEST 2020]
        'M._musculus_gi_566559986_ref_NP_001274450.1_'[&Created=Mon
May 25 14:16:31 CEST 2020]
        'M._musculus_gi_86198301_ref_NP_031704.2_'[&Created=Mon May
25 14:16:31 CEST 2020]
        'M._musculus_gi_566559969_ref_NP_001274443.1_'[&Created=Mon
May 25 14:16:31 CEST 2020]
        'M._musculus_gi_567316233_ref_NP_001274452.1_'[&Created=Mon
May 25 14:16:31 CEST 2020]
        'M._musculus_gi_6753404_ref_NP_034013.1_'[&Created=Mon May
25 14:16:31 CEST 2020]
        'M._musculus_gi_567757569_ref_NP_001274667.1_'[&Created=Mon
May 25 14:16:31 CEST 2020]
        'M._musculus_gi_567757572_ref_NP_001274668.1_'[&Created=Mon
May 25 14:16:31 CEST 2020]
        'H._sapiens_gi_551895123_ref_NP_001272808.1_'[&Created=Mon
May 25 14:16:31 CEST 2020]
        'X._laevis_lcl_NM_001095915.1_prot_NP_001089384.1_1'[&Create
d=Mon May 25 14:16:31 CEST 2020]
        'X._laevis_lcl_NM_001172167.1_prot_NP_001165638.1_1'[&Create
d=Mon May 25 14:16:31 CEST 2020]
        'D._rerio_lcl_NM_131884.2_prot_NP_571959.2_1'[&Created=Mon
May 25 14:16:31 CEST 2020]
        'D._melanogaster_gi_665403432_ref_NP_001286836.1_'[&Created=
Mon May 25 14:16:31 CEST 2020]
        'N._vectensis_gi_156384801_ref_XP_001633321.1_'[&Created=Mon
May 25 14:16:31 CEST 2020]

uncharacterized_protein_LOC110247154__Exaiptasia_pallida_[&Created=M
on May 25 14:16:31 CEST 2020]
        'A._digitifera_lcl_XM_015902730.1_prot_XP_015758216.1_1'[&Cr
eated=Mon May 25 14:16:31 CEST 2020]
        'H._vulgaris_gi_449684526_ref_XP_002164910.2_'[&Created=Mon
May 25 14:16:31 CEST 2020]
        'H._vulgaris_gi_449684528_ref_XP_002164958.2_'[&Created=Mon
May 25 14:16:31 CEST 2020]
        'T._adherens_lcl_XM_002113628.1_prot_XP_002113664.1_1'[&Crea
ted=Mon May 25 14:16:31 CEST 2020]
        'H._vulgaris_gi_449679187_ref_XP_004209260.1_'[&Created=Mon
May 25 14:16:31 CEST 2020]
        'N._vectensis_gi_156375819_ref_XP_001630276.1_'[&Created=Mon
May 25 14:16:31 CEST 2020]
        'A._digitifera_lcl_XM_015919129.1_prot_XP_015774615.1_1'[&Cr
eated=Mon May 25 14:16:31 CEST 2020]
        'CCAAT_enhancer_binding_protein_gamma_like__Exaiptasia_palli
da_'[&Created=Mon May 25 14:16:31 CEST 2020]
        'A._californica_gi_325120973_ref_NP_001191392.1_'[&Created=M
on May 25 14:16:31 CEST 2020]
        'CCAAT_enhancer_binding_protein_gamma_OS_Homo_sapiens_OX_960
6_GN_CEBPG_PE_1_SV_1'[&Created=Mon May 25 14:16:31 CEST 2020]
```

```
        'M._musculus_gi_61966683_ref_NP_034014.1_'[&Created=Mon May
25 14:16:31 CEST 2020]
        'D._rerio_lcl_NM_131886.1_prot_NP_571961.1_1'[&Created=Mon
May 25 14:16:31 CEST 2020]
        'X._laevis_lcl_NM_001095901.1_prot_NP_001089370.1_1'[&Create
d=Mon May 25 14:16:31 CEST 2020]
        'X._laevis_lcl_XM_018261138.1_prot_XP_018116627.1_1'[&Create
d=Mon May 25 14:16:31 CEST 2020]
        'S._purpuratus_gi_390355599_ref_XP_003728584.1_'[&Created=Mo
n May 25 14:16:31 CEST 2020]
        'S._kowalevskii_gi_585693956_ref_XP_006821724.1_'[&Created=M
on May 25 14:16:31 CEST 2020]
        'A._californica_gi_524898017_ref_XP_005105463.1_'[&Created=M
on May 25 14:16:31 CEST 2020]
        'S._kowalevskii_gi_585693879_ref_XP_006821714.1_'[&Created=M
on May 25 14:16:31 CEST 2020]
        'S._purpuratus_gi_390340887_ref_XP_003725328.1_'[&Created=Mo
n May 25 14:16:31 CEST 2020]
        'A._queenslandica_gi_340367947_ref_XP_003382514.1_'[&Created
=Mon May 25 14:16:31 CEST 2020]
;
end;

begin characters;
        dimensions nchar=1022;
        format datatype=protein missing=? gap=-;
        matrix
        'S._kowalevskii_gi_585693090_ref_XP_002738815.2_'       ----
--------------------------------------------------------------------
--------------------------------------------------------------------
--------------------------------------------------------------------
--------------------------------------------------------------------
----------MSEEK-LYDFQQQFEE----D--KQKIPPISTFAKASAY--DGRYPIAAQNT---
NRAFY-LVS-----------GE----GNPS----------------------------FVY--P----
P-------TSSS--TGNLQP-------PLPANMV--GQS-------SMM----------
MYHRGGTD----VSTS--------G----TLP-PT-P-----------
PVSADEKDHNID---------DP--N--AR-------DS---GLIRMLS--Q---------
TKQ-------MEL-P-----SHL--PHPPHLM--------------------MT----------
APQ----------VHLHTHTHAP--GPIS---PMDNES-------------------------F-DG-
EGDDPFDVMSHVRMNH--SKR-RST----KKAVTD-----------DMK-
DHRYFERRKKNNMAAKRSRDARK-YREEQISHRASXLERENAHLRSQLETLKEE---
AYSLRQLLAQ----KS--
NSS-----------------------------------------------------------------
--------------------------------------------------------------------
----------------------------------------------------
        'S._purpuratus_gi_72006198_ref_XP_787318.1_'    ------------
--------------------------------------------------------------------
--------------------------------------------------------------------
--------------------------------------------------------------------
--------------------------------------------------------------------
--MSEEEHIYEFNREYEEKSSTDS-KRSIPPISTFTKGTGNGTEG-
YPTTLYNSKDDNQSMANLESNNNDSSNTDSSSESMSVASPSQQQKLSQPQETQQQPTQHQQLLQQQQQ
PHAYNTPAAGFPYSNQVLVTSSSQETNGVQASHIIHSGAIPATMVPAGMSLQGHTVIPFMTSTSAPSQ
AGIIQAGGPDQMMQYSTSAMHLLTSAGQPVYSMPAPVQPNSSVIYVNDSIPVQPTVLPHNIDLQQHLQ
```

```
HQHQPPKNGSAEEKNDQDDDSRKISLATLVSNAQNMQPLVQAGTNQRPGLPHLMQLNPPAQSMSSLGG
PIPQHFIPGPGPIPI---------QGSIATGNYGNAQVLPPQQQQEVPTS-
VVCMPPHIGPPPHGMVEQPNPMIGAGPGPGGIG------------------
MPDGVEGNQQGPGSSAVRRN---SKR-RGN----KRDVPV-----------DEK-
DNRYYERRDRNNKAAKRSRDARK-IREEQVGMRAHYLEKENDFLRAQLNTLREE---
ANSLRLLLAQ----
RPPVNQMQPTGQPPMQ----------------------------------------------------
--------------------------------------------------------------------
--------------------------------------------------------
        'S._kowalevskii_gi_291227743_ref_XP_002733842.1_'        ----
--------------------------------------------------------------------
--------------------------------------------------------------------
--------------------------------------------------------------------
--------------------------------------------------------------------
-----MLKLEVKEDSVISPMSKWPKDIGQKDKSGKELPKSENITIPAIN--QN-MKQNTENT----
HELGTLMQQDN--STPQLGGE----KTPQ----------------------------
QDNKIP------------SRISRRKGKVPPRKKLKT-DIPEESS-
DDDSECGRLAIHLKED--------MNHKESTIIKQEPMIQDER---CYG----
IMPTSSLTNG---------PNYRSVQSEKTYTN------ELPDKEYSVETN-----
QPQTTRLQTTYP-LQN-----GLITPN------LVPVNP-----AIL--
QDGRQVLIAALPGNV---------SGIAAAGS-------IPY---------
ISSVPQVEGGNPLNLIAATAALASRVH------------------------
VVDSPQSADDSISNEDIDENSCLSKKERAE----KQPVPD-----------EQK-
DHRYFERRRRNNAAAKRSRDARR-LREEEMSLRISQLEHENSFLRSQLAAMREK---
AESLRKNLVV----
DDDVNSTTT-----------------------------------------------------------
--------------------------------------------------------------------
--------------------------------------------------------
        'T._adherens_lcl_XM_002109162.1_prot_XP_002109198.1_1'  ----
--------------------------------------------------------------------
--------------------------------------------------------------------
--------------------------------------------------------------------
--------------------------------------------------------------------
--------------------------------------------------------------------
--------------------------------------------------------------------
--------------------------------------------------------------------
--------------------------------------------------------------------
-----------MTDRNQ--QN------CLSCS--------TLSP----EYCF--
MHRMDVM--------------------------------SQK---------
CLSINPLHNGQIYPTLKS--NLYPTLP------------------------SHEG-
NMDNVRIVVGALDKKSANLSASKKS----KTPVPN-----------DRK-
DDKYWDKRKRNNESAKRSREARK-LKDNQVASRATWLEEENVKIKAENAALREE---
LACWRYYFSQP--
FSAQSNPKPISGIPLKPYLF------------------------------------------------
--------------------------------------------------------------------
---------------------------------------------------------
        'S._kowalevskii_gi_291222488_ref_XP_002731253.1_'        ----
--------------------------------------------------------------------
--------------------------------------------------------------------
--------------------------------------------------------------------
--------------------------------------------------------------------
--------------------------------------------------------------------
--------------------------------------------------------------------
----------------------------------------------------------
```

MSSIMNGESGLKMEKLPLEEKHSPHKRAVDAESEEQQLEAQREISETIINVSKRPNSGEGLRIIPPSF
LTASGSVAASISH-NHPTFADALNSISH---------KPLDLSYPKFNSLHPYR----
LYSHPTHMPGSHLG-IPHYPF-------GDTVPMS-AAAAVAAYGSYP--MYHGLTESSMNSSSIK-
FPLSSLLRKRRNSESSSHQSSQSSQD--------------SPPVKSSVSVPELQDESNNDD-D-
TGSSIKK----ARSVPE-----------EKK-DAAYWERRRKNNDAAKRSRDARR-
AKEEEIALRAAFLEQENMKLRAEVSILKNE---LARLHYMV---------
YSC-----------------------------------------------------------------
--------------------------------------------------------------------
-----------------------------------------------------
        'S._purpuratus_gi_390352986_ref_XP_785519.3_'   ------------
--------------------------------------------------------------------
--------------------------------------------------------------------
--------------------------------------------------------------------
--------------------------------------------------------------------
--------------------------------------------------------------------
--------------------------------------------------------------------
--------------------------------------------------MESKEVIE--VKMTK-
P--EKSDSFEVITPVKPEKMKKSAL-DVSN-MVGPS-RPKSAK----------TSD----AS-----
HP---EA----GG---------KPLDLTYSAIVSASQLKS---VYPYSAMLPAAAAA-
AAVTGY-------SYTEPTL-AASAMASYAGYPALMNYPVAENSGSTQSLSSLRLSALLASRRR--
RAGH-------------------------------VVDLQPE---E------PSRKK----
VKPVPD-----------EKK-DIAYWERRRKNNDAAKRSRDARR-
MKEEEIAIRAVYLEQDNMKLRAEVSILKSE---LARLHYMV---------
YNC-----------------------------------------------------------------
--------------------------------------------------------------------
-----------------------------------------------------
        'A._californica_gi_524899363_ref_XP_005106114.1_'        ----
--------------------------------------------------------------------
-------------------------------------------------------------------
MVVEIQRAFTHRPSDGSIGSYSREGILLDRDLGREEDRRLMEEEERMHALASRALGMQHKISDTWKNA
YDTDYPAMADDGVDTTTTTNTTTPSSPPIHGPNGVTNGPNSIVNGLLRKAGSFSDVVCSGGGSLEQKI
QELRAEKEAGEVKKCPSSMPNDQDSPLDFSVKRRSSFSGSLSDDSQASSSSPGHVTDYNAVSPPPAVE
LGGSANHNGDLTRRATPSPGSKLDEQKVMGLGGMNMPPASPMPPLINGMGFFPGLPSNLLANGSAINA
FSQMAAAAAFMDPRGKKNNRPFKAYPKEALQMPLGFFGLPGLTAAALQQGTDSGGGRGGMNSEEIMNM
YNQQLQLLREKQLSASGHVTGAGKTSSPPSSQVQPHPSANTNHNSHNNNNSNNSGSSPHHSHQAQHHH
QQHQ-HHLHQQQHHHQTQG---------SQLHPHHPHHHHHHHHQGNRRLSSSPTHIPSQHQQ-
GGTGGL-------
SPPSPLPRDTSDLMTAASNNSHHHHPHPHLSSNNNNSSHSSSSPPFPTPTTPLPPPSSSTSPS-----
----------------ASMNTTLVSSTSSSSF-A-YNNSRKR----PRSLPD-----------DQK-
DAAYWERRRKNNDAAKRSRDARR-AKEDEIAIRAALLEQENIKLRVEVAALKTE---
TARLRCLL---------
YNS-----------------------------------------------------------------
--------------------------------------------------------------------
-----------------------------------------------------
        'D._melanogaster_gi_24654082_ref_NP_611101.1_'  ------------
--------------------------------------------------------------------
--------------------------------------------------------------------
--------------------------------------------------------------------
--------------------------------------------------------------------
--------------------------------------------------------------------
--------------------------------------------------------------------
--------------------------------------------------------------------
----------------------------------------------------------------
MHSPAQSPILDVT-------------ETVSMHSELDAELESKN-NPSYSISIP-------

```
QNLRL-------TGLG-FP-GMIGAKRSSETLPAFEYIAP------
PSHALQALEFPLMELNRVGVIGGGM-----------------------------
FPGFVHR--------RVR--GE----KRPIPE-----------AQK-
DAKYFERRKRNNEAAKRSRDARK-IREDRIAFRAALLEQENSILRAQVLALRDE---
LQTVRQLLG------
ATTAGGMLSMARQV------------------------------------------------------
--------------------------------------------------------------------
-------------------------------------------------------
        'S._kowalevskii_gi_291234179_ref_XP_002737021.1_'        ----
--------------------------------------------------------------------
--------------------------------------------------------------------
--------------------------------------------------------------------
--------------------------------------------------------------------
--------------------------------------------------------------------
--------------------------------MASEVTTTAAPVIQSDSVDD--
AVDLVVPKKRIRHSSDMLPAEDHSEGSATDTSEPNSPLPTNAVVDKSRHHRVHSDSESDGESSSGSTS
AGKPHGVRLVPQLSQSALEQQFIFPSTSSAVTMPLRMNTNRMMRPFKAYTASRDLLSYAGLGAAGLHQ
HSAAFSLGMPTLFPFGHGMHPSLAPHLMNLAN---------
EAAKTINPNQHYQLQMPRKRPVPSSTVTTASASASSAASASA-------
SSTSSSPRSQISPDGSASSPPGRKHIQSDKRDSTKNAAISSLNSSLSQKPSSSSPSSSS---------
--------------------SSSSSTS--------ALLPPKK----RRSLPD-----------ELK-
DESYWERRRKNNEAAKRSRDIRK-AKEDEIAIRAALLEQENIRLRVEVASLKEE---
TARLRCIL---------
YNG-----------------------------------------------------------------
--------------------------------------------------------------------
-----------------------------------------------------
        'N._vectensis_gi_156408121_ref_XP_001641705.1_' ------------
--------------------------------------------------------------------
--------------------------------------------------------------------
--------------------------------------------------------------------
--------------------------------------------------------------------
--------------------------------------------------------------------
--------------------------------------------------------------------
--------------------------------------------------------------------
--------------------------------------------------------------------
--------------------------------------------------------------------
--------------------------------------------------------------------
-------------------------K------------RQ------RTSVPG-----------EMK-
DQKYWERRLKNNVAAKRSRDLKR-
QKEMTVAKRAQNLEIENEKLRNEV--------------------------------------------
--------------------------------------------------------------------
--------------------------------------------------------------------
-----------------------------------

thyrotroph_embryonic_factor_like_2__Exaiptasia_pallida_ ------------
--------------------------------------------------------------------
--------------------------------------------------------------------
--------------------------------------------------------------------
--------------------------------------------------------------------
--------------------------------------------------------------------
--------------------------------------------------------------------
--------------------------------------------------------------------
--------MSLCKPSNKDYMD--IDEFLSSRDLCGDLATLTEADVDFTNL-TDPVDFSLSAEQ-
LHEYTAN---QLDI---------EEQ-VESVASPVSSDEGRGKSISS--
```

DTGSSFSFSERSIDGE-------
FPQTRSRKSCDKSVTSESEDSSKARTSTWKVSSCPEKQCLATSVEIEGVAYTY---------------
-------------------NLIPTSKS-----------RKPR----RRSVPG-----------SMK-
DEKYWDRRSKNNEAAKRSRDLRR-KKEMLVASKAADLEMENEELRAEVQFLKER---
VKVLNKRLQ------
EKTT----------------------------------------------------------------
--------------------------------------------------------------------
-------------------------------------------------------
'A._digitifera_lcl_XM_015908039.1_prot_XP_015763525.1_1'
------------------------------------------------------------
--------------------------------------------------------------------
--------------------------------------------------------------------
--------------------------------------------------------------------
--------------------------------------------------------------------
--------------------------------------------------------------------
--------------------------------------------------------------------
----------------------------MSVN--FLKNYMD--LDEFLAVKDLCAGLSR-
TESDYSEDDVDQDHFNFIDEEDV-SPGYDLSP--HFDA---------
NNSSLASPGSGSTSDEGRGDSVCSSENTVPSASESPNPPDRN-------DNDKAPRKRKRKS-
ASQCADKETTPTRKRRRKGGNDARRGSTRGIIDEPAP-------------------------------
-----DQTTAAPI-----------LRIA----RKSVDG-----------NEK-
DDKYWERRRKNNQAAKRSRDLKR-QKEIDVKHKAASLEEENRQLREEVKRLKDY---
LKSLGDKVD-----------------------------------------------------------
--------------------------------------------------------------------
--------------------------------------------------------------------
--
'A._californica_gi_524915872_ref_XP_005112723.1_' ----
--------------------------------------------------------------------
--------------------------------------------------------------------
--------------------------------------------------------------------
--------------------------------------------------------------------
--------------------------------------------------------------------
------------------------------
MSDGALDLSMSSSYEVPAHRGAATTTSTSSSRGHDQLSDSGRASSDSTSSDVHDDVKMRRMTSSPPPL
PPHMSVNSMSASLGASAGATSNSNPIPAPSVTYPPPGHPRDHHLPELHHHQQAQRPVSSSTSPPRMGG
GTTGRYSNQNSPQHSPAPGEEDVK-PSLPIDRISSDARVAP-IRPFKMYPVDPFNM---------
YSSSLMAGGGGSPSPVGLP-SLYDSASPTSFPSFPLTPLTSH-------
LMQRKRRAESRENAASNSSAGMAASSGASNNTAGASSSSNNASAVLEGSGNSNSGLAGMSCS------
-------------------NNVTTSSSG---------SLELK----
NDDLSGSDGEKKVKMMDNMKKDDAYWDRRRKNNEAAKRSRDARR-
QKEEEIAMRAAYLEQENLKLRAQVAILKNE---TAKLHYMLY------
NRI-----------------------------------------------------------------
--------------------------------------------------------------------
-------------------------------------------------------
'A._californica_gi_524895551_ref_XP_005104258.1_' ----
--------------------------------------------------------------------
--------------------------------------------------------------------
--------------------------------------------------------------------
--------------------------------------------------------------------
--------------------------------------------------------------------
---------------------------------------
MNPVSSHAAAAAAILAMDPTLNQYYEELGLDDPDTALIRPEMLVANESLLQQREQDLNITPSSGLGVQ
PNFIFPNRMLRGGASALGGVAST-
LGGVSSTLGGVPSALGGVSMSESLSILFDQPSFSPTLSQHLSHQSLMTPKSMGGNG-

IGMSSSQTSGPSASATVGS-LYSQSSVPMHHLGM---------
AGLTTATSSRSTVGSNSSPIDVDSYSVKKEADEGS-SSTLPP-------
PSPSSSSLHSPTTSDAGSDALDSARRGVKRSLSDASSDNKPSTSGLNGSGTAGP--------------
--------------------STSSGAS----------GGPSP----SKSKAD-----------PNK-
DPAYLEKRRKNNESAKRSREARR-SKEEMVALRVVTLEEENMKLRAESSLLEKE---
LDELRHRL---------
FN------------------------------------------------------------------
--------------------------------------------------------------------
-----------------------------------------------------
'A._californica_gi_524895679_ref_XP_005104321.1_'        ----
--------------------------------------------------------------------
--------------------------------------------------------------------
--------------------------------------------------------------------
--------------------------------------------------------------------
--------------------------------------------------------------------
----------------------------------------------------------MS----MPP-
HFLG---SMTLKALLEDP------NLKNPPLVA------------
QSSDQIKVKKEDKDS----------------------------DLPQFNYGSAFLGPNLWDKT--
YDNVDFNLEYMDLDEFLSENGIPVAGEEGRKDQLP----AGPK---------
EEAPPPAMVANNVPIAMSN---SSPRQYLRTPPMSPSQFLASPGP-----SSPTQYPSVMPAPQP-
VSKS-PS-PPVS---PFSVEFQVSEQDLAL-----
SSIPDYHDGYRPKWQAWTHTSGEESSEANGHEDFDPRKRNFSEEELKPQ---PVIKKSK----
KVFVPD-----------EMK-DEKYWNRRHKNNYAAKRSRDARR-
VKENQIAMRAAFLEKENGCLRDEVNKLRKE---NAKLKAK-------
MSKYEPSAGVDPASPISSLPIS----------------------------------------------
--------------------------------------------------------------------
--------------------------------------------------------
'A._californica_gi_524895681_ref_XP_005104322.1_'        ----
--------------------------------------------------------------------
--------------------------------------------------------------------
--------------------------------------------------------------------
--------------------------------------------------------------------
--------------------------------------------------------------------
----------------------------------------------------------MS----MPP-
HFLG---SMTLKALLEDP------NLKNPPL-------------------
IKVKKEDKDS----------------------------DLPQFNYGSAFLGPNLWDKT--
YDNVDFNLEYMDLDEFLSENGIPVAGEEGRKDQLP----AGPK---------
EEAPPPAMVANNVPIAMSN---SSPRQYLRTPPMSPSQFLASPGP-----SSPTQYPSVMPAPQP-
VSKS-PS-PPVS---PFSVEFQVSEQDLAL-----
SSIPDYHDGYRPKWQAWTHTSGEESSEANGHEDFDPRKRNFSEEELKPQ---PVIKKSK----
KVFVPD-----------EMK-DEKYWNRRHKNNYAAKRSRDARR-
VKENQIAMRAAFLEKENGCLRDEVNKLRKE---NAKLKAK-------
MSKYEPSAGVDPASPISSLPIS----------------------------------------------
--------------------------------------------------------------------
--------------------------------------------------------
'A._californica_gi_524895683_ref_XP_005104323.1_'        ----
--------------------------------------------------------------------
--------------------------------------------------------------------
--------------------------------------------------------------------
--------------------------------------------------------------------
--------------------------------------------------------------------
----------------------------------------------------------MS----MPP-
HFLG---SMTLKALLEDP------NLKNPPLVA------------
QSSDQIKVKKEDKDS----------------------------DLPQFNYGSAFLGPNLWDKT--

YDNVDFNLEYMDLDEFLSENGIPVAGEEGRKDQLP----AGPK---------
EEAPPPAMVANNVPIAMSN---SSPRQYLRTPPMSPSQFLASPGP-----SSPTQYPSVMPAPQP-
VSKS-PS-PPVS---PFSVEFQVSEQDLAL-----SSIP-------------------------
GHEDFDPRKRNFSEEELKPQ---PVIKKSK----KVFVPD-----------EMK-
DEKYWNRRHKNNYAAKRSRDARR-VKENQIAMRAAFLEKENGCLRDEVNKLRKE---
NAKLKAK-------
MSKYEPSAGVDPASPISSLPIS----------------------------------------------
--------------------------------------------------------------------
--------------------------------------------------------
        'D._rerio_lcl_NM_131400.1_prot_NP_571475.1_1'   ------------
--------------------------------------------------------------------
--------------------------------------------------------------------
--------------------------------------------------------------------
--------------------------------------------------------------------
--------------------------------------------------------------------
-----------------------------------------------MKPISI--
TMDAGAETSAAFPVVLKKIMETPPPN---LLEGDD------------------
ENDKEKLFESVE--------------------------SGGVSEMG-
PSAALTPAIWEKTIPYDGDTFHLEYMDLEEFLMENGIAAAEN----EQKS----SEKE---------
N--IQLTAE-EPS-TASAV---KTAPAVTLLPVMALDPCEEEVVT----
ITTSSSSSADNKSEENRMTPD-PI-NPDE--IEVDVNFEPDPTDLVL-----
SSIP-------------------------GGELFDPRKHRFSEEELKPQ---PMIKKAK----
KVFVPE-----------DQK-DDKYWQRRKKNNVAAKRSRDARR-
LKENQITVRAAFLERENSALRQEVAELRKD---FGRCKNT-------VARYEAKYG--
AL------------------------------------------------------------------
--------------------------------------------------------------------
---------------------------------------------
        'D._rerio_lcl_XM_005156135.3_prot_XP_005156192.1_1'     ----
--------------------------------------------------------------------
--------------------------------------------------------------------
--------------------------------------------------------------------
--------------------------------------------------------------------
--------------------------------------------------------------------
-------------------------------------------------------MKPISI--
TMDAGAETSAAFPVVLKKIMETPPPN---LLEGDD------------------
ENDKEKLFESVE--------------------------SGGVSEMG-
PSAALTPAIWEKTIPYDGDTFHLEYMDLEEFLMENGIAAAEN----EQKS----SEKE---------
N--IQLTAE-EPS-TASAV---KTAPAVTLLPVMALDPCEEEVVT----
ITTSSSSSADNKSEENRMTPD-PI-NPDE--IEVDVNFEPDPTDLVL-----
SSIP-------------------------GGELFDPRKHRFSEEELKPQ---PMIKKAK----
KVFVPE-----------DQK-DDKYWQRRKKNNVAAKRSRDARR-
LKENQITVRAAFLERENSALRQEVAELRKD---FGRCKNT-------VARYEAKYG--
ALGPEEDV------------------------------------------------------------
--------------------------------------------------------------------
---------------------------------------------
        'D._rerio_lcl_XM_005156136.3_prot_XP_005156193.1_1'     ----
--------------------------------------------------------------------
--------------------------------------------------------------------
--------------------------------------------------------------------
--------------------------------------------------------------------
--------------------------------------------------------------------
--------------------------------------------------------------------
--MSSEIPEIFKALLEYP-FS---LPSIDDN-----------------
ENDKEKLFESVE--------------------------SGGVSEMG-

```
PSAALTPAIWEKTIPYDGDTFHLEYMDLEEFLMENGIAAAEN----EQKS----SEKE---------
N--IQLTAE-EPS-TASAV---KTAPAVTLLPVMALDPCEEEVVT----
ITTSSSSSADNKSEENRMTPD-PI-NPDE--IEVDVNFEPDPTDLVL-----
SSIP-------------------------GGELFDPRKHRFSEEELKPQ---PMIKKAK----
KVFVPE-----------DQK-DDKYWQRRKKNNVAAKRSRDARR-
LKENQITVRAAFLERENSALRQEVAELRKD---FGRCKNT-------VARYEAKYG--
ALGPEEDV------------------------------------------------------------
--------------------------------------------------------------------
---------------------------------------------
        'D._rerio_lcl_NM_001020661.1_prot_NP_001018497.1_1'     ----
--------------------------------------------------------------------
--------------------------------------------------------------------
--------------------------------------------------------------------
--------------------------------------------------------------------
--------------------------------------------------------------------
--------------------------------------------------------------------
--------------MNIPPPN---ILEDQDD-----------------
DIEKEKQASAGDG------------------------
ASAGSGASGGVSASLTPAIWEKTIPYDGETFHLEYMDLDEFLLENGIPVSLE----
EELSRGLEAERR---------DGETQASSE-NSE-EPAAV---PEMPE-
QMQTEQDEDLSDSQTAE----QELSEETTAEPSSVPERATPS-PV-SPED--
IEVNVSFQPDPTDLVL-----SSVP-------------------------
GGELFNPRKHRFSEDELKPQ---PMIKKAK----KVFVPE-----------DAK-
DDKYWSRRKKNNVAAKRSRDARR-LKENQIAVRASFLERENAALRQQVAELRKD---
CGRCQKI-------MALYEAKYG--
LL------------------------------------------------------------------
--------------------------------------------------------------------
---------------------------------------------
        'M._musculus_gi_226958409_ref_NP_705617.2_'     ------------
--------------------------------------------------------------------
--------------------------------------------------------------------
--------------------------------------------------------------------
--------------------------------------------------------------------
--------------------------------------------------------------------
--------------------------------------------------MSSCSQIGVAPAMD--
MPEVLKSLLEHS-------LPWSEKK-------------
ADKEKGKEKLEEDES---------------------------AAASTMA-
VSASLMPPIWDKTIPYDGESFHLEYMDLDEFLLENGIPASP-----THL-------AQ---------
N-LLLPVAELEGK-ESASS---STASP-PSSSTAIFQPSET-----------VSSTESSLEKERE--
TPS-PI-DPSC--VEVDVNFNPDPADLVL-----SSVP-------------------------
GGELFNPRKHRFAEEDLKPQ---PMIKKAK----KVFVPD-----------EQK-
DEKYWTRRKKNNVAAKRSRDARR-LKENQITIRAAFLEKENTALRTEVAELRKE---
VGKCKTI-------VSKYETKYG--
PL------------------------------------------------------------------
--------------------------------------------------------------------
---------------------------------------------
        'M._musculus_gi_568991979_ref_XP_006520806.1_'  ------------
--------------------------------------------------------------------
--------------------------------------------------------------------
--------------------------------------------------------------------
--------------------------------------------------------------------
--------------------------------------------------------------------
--------------------------------------------------------------------
--------------------------------------------------------------------
```

```
----------------MA-VSASLMPPIWDKTIPYDGESFHLEYMDLDEFLLENGIPASP-----
THL-------AQ---------N-LLLPVAELEGK-ESASS---STASP-
PSSSTAIFQPSET-----------VSSTESSLEKERE--TPS-PI-DPSC--
VEVDVNFNPDPADLVL-----SSVP-------------------------
GGELFNPRKHRFAEEDLKPQ---PMIKKAK----KVFVPD-----------EQK-
DEKYWTRRKKNNVAAKRSRDARR-LKENQITIRAAFLEKENTALRTEVAELRKE---
VGKCKTI-------VSKYETKYG--
PL------------------------------------------------------------------
--------------------------------------------------------------------
---------------------------------------------
        'H._sapiens_gi_223972670_ref_NP_001138870.1_'   ------------
--------------------------------------------------------------------
--------------------------------------------------------------------
--------------------------------------------------------------------
--------------------------------------------------------------------
--------------------------------------------------------------------
--------------------------------------------------------------MD--
MPEVLKSLLEHS-------LPWPEKR-------------
TDKEKGKEKLEEDEA---------------------------AAASTMA-
VSASLMPPIWDKTIPYDGESFHLEYMDLDEFLLENGIPASP-----THL-------AH---------
N-LLLPVAELEGK-ESASS---STASP-PSSSTAIFQPSET-----------VSSTESSLEKERE--
TPS-PI-DPNC--VEVDVNFNPDPADLVL-----SSVP-------------------------
GGELFNPRKHKFAEEDLKPQ---PMIKKAK----KVFVPD-----------EQK-
DEKYWTRRKKNNVAAKRSRDARR-LKENQITIRAAFLEKENTALRTEVAELRKE---
VGKCKTI-------VSKYETKYG--
PL------------------------------------------------------------------
--------------------------------------------------------------------
---------------------------------------------
        'M._musculus_gi_8394435_ref_NP_059072.1_'       ------------
--------------------------------------------------------------------
--------------------------------------------------------------------
--------------------------------------------------------------------
--------------------------------------------------------------------
--------------------------------------------------------------------
--------------------------------MSDAGGGKKPPVEPQAGPGPG--
RAAGERGLSGSFPLVLKKLMEN---------PPRETR-------------
LDKEKGKEKLEEDES---------------------------AAASTMA-
VSASLMPPIWDKTIPYDGESFHLEYMDLDEFLLENGIPASP-----THL-------AQ---------
N-LLLPVAELEGK-ESASS---STASP-PSSSTAIFQPSET-----------VSSTESSLEKERE--
TPS-PI-DPSC--VEVDVNFNPDPADLVL-----SSVP-------------------------
GGELFNPRKHRFAEEDLKPQ---PMIKKAK----KVFVPD-----------EQK-
DEKYWTRRKKNNVAAKRSRDARR-LKENQITIRAAFLEKENTALRTEVAELRKE---
VGKCKTI-------VSKYETKYG--
PL------------------------------------------------------------------
--------------------------------------------------------------------
---------------------------------------------
        'Thyrotroph_embryonic_factor_OS_Homo_sapiens_OX_9606_GN_TEF_
PE_1_SV_3'      ----------------------------------------------------
--------------------------------------------------------------------
--------------------------------------------------------------------
--------------------------------------------------------------------
--------------------------------------------------------------------
------------------------------------------------------------
MSDAGGGKKPPVDPQAGPGPGPGRAAGERGLSGSFPLVLKKLMEN---------
```

```
PPREAR-------------LDKEKGKEKLEEDEA---------------------------
AAASTMA-VSASLMPPIWDKTIPYDGESFHLEYMDLDEFLLENGIPASP-----THL-------
AH---------N-LLLPVAELEGK-ESASS---STASP-PSSSTAIFQPSET-----------
VSSTESSLEKERE--TPS-PI-DPNC--VEVDVNFNPDPADLVL-----
SSVP-------------------------GGELFNPRKHKFAEEDLKPQ---PMIKKAK----
KVFVPD-----------EQK-DEKYWTRRKKNNVAAKRSRDARR-
LKENQITIRAAFLEKENTALRTEVAELRKE---VGKCKTI-------VSKYETKYG--
PL------------------------------------------------------------------
--------------------------------------------------------------------
---------------------------------------------

TEF_CHICK       ----------------------------------------------------
--------------------------------------------------------------------
--------------------------------------------------------------------
--------------------------------------------------------------------
--------------------------------------------------------------------
----------------------------------------------------
MPGRAAHQEAAAAGGAAAEPTAAGGSAGAVAQQPEQQGLAGAFPLVLKKLMEN---------
PPRDAR-------------LDKEK-KIKLEEDEA---------------------------
AAASTMA-VSASLMPPIWDKTIPYDGESFHLEYMDLDEFLLENGIPSSP-----THLD-----
LNQ---------N-PLMPVAKLEEK-EPASA---STGSPVSSSSTAVYQQSEA-----------
ASSTESPPQNERN--TPS-PI-DPDC--VEVEVNFNPDPADLVL-----
SSVP-------------------------GGELFNPRKHKFTEEDLKPQ---PMIKKAK----
KVFVPD-----------EQK-DEKYWTRRKKNNVAAKRSRDARR-
LKENQITIRAAFLEKENTALRTEVAELRKE---VGRCKNI-------VSKYETRYG--
PFDLSDSE------------------------------------------------------------
--------------------------------------------------------------------
---------------------------------------------
        'X._laevis_lcl_NM_001086779.1_prot_NP_001080248.1_1'    ----
--------------------------------------------------------------------
--------------------------------------------------------------------
--------------------------------------------------------------------
--------------------------------------------------------------------
--------------------------------------------------------------------
----------------------------------------------------M--SQTES----
IRG--FL--DIPEMLKSLLDYP-------LPQNGT-----------------
EKDKTKLDKDD-----------------------------DHDSMV-
ASASLMPPIWDKTIPYDGESFHLEYMDLDEFLLENGIPSSP-----TQLS----QAIQ---------
TIPLMPVVELECDNEPAST---SSASP--MSPSVLLGNSE------------VED--SDLEDEED--
PPS-PV-DPEK--VEVEVNFNPDPTDLLL-----SSVP-------------------------
GGELFDPRKHRFAEEELKPQ---PMIKKAK----KIYVSE-----------ERK-
DEKYWNRRKKNNIAAKRSRDARR-LKENQITVRAAFLEKENTALRSEVADLRKE---
LGKCRNI-------ISKYETQCG--
LL------------------------------------------------------------------
--------------------------------------------------------------------
---------------------------------------------
        'X._laevis_lcl_XM_018256478.1_prot_XP_018111967.1_1'    ----
--------------------------------------------------------------------
--------------------------------------------------------------------
--------------------------------------------------------------------
--------------------------------------------------------------------
--------------------------------------------------------------------
----------------------------------------------------M--SQTES----
IRG--FL--DIPEMLKSLLDYP-------LPQNGT-----------------
```

```
EKDKTKLDKDD-----------------------------DHDSMV-
ASASLMPPIWDKTIPYDGESFHLEYMDLDEFLLENGIPSSP-----TQLS----QAIQ---------
TIPLMPVVELECDNEPAST---SSASP--MSPSVLLGNSE------------VED--SDLEDEED--
PPS-PV-DPEK--VEVEVNFNPDPTDLLL-----SSVP------------------------
GGELFDPRKHRFAEEELKPQ---PMIKKAK----KIYVSE-----------ERK-
DEKYWNRRKKNNIAAKRSRDARR-LKENQITVRAAFLEKENTALRSEVADLRKE---
LGKCRNI-------ISKYETQCG--
LFEFSDSE------------------------------------------------------------
--------------------------------------------------------------------
---------------------------------------------
        'X._laevis_lcl_XM_018256479.1_prot_XP_018111968.1_1'    ----
--------------------------------------------------------------------
--------------------------------------------------------------------
--------------------------------------------------------------------
--------------------------------------------------------------------
--------------------------------------------------------------------
----------------------------------------------------M--
EASSSPSSPVTPSPILGGSFQLALRKLIENPPKN---FLETSDV-----------------
EKDKTKLDKDD-----------------------------DHDSMV-
ASASLMPPIWDKTIPYDGESFHLEYMDLDEFLLENGIPSSP-----TQLS----QAIQ---------
TIPLMPVVELECDNEPAST---SSASP--MSPSVLLGNSE------------VED--SDLEDEED--
PPS-PV-DPEK--VEVEVNFNPDPTDLLL-----SSVP------------------------
GGELFDPRKHRFAEEELKPQ---PMIKKAK----KIYVSE-----------ERK-
DEKYWNRRKKNNIAAKRSRDARR-LKENQITVRAAFLEKENTALRSEVADLRKE---
LGKCRNI-------ISKYETQCG--
LFEFSDSE------------------------------------------------------------
--------------------------------------------------------------------
---------------------------------------------
        'X._laevis_lcl_NM_001094595.1_prot_NP_001088064.1_1'    ----
--------------------------------------------------------------------
--------------------------------------------------------------------
--------------------------------------------------------------------
--------------------------------------------------------------------
--------------------------------------------------------------------
---------------------------------------------------MMDSSSSPSS--
PVAPSPLVGGSFPLALRKLIENPPKN---FLEERDV-----------------
EKDKTKLDKDD-----------------------------DYDSIV-
ASASLMPPIWDKTIPYDGESFHLEYMDLDEFLLENGISSNP-----TQLA----RAIQ---------
D-PLMSVADLESDIEPAST---LSASP--LSPSVLLGNSE------------DRDLKVDSKAEED--
PPS-PV-EPEK--VEVEVNFNPDPTDLLL-----STVP------------------------
GEELFDPRKHKFAEEELKPQ---PMVKKAK----KIYVPE-----------DLK-
DEKYWNRRKKNNVAAKRSREARR-LKENQITVRAAFLEKENTALRSEVADLRKE---
LGKSRNI-------ISKYEKQFG--
LFDFNDSE------------------------------------------------------------
--------------------------------------------------------------------
---------------------------------------------
        'X._laevis_lcl_XM_018259674.1_prot_XP_018115163.1_1'    ----
--------------------------------------------------------------------
--------------------------------------------------------------------
--------------------------------------------------------------------
--------------------------------------------------------------------
--------------------------------------------------------------------
---------------------------------------------------MMDSSSSPSS--
PVAPSPLVGGSFPLALRKLIENPPKN---FLEERDVG-------------
```

```
EGHEKDKTKLDKDD-----------------------------DYDSIV-
ASASLMPPIWDKTIPYDGESFHLEYMDLDEFLLENGISSNP-----TQLA----RAIQ---------
D-PLMSVADLESDIEPAST---LSASP--LSPSVLLGNSE------------DRDLKVDSKAEED--
PPS-PV-EPEK--VEVEVNFNPDPTDLLL-----STVP-------------------------
GEELFDPRKHKFAEEELKPQ---PMVKKAK----KIYVPE-----------DLK-
DEKYWNRRKKNNVAAKRSREARR-LKENQITVRAAFLEKENTALRSEVADLRKE---
LGKSRNI-------ISKYEKQFG--
LFDFNDSE------------------------------------------------------------
--------------------------------------------------------------------
---------------------------------------------
        'X._laevis_lcl_XM_018259675.1_prot_XP_018115164.1_1'    ----
--------------------------------------------------------------------
--------------------------------------------------------------------
--------------------------------------------------------------------
--------------------------------------------------------------------
--------------------------------------------------------------------
----------------------------------------------------M--SQTES----
IRG--FL--DIPEMLKSLLDYP-------LPQHMT-----------------
EKDKTKLDKDD-----------------------------DYDSIV-
ASASLMPPIWDKTIPYDGESFHLEYMDLDEFLLENGISSNP-----TQLA----RAIQ---------
D-PLMSVADLESDIEPAST---LSASP--LSPSVLLGNSE------------DRDLKVDSKAEED--
PPS-PV-EPEK--VEVEVNFNPDPTDLLL-----STVP-------------------------
GEELFDPRKHKFAEEELKPQ---PMVKKAK----KIYVPE-----------DLK-
DEKYWNRRKKNNVAAKRSREARR-LKENQITVRAAFLEKENTALRSEVADLRKE---
LGKSRNI-------ISKYEKQFG--
LFDFNDSE------------------------------------------------------------
--------------------------------------------------------------------
---------------------------------------------
        'X._laevis_lcl_XM_018259676.1_prot_XP_018115165.1_1'    ----
--------------------------------------------------------------------
--------------------------------------------------------------------
--------------------------------------------------------------------
--------------------------------------------------------------------
--------------------------------------------------------------------
----------------------------------------------------M--SQTES----
IRG--FL--DIPEMLKSLLDYP-------LPQHMT-----------------
EKDKTKLDKDD-----------------------------DYDSIV-
ASASLMPPIWDKTIPYDGESFHLEYMDLDEFLLENGISSNP-----TQLA----RAIQ---------
D-PLMSVADLESDIEPAST---LSASP--LSPSVLLGNSE------------DRDLKVDSKAEED--
PPS-PV-EPEK--VEVEVNFNPDPTDLLL-----STVP-------------------------
GEELFDPRKHKFAEEELKPQ---PMVKKAK----KIYVPE-----------DLK-
DEKYWNRRKKNNVAAKRSREARR-LKENQITVRAAFLEKENTALRSEVADLRKE---
LGKSRNI-------ISKYEKQFG--
LL------------------------------------------------------------------
--------------------------------------------------------------------
---------------------------------------------
        'H._sapiens_gi_530412024_ref_XP_005257326.1_'   ------------
--------------------------------------------------------------------
--------------------------------------------------------------------
--------------------------------------------------------------------
--------------------------------------------------------------------
--------------------------------------------------------------------
-----------------------------------------------MEKMSRPLPLNP-TFIPP-
PYGVLRSLLENPLKL---PLHHEDA--------------
```

```
FSKDKDKEKKLDDE-----------------------------
SNSPTVPQSAFLGPTLWDKTLPYDGDTFQLEYMDLEEFLSENGIPPSP-----SQHD----
HSPH---------PPGLQPASS-AAP-SVMDL---SSRASAPLHPGIPSPNCMQS----------
PIRPGQLLPANRN--TPS-PI-DPDT--IQVPVGYEPDPADLAL-----
SSIP-------------------------GQEMFDPRKRKFSEEELKPQ---PMIKKAR----
KVFIPD-----------DLKQDDKYWARRRKNNMAAKRSRDARR-
LKENQIAIRASFLEKENSALRQEVADLRKE---LGKCKNI-------LAKYEARHG--
PL------------------------------------------------------------------
--------------------------------------------------------------------
---------------------------------------------

HLF_HUMAN       ----------------------------------------------------
--------------------------------------------------------------------
--------------------------------------------------------------------
--------------------------------------------------------------------
--------------------------------------------------------------------
--------------------------------------------------------------------
-------MEKMSRPLPLNP-TFIPP-PYGVLRSLLENPLKL---PLHHEDA--------------
FSKDKDKEKKLDDE-----------------------------
SNSPTVPQSAFLGPTLWDKTLPYDGDTFQLEYMDLEEFLSENGIPPSP-----SQHD----
HSPH---------PPGLQPASS-AAP-SVMDL---SSRASAPLHPGIPSPNCMQS----------
PIRPGQLLPANRN--TPS-PI-DPDT--IQVPVGYEPDPADLAL-----
SSIP-------------------------GQEMFDPRKRKFSEEELKPQ---PMIKKAR----
KVFIPD-----------DLK-DDKYWARRRKNNMAAKRSRDARR-
LKENQIAIRASFLEKENSALRQEVADLRKE---LGKCKNI-------LAKYEARHG--
PL------------------------------------------------------------------
--------------------------------------------------------------------
---------------------------------------------
        'M._musculus_gi_31982951_ref_NP_766151.1_'      ------------
--------------------------------------------------------------------
--------------------------------------------------------------------
--------------------------------------------------------------------
--------------------------------------------------------------------
--------------------------------------------------------------------
-----------------------------------------------MEKMSRQLPLNP-TFIPP-
PYGVLRSLLENPLKL---PLHPEDA--------------
FSKEKDKGKKLDDE-----------------------------
SSSPTVPQSAFLGPTLWDKTLPYDGDTFQLEYMDLEEFLSENGIPPSP-----SQHD----
HSPH---------PPGLQPASS-TAP-SVMDL---SSRATAPLHPGIPSPNCMQS----------
PIRPGQLLPANRN--TPS-PI-DPDT--IQVPVGYEPDPADLAL-----
SSIP-------------------------GQEMFDPRKRKFSEEELKPQ---PMIKKAR----
KVFIPD-----------DLK-DDKYWARRRKNNMAAKRSRDARR-
LKENQIAIRASFLEKENSALRQEVADLRKE---LGKCKNI-------LAKYEARHG--
PL------------------------------------------------------------------
--------------------------------------------------------------------
---------------------------------------------
        'M._musculus_gi_568973277_ref_XP_006533067.1_'  ------------
--------------------------------------------------------------------
--------------------------------------------------------------------
--------------------------------------------------------------------
--------------------------------------------------------------------
--------------------------------------------------------------------
-----------------------------------------------MEKMSRQLPLNP-TFIPP-
PYGVLRSLLENPLKL---PLHPEDA--------------
```

```
FSKEKDKGKKLDDE-----------------------------
SSSPTVPQSAFLGPTLWDKTLPYDGDTFQLEYMDLEEFLSENGIPPSP-----SQHD----
HSPH---------PPGLQPASS-TAP-SVMDL---SSRATAPLHPGIPSPNCMQS----------
PIRPGQLLPANRN--TPS-PI-DPDT--IQVPVGYEPDPADLAL-----
SSIP-------------------------GQEMFDPRKRKFSEEELKPQ---PMIKKAR----
KVFIPD-----------DLKQDDKYWARRRKNNMAAKRSRDARR-
LKENQIAIRASFLEKENSALRQEVADLRKE---LGKCKNI-------LAKYEARHG--
PL------------------------------------------------------------------
--------------------------------------------------------------------
---------------------------------------------
        'M._musculus_gi_568973279_ref_XP_006533068.1_'  ------------
--------------------------------------------------------------------
--------------------------------------------------------------------
--------------------------------------------------------------------
--------------------------------------------------------------------
--------------------------------------------------------------------
--------------------------------------------------------------------
--------------------------------------------------------------------
--------------------------------------------MDLEEFLSENGIPPSP-----
SQHD----HSPH---------PPGLQPASS-TAP-SVMDL---
SSRATAPLHPGIPSPNCMQS----------PIRPGQLLPANRN--TPS-PI-DPDT--
IQVPVGYEPDPADLAL-----SSIP-------------------------
GQEMFDPRKRKFSEEELKPQ---PMIKKAR----KVFIPD-----------
DLKQDDKYWARRRKNNMAAKRSRDARR-LKENQIAIRASFLEKENSALRQEVADLRKE---
LGKCKNI-------LAKYEARHG--
PL------------------------------------------------------------------
--------------------------------------------------------------------
---------------------------------------------
        'M._musculus_gi_568973283_ref_XP_006533070.1_'  ------------
--------------------------------------------------------------------
--------------------------------------------------------------------
--------------------------------------------------------------------
--------------------------------------------------------------------
--------------------------------------------------------------------
--------------------------------------------------------------------
--------------------------------------------------------------------
--------------------------------------------MDLEEFLSENGIPPSP-----
SQHD----HSPH---------PPGLQPASS-TAP-SVMDL---
SSRATAPLHPGIPSPNCMQS----------PIRPGQLLPANRN--TPS-PI-DPDT--
IQVPVGYEPDPADLAL-----SSIP-------------------------
GQEMFDPRKRKFSEEELKPQ---PMIKKAR----KVFIPD-----------DLK-
DDKYWARRRKNNMAAKRSRDARR-LKENQIAIRASFLEKENSALRQEVADLRKE---
LGKCKNI-------LAKYEARHG--
PL------------------------------------------------------------------
--------------------------------------------------------------------
---------------------------------------------
        'X._laevis_lcl_XM_018235125.1_prot_XP_018090614.1_1'    ----
--------------------------------------------------------------------
--------------------------------------------------------------------
--------------------------------------------------------------------
--------------------------------------------------------------------
--------------------------------------------------------------------
-------------------------------------------------------MEKMSRVLPLNP-
TFIPP-TYGVLKSLLENPLKL---PMHHEDA--------------FCKEKEK--
```

KLEDD-----------------------------
NTASTVPQSAFLGPTLWDKTLPYDGDTFQLEYMDLEEFLSENGIPPSQ-----SSHE----
LSHH---------QPSHPQASA-TSP-SVIDL---SNRASTSVLPGLVPHNCMHS----------
PVRPGQILPANRN--TPS-PI-DPES--IQVAVGYEPDPADLAL-----
SSIP-------------------------GQEMFDPRKRKFSDEELKPQ---PMIKKAR----
KIFISE-----------DLKQDEKYWARRKKNNLAAKRSRDARR-
LKENQIAIRASFLEKENSALRMEVVDLRKE---LGKCKNI-------LAKYEARHG--
PL------------------------------------------------------------------
--------------------------------------------------------------------
---------------------------------------------
        'X._laevis_lcl_XM_018235126.1_prot_XP_018090615.1_1'    ----
--------------------------------------------------------------------
--------------------------------------------------------------------
--------------------------------------------------------------------
--------------------------------------------------------------------
--------------------------------------------------------------------
-------------------------------------------------------MEKMSRVLPLNP-
TFIPP-TYGVLKSLLENPLKL---PMHHEDA--------------FCKEKEK--
KLEDD-----------------------------
NTASTVPQSAFLGPTLWDKTLPYDGDTFQLEYMDLEEFLSENGIPPSQ-----SSHE----
LSHH---------QPSHPQASA-TSP-SVIDL---SNRASTSVLPGLVPHNCMHS----------
PVRPGQILPANRN--TPS-PI-DPES--IQVAVGYEPDPADLAL-----
SSIP-------------------------GQEMFDPRKRKFSDEELKPQ---PMIKKAR----
KIFISE-----------DLK-DEKYWARRKKNNLAAKRSRDARR-
LKENQIAIRASFLEKENSALRMEVVDLRKE---LGKCKNI-------LAKYEARHG--
PL------------------------------------------------------------------
--------------------------------------------------------------------
---------------------------------------------
        'X._laevis_lcl_XM_018240540.1_prot_XP_018096029.1_1'    ----
--------------------------------------------------------------------
--------------------------------------------------------------------
--------------------------------------------------------------------
--------------------------------------------------------------------
--------------------------------------------------------------------
-------------------------------------------------------MEKMSRVLPLNS-
TFIPP-TYGVLKSLLENPLKL---PLHHEDA--------------FCKEKEK--
KLEDD-----------------------------
NTASTVPQSAFLGPTLWDKTLPYDGDTFQLEYMDLEEFLSENGIPPSQ-----SSHE----
LSHH---------QPSHPQASA-TSP-SVIDL---SNRASTSVLPGLVPHNCMHS----------
PVRPGQILAANRN--TPS-PI-DPES--IQVVVGYEPDPADLAL-----
SSIP-------------------------GQEMFDPRKRKFSEEELKPQ---PMIKKAR----
KVFISE-----------DLKQDEKYWTRRKKNNLAAKRSRDARR-
LKENQIAIRASFLEKENSALRMEVADLRKE---LGKCKNV-------LAKYEARHG--
HL------------------------------------------------------------------
--------------------------------------------------------------------
---------------------------------------------
        'X._laevis_lcl_XM_018240541.1_prot_XP_018096030.1_1'    ----
--------------------------------------------------------------------
--------------------------------------------------------------------
--------------------------------------------------------------------
--------------------------------------------------------------------
--------------------------------------------------------------------
-------------------------------------------------------MEKMSRVLPLNS-
TFIPP-TYGVLKSLLENPLKL---PLHHEDA--------------FCKEKEK--

```
KLEDD-----------------------------
NTASTVPQSAFLGPTLWDKTLPYDGDTFQLEYMDLEEFLSENGIPPSQ-----SSHE----
LSHH---------QPSHPQASA-TSP-SVIDL---SNRASTSVLPGLVPHNCMHS----------
PVRPGQILAANRN--TPS-PI-DPES--IQVVVGYEPDPADLAL-----
SSIP-------------------------GQEMFDPRKRKFSEEELKPQ---PMIKKAR----
KVFISE-----------DLK-DEKYWTRRKKNNLAAKRSRDARR-
LKENQIAIRASFLEKENSALRMEVADLRKE---LGKCKNV-------LAKYEARHG--
HL------------------------------------------------------------------
--------------------------------------------------------------------
---------------------------------------------
        'D._rerio_lcl_NM_001077334.2_prot_NP_001070802.1_1'     ----
--------------------------------------------------------------------
--------------------------------------------------------------------
--------------------------------------------------------------------
--------------------------------------------------------------------
--------------------------------------------------------------------
-------------------------------------------------------MEKMSRPLPINA-
TFLPP-THGVLKSLLENPMKL---PFHHDEG--------------
FGKEKEKEKKLEDD-----------------------------
ASTLNTPQSAFLGPTLWDKTLPYNADNFQLEYMDLEEFLLENNIPANP-----QSEQ----
SQPS---------QPPLQPPSAPPTP-SVVDL---SNRDNSSSHNGMVAQNCLQN----------
PTRPG--LPASRD--TPS-PI-DPDS--IQVPLAYEPDPADLAL-----
SSVP-------------------------GQEIFDPRKRKFSAEELKPQ---PMIKKAR----
KVFIPE-----------DLK-DDRYWARRRKNNIAAKRSRDARR-
LKENQIAIRAGFLEKENAALRAEVADLRKE---LGRCKNV-------LAKYEARHG--
PL------------------------------------------------------------------
--------------------------------------------------------------------
---------------------------------------------
        'D._rerio_lcl_NM_001197059.1_prot_NP_001183988.1_1'     ----
--------------------------------------------------------------------
--------------------------------------------------------------------
--------------------------------------------------------------------
--------------------------------------------------------------------
--------------------------------------------------------------------
----------------------------------------------------------MSRQLTMNP-
AFLPPQTNGVLKALLEKPLKL---PLHQDEG--------------
YEKERDKVKKLDEE-----------------------------GNP---
PQSAFLGPTLWDKTLSYDGDSFQLEYMDLEEFLSENGIPSSP-----AQHDQNL-HQHHHQQQQQ--
HQQQQQQVSMPQGPISVMDL---SSR---SIHTAISPQNCLHS----------PGRS--VLPPSRN--
TPS-PV-DPEA--LHIPVSYEPDPADLAL-----SSVP-------------------------
GQEVFDPRKRKFSEEELKPQ---PMIKKAR----KIFIPD-----------DLK-
DEKYWARRRKNNVAAKRSRDARR-LKENQIAIRAGFLEKENMALRQEVADLRKE---
LGRCKNI-------LTKYEAQHG--
PL------------------------------------------------------------------
--------------------------------------------------------------------
---------------------------------------------
        'D._rerio_lcl_XM_005156334.4_prot_XP_005156391.1_1'     ----
--------------------------------------------------------------------
--------------------------------------------------------------------
--------------------------------------------------------------------
--------------------------------------------------------------------
--------------------------------------------------------------------
----------------------------------------------------------MSRQLTMNP-
AFLPPQTNGVLKALLEKPLKL---PLHQDEG--------------
```

YGKERDKVKKLDEE-----------------------------GNP---
PQSAFLGPTLWDKTLSYDGDSFQLEYMDLEEFLSENGIPSSP-----AQHDQNL-
HQHHHQQQQQQQHQQQQQQVSMPQGPISVMDL---SSR---SIHTAISPQNCLHS----------
PGRS--VLPPSRN--TPS-PV-DPEA--LHIPVSYEPDPADLAL-----
SSVP-------------------------GQEVFDPRKRKFSEEELKPQ---PMIKKAR----
KIFIPD-----------DLKQDEKYWARRRKNNVAAKRSRDARR-
LKENQIAIRAGFLEKENMALRQEVADLRKE---LGRCKNI-------LTKYEAQHG--
PL------------------------------------------------------------------
--------------------------------------------------------------------
---------------------------------------------

'D._rerio_lcl_NM_001197062.1_prot_NP_001183991.1_1'      ----
--------------------------------------------------------------------
--------------------------------------------------------------------
--------------------------------------------------------------------
--------------------------------------------------------------------
--------------------------------------------------------------------
----------------------------------
MARPLSQLLPPDLPSAGASPQFGNSSQAGVTHNGGHLN--STGNLKSLLQLPVKC---
DQRVKDCG-------------EMK--GKER-LDIDED-
SLGRCPLRNGCSNGLVSDSNGAGTGSFSNNSNNNNSFLGPLLWERTLPCDGGLFQLQYMDLEEFLTEN
GMSSMHN-TSNSTSAQIPSQSSQSAVPNQG-SQCLPTSPPHCSSSSSPTS---
ATASSPSLLGLDMHTPQSMMGSTDCLHGTPPGSLEPTPSPSSTT--CPPLPT-
PPATNCNELLASFDPDPADVAL-----SSVP-------------------------
GQEAFDPRRHHFSDEDLKPQ---PMIKKAR----KMLVPE-----------DLK-
DEKYWSRRCKNNEAAKRSRDARR-LKENQISVRAAFLERENAALRQEVADMRKE---
LGRCRNI-------LNKYESHHL--
DQ------------------------------------------------------------------
--------------------------------------------------------------------
---------------------------------------------

'D._rerio_lcl_XM_005157897.4_prot_XP_005157954.1_1'      ----
--------------------------------------------------------------------
--------------------------------------------------------------------
--------------------------------------------------------------------
--------------------------------------------------------------------
--------------------------------------------------------------------
---------------------------------
MMARPLSQLLPPDLPSAGASPQFGNSSQAGVTHNGGHLN--STGNLKSLLQLPVKC---
DQRVKDCG-------------EMK--GKER-LDIDED-
SLGRCPLRNGCSNGLVTDSNGAGTGSFSNNSNNNNSFLGPLLWERTLPCDGGLFQLQYMDLEEFLTEN
GMSSMHN-TSNSTSAQIPSQSSQSAVPNQG-SQCLPTSPPHCSSSSSPTS---
ATASSPSLLGLDMHTPQSMMGSTDCLHGTPPGSLEPTPSPSSTT--CPPLPT-
PPATNCNELLAPFDPDPADVAL-----SSVP-------------------------
GQEAFDPRRHHFSDEDLKPQ---PMIKKAR----KMLVPE-----------DLK-
DEKYWSRRCKNNEAAKRSRDARR-LKENQISVRAAFLERENAALRQEVADMRKE---
LGRCRNI-------LNKYESHHL--
DQ------------------------------------------------------------------
--------------------------------------------------------------------
---------------------------------------------

'D._rerio_lcl_NM_001197060.1_prot_NP_001183989.1_1'      ----
--------------------------------------------------------------------
--------------------------------------------------------------------
--------------------------------------------------------------------
--------------------------------------------------------------------
--------------------------------------------------------------------

---------------------------------MSRPISQILPPDLP-
AGTSPQLGPANPAGTTTNG-HLNN-SMANLKTLLQLPIKG---DQRGKDCC-------------
EMKVSDKDKPLDSDED-SLG-----GGGGGG--GGMNG-GNGVLR--
STNQSAFLGPLLWERTLPCDGGLFQLQYMDLEEFLTENGMCMPSGNTCSTAAQVPSQSTQSAVPSQS
-SQCPPSSSPPCSSSASSISSLSSSSSSSSLLGLDVPQGPGLLGGPECLHGAQ--TVPPDPS-
QSPS--CPPPPV-VPPTNAADVMVNFDPDPADLAL-----
SSVP-------------------------GQEAFDPRRHRFSEEELKPQ---PMIKKAR----
KMLVPD-----------EQK-DDKYWCRRLKNDEAAKRSRDARR-
LKENQISVRAAFLERENAALRQEVADMRKE---LGRCRNI-------INKYESRHG--
DL-----------------------------------------------------------------
-------------------------------------------------------------------
--------------------------------------------

'M._musculus_gi_170650717_ref_NP_058670.2_' ------------
-------------------------------------------------------------------
-------------------------------------------------------------------
-------------------------------------------------------------------
-------------------------------------------------------------------
-------------------------------------------------------------------
-------------------------MARPLSDRTPGPLL-------LG--GPAG-APPGG--G--
ALLGLRSLLQG-------NSKPKEPA-------------
SCLLKEKERKATLPSAPVPGPGLETAGPADAPSGAVSGGGSPRGRSGPVAGPSLFAPLLWERTLPFG-
---DVEYVDLDAFLLEHGLPPSPP----PPGGLSPAPSPARTPA--------
PSPGPGSCSSSSPRS-----SPGHAPARATLGAAGGHRAG---------------LT--SRD--TPS-
PV-DPDT--VEVLMTFEPDPADLAL-----SSIP-------------------------
GHETFDPRRHRFSEEELKPQ---PIMKKAR----KVQVPE-----------EQK-
DEKYWSRRYKNNEAAKRSRDARR-LKENQISVRAAFLEKENALLRQEVVAVRQE---
LSHYRAV-------LSRYQAQHG--
TL-----------------------------------------------------------------
-------------------------------------------------------------------
--------------------------------------------

'M._musculus_gi_568947265_ref_XP_006540659.1_' ------------
-------------------------------------------------------------------
-------------------------------------------------------------------
-------------------------------------------------------------------
-------------------------------------------------------------------
-------------------------------------------------------------------
-------------------------MARPLSDRTPGPLL-------LG--GPAG-APPGG--G--
ALLGLRSLLQG-------NSKPKEPA-------------
SC-----------------------------------------------------------------
-------------------------------------------------------------------
--------------------------------------LT--SRD--TPS-PV-DPDT--
VEVLMTFEPDPADLAL-----SSIP-------------------------
GHETFDPRRHRFSEEELKPQ---PIMKKAR----KVQVPE-----------EQK-
DEKYWSRRYKNNEAAKRSRDARR-LKENQISVRAAFLEKENALLRQEVVAVRQE---
LSHYRAV-------LSRYQAQHG--
TL-----------------------------------------------------------------
-------------------------------------------------------------------
--------------------------------------------

DBP_HUMAN ---------------------------------------------------
-------------------------------------------------------------------
-------------------------------------------------------------------
-------------------------------------------------------------------
-------------------------------------------------------------------

```
------------------------------------------------------
MARPVSDRTPAPLL-------LG--GPAG-TPPGG--G--ALLGLRSLLQG-------
TSKPKEPA-------------
SCLLKEKERKAALPAATTPGPGLETAGPADAPAGAVVGGGSPRGRPGPVAPAGLLAPLLWERTLPFG-
---DVEYVDLDAFLLEHGLPPSPP----PPGGPSPEPSPARTPA--------
PSPGPGSCGSASPRS-----SPGHAPARAALGTASGHRAG---------------LT--SRD--TPS-
PV-DPDT--VEVLMTFEPDPADLAL-----SSIP-------------------------
GHETFDPRRHRFSEEELKPQ---PIMKKAR----KIQVPE-----------EQK-
DEKYWSRRYKNNEAAKRSRDARR-LKENQISVRAAFLEKENALLRQEVVAVRQE---
LSHYRAV-------LSRYQAQHG--
AL------------------------------------------------------------------
--------------------------------------------------------------------
---------------------------------------------
        'M._musculus_gi_568947267_ref_XP_006540660.1_'  ------------
--------------------------------------------------------------------
--------------------------------------------------------------------
--------------------------------------------------------------------
--------------------------------------------------------------------
--------------------------------------------------------------------
--------------------------MLE-L--RPKGTRR-------EA--TPIK-TRSLD--V--
GRLG-TPVPDP-------TSRQK-PV-------------LH----------
IP------------------------------------------------------------------
---------------------------------------------------
SPR-----------------------------------------LT--SRD--TPS-PV-DPDT--
VEVLMTFEPDPADLAL-----SSIP-------------------------
GHETFDPRRHRFSEEELKPQ---PIMKKAR----KVQVPE-----------EQK-
DEKYWSRRYKNNEAAKRSRDARR-LKENQISVRAAFLEKENALLRQEVVAVRQE---
LSHYRAV-------LSRYQAQHG--
TL------------------------------------------------------------------
--------------------------------------------------------------------
---------------------------------------------
        'X._laevis_lcl_XM_018226265.1_prot_XP_018081754.1_1'    ----
--------------------------------------------------------------------
--------------------------------------------------------------------
--------------------------------------------------------------------
--------------------------------------------------------------------
--------------------------------------------------------------------
----------------------------------MARPTTQLPASEKL-------QG--
HLLHPTPPIGVPGS-GMVSLKSLLQGPIKGQ--DRRTREMS-------------NCIMKDKERK--
LED--------DMTGP----------------RPSHCA---LFGSLLWERALPCN----
EIEYVDLDEFLRENGLPPSPPHQPFSPANLTPPPSNQSVVD--------
LSRPASCASSSSTCS-----SPVQSIMDSEYEHPSSKAGQ---------------MTPVSSD--SPS-
PE-DPES--IEVISKFDLDPADLAL-----SSVP-------------------------
GHETFDPRKHRFSEEELKPQ---PIMKKAR----KIQVPD-----------NHK-
DEKYWNRRYKNNEAAKRSRDARR-LKENQITVRAAFLEKENSVLRQEVSRIRQE---
LSRYRNI-------LSKYESQHG--
AL------------------------------------------------------------------
--------------------------------------------------------------------
---------------------------------------------
        'X._laevis_lcl_XM_018228082.1_prot_XP_018083571.1_1'    ----
--------------------------------------------------------------------
--------------------------------------------------------------------
--------------------------------------------------------------------
--------------------------------------------------------------------
```

--------------------------------------------------------------------
----------------------------------MSRPTTQLTTCEKL-------QG--
HQLHQTPPLGLPGP-GMVSLKSLLQGSIKGP--DRRTRELS-------------NSIMKDKERK--
LED--------DMTGP----------------RPNHCA---FLGSLVWERTVPYN----
EIEYVDLDEFLRENGLPSSPPHQPFSPANLTPPPSNQSVFD--------
LSRPASCASSTSACS-----SPVQSMVDSEY--PSSNAGE---------------MTPVSRD--SPS-
PV-DPES--IEVISKFDLDPADLAL-----SSVP-------------------------
GHETFDPRKHRFSEEDLKPQ---PIMKKAK----KIQVPH-----------EQK-
DEKYWNRRYKNNEAAKRSRDARR-LKENQITVRAAFLEKENSVLRQEVSRIRQE---
LSRYRNI-------LSKYESQHG--
AL------------------------------------------------------------------
--------------------------------------------------------------------
---------------------------------------------

'S._purpuratus_gi_193788685_ref_NP_001123286.1_' ----
--------------------------------------------------------------------
--------------------------------------------------------------------
--------------------------------------------------------------------
--------------------------------------------------------------------
--------------------------------------------------------------------
-------------------------------------------------------MDFMEG---
MEE--RLSDIRGITLKSLLENPS-----FLQPPTG---------------
NNNEDKNAKKDKEAG-----------------------LFGLDIGSEDVASAFLGPTLWEKT-PYD--
DLKLEYMDLDEFLSENGIAVSDG---KTSSQ----EKPQEK-----
VVSEGASPVSRGASPVSETSTDKENRYLMSTQSPTPSPPPPMSP----------
DSISEELVYSPTN--TMS-PVREPSP--VHVNVPFELSETDVAL-----
ATAP-------------------------GQDTFDPTECTFTAEELKPQ---PMIKKSR----
KIYVPD-----------EQK-DDKYWERRRKNNIAAKRSREARR-
VKENQIVIRASYLGKENSALRDDVLELKKE---NSTLKKI-------
IASYEKRLKTAKMD------------------------------------------------------
--------------------------------------------------------------------
--------------------------------------------------------

'D._melanogaster_gi_24660427_ref_NP_729297.1_' ------------
--------------------------------------------------------------------
--------------------------------------------------------------------
--------------------------------------------------------------------
--------------------------------------------------------------------
--------------------------------------------------------------------
------------------------------------------------------------------
MSEPKENIMD-LSVR------G-----------------
KSTSNSKEIICPDDKYK-----------------------
EEGDIWNVEAQTAFLGPNLWDKTLPYDADLKVTQYADLDEFLSENNIP--------DGLP-----
GTH------------LGHSSGLGHRSDSLG-----HAAG--
LSLGLG-----------------------HITTKRE-RSPS-PS---DC--
ISPDTLNPPSPAESTF-----SFAS-------------------------
SGRDFDPRTRAFSDEELKPQ---PMIKKSR----KQFVPD-----------ELK-
DDKYWARRRKNNIAAKRSRDARR-QKENQIAMRARYLEKENATLHQEVEQLKQE---
NMDLRAR-------
LSKFQDV-------------------------------------------------------------
--------------------------------------------------------------------
--------------------------------------------------------

'D._melanogaster_gi_442630879_ref_NP_001261547.1_' ----
--------------------------------------------------------------------
--------------------------------------------------------------------
--------------------------------------------------------------------

--------------------------------------------------------------------
--------------------------------------------------------------------
--------------------------------------------------------------------
------MSEPKENIMD-LSVR------GRKS---
PFLGFICKSNGKSTSNSKEIICPDDKYK-----------------------
EEGDIWNVEAQTAFLGPNLWDKTLPYDADLKVTQYADLDEFLSENNIP--------DGLP-----
GTH------------LGHSSGLGHRSDSLG-----HAAG--
LSLGLG-----------------------HITTKRE-RSPS-PS---DC--
ISPDTLNPPSPAESTF-----SFAS-------------------------
SGRDFDPRTRAFSDEELKPQ---PMIKKSR----KQFVPD-----------ELK-
DDKYWARRRKNNIAAKRSRDARR-QKENQIAMRARYLEKENATLHQEVEQLKQE---
NMDLRAR-------
LSKFQDV-------------------------------------------------------------
--------------------------------------------------------------------
--------------------------------------------------------

'D._melanogaster_gi_24660458_ref_NP_729301.1_'  ------------
-----------------------------
MSSDTNSCTSFFAQEHANATLSQYFQQLNSHVAAAAAGNSGGNSSSSNNNNSSGNNSSSGNSDSGNDG
SMTGSSGTMDNHRSSVSSNDSGKSSATGGGGAGATSNNGVNAWALQQQQQTAALHHQQQQQHQQQQQH
QQHLQQHHQAHHQHVLQQQQQQQQQHSQHQHQQQHHGHPLHPHHTHQQQQQQQQQQQPPPQHHQQQQQ
QHAQQQQQQQHHAAQLAHTLSSAAVAAAAGNTGGNTNPHSIFGTGNFHYKTNNSWTLPTLTYQRIYQEN
PHYQRNSFMDPQSNAAAAAAAAAAASCNAAAVAAAAAVASGNQGSNGNASGNGNAVVAATGNGNGGNPG
QNNNNNNGNNSGNSNNNSNNNVSSV-QHVANAVAAAVIANEHHNHLNSLKARFQPASSGRKS---
PFLGFICKSNGKSTSNSKEIICPDDKYK-----------------------
EEGDIWNVEAQTAFLGPNLWDKTLPYDADLKVTQYADLDEFLSENNIP--------DGLP-----
GTH------------LGHSSGLGHRSDSLG-----HAAG--
LSLGLG-----------------------HITTKRE-RSPS-PS---DC--
ISPDTLNPPSPAESTF-----SFAS-------------------------
SGRDFDPRTRAFSDEELKPQ---PMIKKSR----KQFVPD-----------ELK-
DDKYWARRRKNNIAAKRSRDARR-QKENQIAMRARYLEKENATLHQEVEQLKQE---
NMDLRAR-------
LSKFQDV-------------------------------------------------------------
--------------------------------------------------------------------
--------------------------------------------------------

'D._melanogaster_gi_281365802_ref_NP_729302.2_' ------------
-----------------------------
MSSDTNSCTSFFAQEHANATLSQYFQQLNSHVAAAAAGNSGGNSSSSNNNNSSGNNSSSGNSDSGNDG
SMTGSSGTMDNHRSSVSSNDSGKSSATGGGGAGATSNNGVNAWALQQQQQTAALHHQQQQQHQQQQQH
QQHLQQHHQAHHQHVLQQQQQQQQQHSQHQHQQQHHGHPLHPHHTHQQQQQQQQQQQPPPQHHQQQQQ
QHAQQQQQQQHHAAQLAHTLSSAAVAAAAGNTGGNTNPHSIFGTGNFHYKTNNSWTLPTLTYQRIYQEN
PHYQRNSFMDPQSNAAAAAAAAAAASCNAAAVAAAAAVASGNQGSNGNASGNGNAVVAATGNGNGGNPG
QNNNNNNGNNSGNSNNNSNNNVSSV-
QHVANAVAAAVIANEHHNHLNSLKARFQPASSG-----------------
KSTSNSKEIICPDDKYK-----------------------
EEGDIWNVEAQTAFLGPNLWDKTLPYDADLKVTQYADLDEFLSENNIP--------DGLP-----
GTH------------LGHSSGLGHRSDSLG-----HAAG--
LSLGLG-----------------------HITTKRE-RSPS-PS---DC--
ISPDTLNPPSPAESTF-----SFAS-------------------------
SGRDFDPRTRAFSDEELKPQ---PMIKKSR----KQFVPD-----------ELK-
DDKYWARRRKNNIAAKRSRDARR-QKENQIAMRARYLEKENATLHQEVEQLKQE---
NMDLRAR-------
LSKFQDV-------------------------------------------------------------
--------------------------------------------------------------------
--------------------------------------------------------

        'D._melanogaster_gi_24660450_ref_NP_729300.1_'  ------------
--------------------------------------------------------------------
--------------------------------------------------------------------
--------------------------------------------------------------------
--------------------------------------------------------------------
--------------------------------------------------------------------
--------------------------------------------------MSDR-ERSS----
PTLIETGLKNLIGGRDGNY-
PLISGINGRKSPFLGFICKSNGKSTSNSKEIICPDDKYK-----------------------
EEGDIWNVEAQTAFLGPNLWDKTLPYDADLKVTQYADLDEFLSENNIP--------DGLP-----
GTH------------LGHSSGLGHRSDSLG-----HAAG--
LSLGLG-----------------------HITTKRE-RSPS-PS---DC--
ISPDTLNPPSPAESTF-----SFAS-------------------------
SGRDFDPRTRAFSDEELKPQ---PMIKKSR----KQFVPD-----------ELK-
DDKYWARRRKNNIAAKRSRDARR-QKENQIAMRARYLEKENATLHQEVEQLKQE---
NMDLRAR-------
LSKFQDV-------------------------------------------------------------
--------------------------------------------------------------------
--------------------------------------------------------
        'D._melanogaster_gi_281365800_ref_NP_729298.2_' ------------
--------------------------------------------------------------------
--------------------------------------------------------------------
--------------------------------------------------------------------
--------------------------------------------------------------------
--------------------------------------------------------------------
--------------------------------------------------MSDR-ERSS----
PTLIETGLKNLIGGRDGNY-PLISGING--------------
KSTSNSKEIICPDDKYK-----------------------
EEGDIWNVEAQTAFLGPNLWDKTLPYDADLKVTQYADLDEFLSENNIP--------DGLP-----
GTH------------LGHSSGLGHRSDSLG-----HAAG--
LSLGLG-----------------------HITTKRE-RSPS-PS---DC--
ISPDTLNPPSPAES------------------------------------------------------
---------------------N-----------ELK-DDKYWARRRKNNIAAKRSRDARR-
QKENQIAMRARYLEKENATLHQEVEQLKQE---NMDLRAR-------
LSKFQDV-------------------------------------------------------------
--------------------------------------------------------------------
--------------------------------------------------------
        'D._melanogaster_gi_24660438_ref_NP_729299.1_'  ------------
--------------------------------------------------------------------
--------------------------------------------------------------------
--------------------------------------------------------------------
--------------------------------------------------------------------
--------------------------------------------------------------------
--------------------------------------------------MSDR-ERSS----
PTLIETGLKNLIGGRDGNY-PLISGING--------------
KSTSNSKEIICPDDKYK-----------------------
EEGDIWNVEAQTAFLGPNLWDKTLPYDADLKVTQYADLDEFLSENNIP--------DGLP-----
GTH------------LGHSSGLGHRSDSLG-----HAAG--
LSLGLG-----------------------HITTKRE-RSPS-PS---DC--
ISPDTLNPPSPAESTF-----SFAS-------------------------
SGRDFDPRTRAFSDEELKPQ---PMIKKSR----KQFVPD-----------ELK-
DDKYWARRRKNNIAAKRSRDARR-QKENQIAMRARYLEKENATLHQEVEQLKQE---
NMDLRAR-------
LSKFQDV-------------------------------------------------------------

```
--------------------------------------------------------------------
--------------------------------------------------------
        'D._melanogaster_gi_442630877_ref_NP_001261546.1_'     ----
--------------------------------------------------------------------
--------------------------------------------------------------------
--------------------------------------------------------------------
--------------------------------------------------------------------
--------------------------------------------------------------------
----------------------------------------------------------MSDR-
ERSS----PTLIETGLKNLIGGRDGNY-
PLISGINGRKSPFLGFICKSNGKSTSNSKEIICPDDKYK-----------------------
EEGDIWNVEAQTAFLGPNLWDKTLPYDADLK---YADLDEFLSENNIP--------DGLP-----
GTH------------LGHSSGLGHRSDSLG-----HAAG--
LSLGLG-----------------------HITTKRE-RSPS-PS---DC--
ISPDTLNPPSPAESTF-----SFAS-------------------------
SGRDFDPRTRAFSDEELKPQ---PMIKKSR----KQFVPD-----------ELK-
DDKYWARRRKNNIAAKRSRDARR-QKENQIAMRARYLEKENATLHQEVEQLKQE---
NMDLRAR-------
LSKFQDV-------------------------------------------------------------
--------------------------------------------------------------------
--------------------------------------------------------
        'D._melanogaster_gi_442630875_ref_NP_001261545.1_'     ----
--------------------------------------------------------------------
--------------------------------------------------------------------
--------------------------------------------------------------------
--------------------------------------------------------------------
--------------------------------------------------------------------
----------------------------------------------------------MSDR-
ERSS----PTLIETGLKNLIGGRDGNY-PLISGING--------------
KSTSNSKEIICPDDKYK-----------------------
EEGDIWNVEAQTAFLGPNLWDKTLPYDADLKVTQYADLDEFLSENNIP--------DGLP-----
GTH------------LGHSSGLGHRSDSLG-----HAAG--
LSLGLG-----------------------HITTKRE-RSPS-PS---DC--
ISPDTLNPPSPAESIFDIPAFSFAS-------------------------
SGRDFDPRTRAFSDEELKPQ---PMIKKSR----KQFVPD-----------ELK-
DDKYWARRRKNNIAAKRSRDARR-QKENQIAMRARYLEKENATLHQEVEQLKQE---
NMDLRAR-------
LSKFQDV-------------------------------------------------------------
--------------------------------------------------------------------
--------------------------------------------------------
        'D._melanogaster_gi_24660469_ref_NP_729303.1_'  ------------
--------------------------------------------------------------------
MRMDYQMPPPALALQQQQQHGMQHQQQQLQLLQNVQQQQQQQQQQQQQHQQQQLLGNMSQTAALPPLS
SLPLQVVHNLPHLLAGNSGGNTGNNPNTSNSNLNVACNNNLLRGNMQHQQQQQQLLNNLSNGNNVASG
TLPPPTQLLQHHLQHLAHVNVNAAAAVVAVAANNLQQQQQQQHASS-
NNVTGGSPGSNSNNNNNNNILNHNNLNIIINNHASLN----------
NNNNNSNNNNNNNNNPATPNANLAASNNATPTVNTASAVAQQQQAHEN--
SLVNSLVGVINNSNNNNNNTNNNNNNTNS-NSNNNNNNSVGGD-DNNR------WTQFQVQQLWK-
QHANY---LNG-----------------KSTSNSKEIICPDDKYK-----------------------
EEGDIWNVEAQTAFLGPNLWDKTLPYDADLKVTQYADLDEFLSENNIP--------DGLP-----
GTH------------LGHSSGLGHRSDSLG-----HAAG--
LSLGLG-----------------------HITTKRE-RSPS-PS---DC--
ISPDTLNPPSPAESTF-----SFAS-------------------------
SGRDFDPRTRAFSDEELKPQ---PMIKKSR----KQFVPD-----------ELK-
```

DDKYWARRRKNNIAAKRSRDARR-QKENQIAMRARYLEKENATLHQEVEQLKQE---
NMDLRAR-------
LSKFQDV-------------------------------------------------------------
--------------------------------------------------------------------
--------------------------------------------------------
        'D._melanogaster_gi_442630873_ref_NP_001261544.1_'      ----
--------------------------------------------------------------------
--------
MRMDYQMPPPALALQQQQQHGMQHQQQQLQLLQNVQQQQQQQQQQQQQHQQQQLLGNMSQTAALPPLS
SLPLQVVHNLPHLLAGNSGGNTGNNPNTSNSNLNVACNNNLLRGNMQHQQQQQQLLNNLSNGNNVASG
TLPPPTQLLQHHLQHLAHVNVNAAAAVVAVAANNLQQQQQQQQHASS-
NNVTGGSPGSNSNNNNNNNNILNHNNLNIIINNHASLN----------
NNNNNSNNNNNNNNNPATPNANLAASNNATPTVNTASAVAQQQQAHEN--
SLVNSLVGVINNSNNNNNNTNNNNNNNTNS-NSNNNNNNSVGGD-DNNR------WTQFQVQQLWK-
QHANY---LNGRKS---PFLGFICKSNGKSTSNSKEIICPDDKYK-----------------------
EEGDIWNVEAQTAFLGPNLWDKTLPYDADLK---YADLDEFLSENNIP--------DGLP-----
GTH------------LGHSSGLGHRSDSLG-----HAAG--
LSLGLG-----------------------HITTKRE-RSPS-PS---DC--
ISPDTLNPPSPAESTF-----SFAS-------------------------
SGRDFDPRTRAFSDEELKPQ---PMIKKSR----KQFVPD-----------ELK-
DDKYWARRRKNNIAAKRSRDARR-QKENQIAMRARYLEKENATLHQEVEQLKQE---
NMDLRAR-------
LSKFQDV-------------------------------------------------------------
--------------------------------------------------------------------
--------------------------------------------------------
        'N._vectensis_gi_156336982_ref_XP_001619763.1_' ------------
--------------------------------------------------------------------
--------------------------------------------------------------------
--------------------------------------------------------------------
--------------------------------------------------------------------
--------------------------------------------------------------------
--------------------------------------------------------------------
--------------------------------------------------------------------
--------------------------------------------------------------------
--------------------------------------------------------------------
--------------------------------------------------------------------
------------R-KKPKD-----EYEY-NPL---PIGKKAR----RKFVPD-----------QEK-
DDRYWARRVKNNVAARRSRDMRR-QKEIEISMKWKQLEKENARLREELQQLKDR---
ASELEKK-------
LSEKQAGH------------------------------------------------------------
--------------------------------------------------------------------
--------------------------------------------------------
        'N._vectensis_gi_156400985_ref_XP_001639072.1_' ------------
--------------------------------------------------------------------
--------------------------------------------------------------------
--------------------------------------------------------------------
--------------------------------------------------------------------
--------------------------------------------------------------------
--------------------------------------------------------------------
--------------------------------------------------------------------
--------------------------------------------------------------------
--------------------------------------------------------------------
--------------------------------------------------------------------
------------------------EYEY-NPL---PIGKKAR----RKFVPD-----------QEK-

DDRYWARRVKNNVAARRSRDMRR-QKEIEISMKWKQLEKENARLREELQQLKDR---
ASELEKK-------
LSEKQAGH------------------------------------------------------------
--------------------------------------------------------------------
--------------------------------------------------------
        'A._digitifera_lcl_XM_015892501.1_prot_XP_015747987.1_1'
        ------------------------------------------------------------
--------------------------------------------------------------------
--------------------------------------------------------------------
--------------------------------------------------------------------
--------------------------------------------------------------------
--------------------------------------------------------------------
--------------------------------------------------------------------
--------------------------MSRRSSPSAPNQSDGESSDSGLDTMS-
SSTGNSFPSDYMDLDEFLSAQTHNGPGN----EAVPKASESTANQR-
VACPTRAVVLAKTNSNQINEGTSVTNETAGKANDAIMLKTSSFSLKVDS-----------
SVLQSIPSNST--AQPSATNAIIQWCRSVG---VPQQSDLK-
ALDQTSGVAP-----------------------LR-RRRSA-----KYIY-NPL---PISKKAD----
RKFVCQ-----------AEK-DEKYWERRIKNNVAAKRSRDMRR-
QKEIEISEKFKSLERENEDLKDEVQRLRVK---AVELERK-------
LANLQGGNV-----------------------------------------------------------
--------------------------------------------------------------------
--------------------------------------------------------

thyrotroph_embryonic_factor_like_1__Exaiptasia_pallida_ ------------
--------------------------------------------------------------------
--------------------------------------------------------------------
--------------------------------------------------------------------
--------------------------------------------------------------------
--------------------------------------------------------------------
--------------------------------------------------------------------
--------------------------------------------------------------------
--------------------------------------------------MNAPGSPCPSS----
TNSSLSDSEVSNDS-GIESTSGGSLP-
PDYMDLDEFLVATGVKLVVSKCDLPSRVTQRNTEVNV-----------DHENSKESNAR--
KTIDTKDNVNNSEFIID---KKKTSEEKPEKTHTEKVKN-----------------------SK-
TSQEDNDINYKYNY-NPL---PMKRKAQ----RQFVTD-----------TEK-
DERYWKRRTRNNEAARRSRDMRR-QREIEISTQCKELEKENARLKKELQKLKDK---
ANKLERQ-------
LMEK----------------------------------------------------------------
--------------------------------------------------------------------
--------------------------------------------------------

Transcription_factor_VBP__Exaiptasia_pallida_   --------------------
--------------------------------------------------------------------
--------------------------------------------------------------------
--------------------------------------------------------------------
--------------------------------------------------------------------
--------------------------------------------------------------------
--------------------------------------------------------------------
-------------------------------------------------------
MCPKDTEVLRRTAEDGGRIQQKVFKRYSRDRPIKSFDRIPIYTSILKHQVSLEKIMNAPGSPCPSS--
--TNSSLSDSEVSNDS-GIESTSGGSLP-
PDYMDLDEFLVATGVKLVVSKCDLPSRVTQRNTEVNV-----------DHENSKESNAR--

```
KTIDTKDNVNNSEFIID---KKKTSEEKPEKTHTEKVKN-----------------------SK-
TSQEDNDINYKYNY-NPL---PMKRKAQ----RQFVTD-----------TEK-
DERYWKRRTRNNEAARRSRDMRR-QREIEISTQCKELEKENARLKKELQKLKDK---
ANKLERQ-------
LMEK----------------------------------------------------------------
--------------------------------------------------------------------
--------------------------------------------------------
        'H._vulgaris_gi_221106316_ref_XP_002167159.1_'  ------------
--------------------------------------------------------------------
--------------------------------------------------------------------
--------------------------------------------------------------------
--------------------------------------------------------------------
--------------------------------------------------------------------
--------------------------------------------------------------------
--------------------------------------------------------------------
--------------------------------------------------------------------
-------------MMSKINTHYTPT-VCESSQICTASKCIKQT-YYHIPRTLK-
KTIIDN--------------NEFLISRS---LPFSETNKRYYNS--IP--
ANFKSSPPTLANSENIVIP------------------------LK-HNVNVAKSNVEYNY-KPQ---
PMIRKSK----RQFVDE-----------KKK-DETYWERRQRNNEAAKRSREQRR-
EKEIEINRKCELLEVENANLKFTVENLQEN---IQKLEDT------
VSMYKEILRRQNVS------------------------------------------------------
--------------------------------------------------------------------
---------------------------------------------------------
        'S._kowalevskii_gi_585674208_ref_XP_002736397.2_'
MGVNIIFDFLSSAQKNVSFLDDFAEKLPSVRTWQVIVVFICSVIVIQAIAIVFKLFKMKKTSEMAFKP
FATPSLDIFKSPILGHLGLLSVNKDEIMQMAGDITHEFKYALPLWLGPFQAALICHHPSTVQPILATT
EPKDDFSYGMLKPWLGDGLLISSGNKWSRNRKLLTPGFHFDILRPYVKVFNECAITMTDKWSTMCDTG
PLEMFQHISLMTLDSLLKCIFSQESHCQTDSDVNPYIKAVYTLTDLIMERINFPPYYSDTVYSLTYEG
VKWRKALNDVHNHSRRVIKERKSALKDEVERGTVNKRKYIDFLDILLAAKDEDGNGLTDKEIQDEVDT
FMFEGHDTTASGISWCFYNLARHPKYQQMCRDEVDQLLDKKENDELDWDDYAKLPFLTMCIKESLRLH
PTVPFVGRRNNKPLTFPELGITIPAGQFLGISVIGLHHNVHLWEEPLVYNPYRFTTENTKVRQNYSFL
PFSAGPRNCIGQNFAMNEMKTAIALVLRKFILSVEDDYPVRRMYNVVLRAEEGLHIVVKPREEMNPGF
GANRK----
ILVNFQFLLIGNFYPIHLKESADFYKLTDVSMFPTLNRSETVVNNTGYGDLPGEMASRPVIPTP----
-------
IYTQRITSTTRLPLPFEVPPASLIGSFVVPDYTTVRENHRIPTDTAPVIVPQ----------------
-------FRAYDEKAATFRLRRAHVRPKTF-PITRK-Q----REFIPE-----------KEK-
DEQYWEKRRKNNEAAKRSRERRR-
LSDIMVECKMAQMSDENDSMKAQLLALQHQTALLCNTDETRQAKAQSMMYLNPLYTNYNSTNPFPVPY
GMFGPPQPTFPRFTSIPPGFTEHADLLRRHNALSQRAQYHERSSRPEYSTNDVKQRESDTERKDDTTD
CTSSVACATQSVHGYTE---------------------------------------------------
-----------------------------------
        'A._digitifera_lcl_XM_015892500.1_prot_XP_015747986.1_1'
        ------------------------------------------------------------
--------------------------------------------------------------------
--------------------------------------------------------------------
--------------------------------------------------------------------
--------------------------------------------------------------------
--------------------------------------------------------------------
MSIHAMESLSSARPEKFHDCNSFTFQDSDDSGLSSSPSSPTPVGVDFLGDLFDLDTNFSNLLNTSLFD
GEFPSKLYTHIKIKDDDDEDMWFNASSSKVNTKTSSLSAGDSSTGQDYFDDSFELIAELLN-
PQAEVEEKMTGNSSKGKN-----LTIQQ----SAE------------
ISEVESKNNEACTPTSESGENSSNNDTISAINSPDTSV---------------
```

```
KRELRTRRKRVVVNTDH-DYCR--
QRLVSDDEPDSDDDYK---------------------------------
FVEDIEKEESDDSDDEDFKAP-C-P-AKKVR----KA-SAK-----------GVK-
DSKYWERRQRNNLAAKRSREAKR-AREIQVAKKTVALEKENANLKKQVRKLKAN---
IKRAEKM-------
LRLMV---------------------------------------------------------------
--------------------------------------------------------------------
--------------------------------------------------------
        'N._vectensis_gi_156408099_ref_XP_001641694.1_' ------------
--------------------------------------------------------------------
--------------------------------------------------------------------
--------------------------------------------------------------------
--------------------------------------------------------------------
--------------------------------------------------------------------
--------------------------------------------------------------------
--------------------------------------------------------------------
--------------------------------------------------------------------
--------------------------------------------------------------------
--------------------------------------------------------------------
-----------------------------------------------A-YGR-----------D-K-
DQKYIEKRMKNNLAAKRSREAKR-QREIEAMQKTLTLEKENSDLNKEVNKLKKM---
IARLENK-------
LR------------------------------------------------------------------
--------------------------------------------------------------------
--------------------------------------------------------
        'A._queenslandica_gi_340372715_ref_XP_003384889.1_'      ----
--------------------------------------------------------------------
--------------------------------------------------------------------
--------------------------------------------------------------------
--------------------------------------------------------------------
--------------------------------------------------------------------
--------------------------------------------------------------------
--------------------------------------------------------------------
------------------------------------------------------
MDDSSSKNLCVGSNG----LFLNG----QED------------LGLFASFQHQWSN--
MAGGASSSS--SLAAMAGLQN-----------------DKSLGTTGSSSSPEDAS-FLNS--
HAMIS--AVDSNRS-----------------------------------------
SAISDHTDTEDYEDD-R-PKRKRPR----NP-IPN-----------EKK-
DDKYWERRRKNNIAAKRSRETKK-QKVDEELLKAKDAIQENHKLKQEIEVLKAE---
INSLRRL-------
LKDANMTLSLWIRAKQTSEPASQLPPMLRNTTGNMPFVTFPVTTSNL---------------------
--------------------------------------------------------------------
--------------------------------------------------------

hepatic_leukemia_factor_like__Exaiptasia_pallida_       ------------
--------------------------------------------------------------------
--------------------------------------------------------------------
--------------------------------------------------------------------
--------------------------------------------------------------------
--------------------------------------------------------------------
--------------------------------------------------------------------
--------------------------------------------------------------------
-----------------------------M-EDDEVCPSP-VKRERLNSEVVADHQRVCV-----
IQRP---KQRI------------ELMSDD--EQNSAYYCKRGPVLSEN-SINNIL-
```

```
KVVSLQS------------FSSSR-AQ-Q--SNN-EVN--SHRSELGSDSPLSSIRSSGESS----
PSSED------------------------SNPQFTP-IPEIERNA---SR--KGSSR-R----RGE-
K---------VASSMK-DDKYWEKRLRNNASAKRSRDARR-
VRELECQIRSEFMEEENRKLEEENKMLREE---NARLSK-------
MIQELKSRA-----------------------------------------------------------
--------------------------------------------------------------------
---------------------------------------------------------
        'N._vectensis_gi_156388041_ref_XP_001634510.1_' ------------
--------------------------------------------------------------------
--------------------------------------------------------------------
--------------------------------------------------------------------
--------------------------------------------------------------------
--------------------------------------------------------------------
--------------------------------------------------------------------
--------------------------------------------------------------------
-----------------------------MSEETKECKRRCVKRSK--SETESEDEDSYP-----
TER----RQQIN-----------EGESED--EIRSVYCCRSTPHYGETTSLNNIL-
KVVSLQS------------FSNSRLAS-DPGSNNRHFDS-SAREQVQRNSVIISAPSKTKCE----
PEKSD------------------------REKE-TPQINGIYTPTVF-NT--KSSSRGK----RTE-
K---------HFNNNK-DNKYWEKRQRNNASAKRSRDARR-
VRELECQIRAEFLEEENHKYKVENEMLREE---NERLLK-------
IIESFNNKQ-----------------------------------------------------------
--------------------------------------------------------------------
---------------------------------------------------------
        'A._digitifera_lcl_XM_015913431.1_prot_XP_015768917.1_1'
        ------------------------------------------------------------
--------------------------------------------------------------------
--------------------------------------------------------------------
--------------------------------------------------------------------
--------------------------------------------------------------------
--------------------------------------------------------------------
--------------------------------------------------------------------
-------------------------------------------------MEEEDVRAQKRTRKRSR--
SDSSDDSISGCSR----RDLE---KHATN-----------TSLTLI--EDPGVY-GRRIPYFGEN-
SINDIL-KVVSLQS------------
FSNPRLSSLNEGERDEMIHQEDHKRAKTTESDCQSDSSEELSR----
KERSS------------------------SSSE-DA-VPSSGYSQV--NF--PSKSRGK----
RANFKNGSITN--AIVGNGK-DQRYWEKRQRNNASAKRSRDARR-
VRELETQIRAEYLEDENYRLKGENERLREE---NVRLQQ-------
AMERLKENGVSSAKDCDD--------------------------------------------------
--------------------------------------------------------------------
---------------------------------------------------------
        'T._adherens_lcl_XM_002117855.1_prot_XP_002117891.1_1'  ----
--------------------------------------------------------------------
--------------------------------------------------------------------
--------------------------------------------------------------------
--------------------------------------------------------------------
--------------------------------------------------------------------
--------------------------------------------------------------------
----
MQSEKADFGELEHYLKPVESIFTGDELLEDTSEFIINNIENKRFQVGQQQRQNHQQRQQLQHQVNYDD
ATRLWMWNNHTTRGRNEHKSQNMINTTTNYNHFAVDSNSKSPYSNLNYSSFTPYHHSLDLSGRSP---
-LQVTTVLSNTSN-----------
HLLQQQQHQQHQQQPSTQQPNHHSTVTLTNSKDKDVSLANN----------
```

HHFKTKITTSTFTLSEAESDLLSAQPIQQQQSMQEDKSDDTLLT----
AYFSH------------------------DHDKHSAMIQHASPATTDENS--RDSPELK----RAK-
KT-------LVPDNCK-DTKYWARRIKNNKAAKRSRDARK-
ERETRITQECQMLAEANLKLKSEIVTLKKLL--NDALAKQN-----
VLQEQQQHQQR---------------------------------------------------------
--------------------------------------------------------------------
---------------------------------------------------------
        'A._californica_gi_524907652_ref_XP_005108942.1_'       ----
--------------------------------------------------------------------
--------------------------------------------------------------------
--------------------------------------------------------------------
--------------------------------
MYWERRRKNNESARKSREKRRVHDMALEGRICSLEDDNTKLRRELLALKKKFNLPESQPVIVEEESEE
KASPPSRRVNVSPPRTKNQGQSAMLHSPVRGMSSKPSGPSHMAYNSPPPLLAVAGAMPMGVAMYPDSS
PGLPYFMSESSRAGQEGVKSVMPSGLSHHRHQLPSRPQQQQQHHHHHHHQLPHPPLPVDVLKHEPMEE
GEVR---
TRERSNSSVERRPSLHLDLPDTPMLRRGSVGSFPPYPHPSLSPHPVPSTQVSHHHHHHHHATPHTYPN
PHLNPASGSSNPSHFFPRSYESYAPPPRP---VRSPVS---SHSSDDNYDEPL--
QLTVRRDSAVSNQSNQQHDDSSRESEPGCVMGGDNKPASVSPP---------
ASAFPLKLRHKLPSHELAYPKEMFPSMAATAAVSAAQPSFPTHPFMNGLAHLS---------------
----------DMALSQTNPLSLVKSENPPAPS-HYGRQRKESSSRSALRRN-------
SNETKYMDPKYLERRRRNNEAARKCRENRKALTKLREAKS-DYLESENSKLRDELTSLQEE---
MKQLRELIE-----
KKRLEQGIKEEPHAES----------------------------------------------------
--------------------------------------------------------------------
--------------------------------------------------------
        'H._vulgaris_gi_221116695_ref_XP_002159994.1_'  ------------
--------------------------------------------------------------------
--------------------------------------------------------------------
--------------------------------------------------------------------
--------------------------------------------------------------------
--------------------------------------------------------------------
--------------------------------------------------------------------
--------------------------------------------------------------------
-----------------------MSPEE-ENNWMN-GQGRMGG---MEDEIDDF---------
LKILMD---EHKSD---------QIVLDQNTKNTD-MHFLCTNSLLGHKP-
LQLTSNNCDGAESP----------ASSYG-TL-YSPPSSPESIRIQMNSS--
CNKNISINEP----------VVHIP-------------------------SIMLGQDNDGS-YKYD--
PRP---LGRKKR----RNLVP----------DEKKTID--YWERRKRNNQAARRSREERR-
LKELQTFHCMKLLQNENTSLKQTIQIMLRN---QQQLQEEINMM---KQVIDQSYKQR-
HGT-----------------------------------------------------------------
--------------------------------------------------------------------
--------------------------------------------

A0A1D5NSQ7_DANRE        --------------------------------------------
--------------------------------------------------------------------
--------------------------------------------------------------------
--------------------------------------------------------------------
--------------------------------------------------------------------
--------------------------------------------------------------------
--------------------------------------------------------------------
--------------------------------------------------------------------
----------------------------------------------------------
MSVSDNIFSVHEASSDSSAQTPMDSALYTQT--ISFTK---------------------

```
SPEVMMGY---LPYSSCLNNPNT--NTERSVQ-QSSAHTQD------------------------
FAQFLEPPPASALRLCAQ---------------------KRG-----------
VSKDSAEYRQRRERNNIAVRKSRDKAR-RRIQMTQQRALQLQDENQRLQVHIQRLLHE---
VEALRHY-------LSQRH-----------LQDTSE--
EH------------------------------------------------------------------
--------------------------------------------------------------------
--------------------------------
        'D._rerio_lcl_NM_131837.1_prot_NP_571912.1_1'   ------------
--------------------------------------------------------------------
--------------------------------------------------------------------
--------------------------------------------------------------------
--------------------------------------------------------------------
--------------------------------------------------------------------
--------------------------------------------------------------------
--------------------------------------------------------------------
--------------------------------------------------------------------
----------------------MSVSDNIFSVHEASSDSSAQTPMDSALYTQT--
ISFTK---------------------SPEVMMGY---LPYSSCLNNPNT--NTERSVQ-
QSSAHTQD------------------------
FAQFLEPPPASALRLCAQ---------------------KRG-----------
VSKDSAEYRQRRERNNIAVRKSRDKAR-RRIQMTQQRALQLQDENHRLQVHIQRLLHE---
VEALRHY-------LSQRH-----------LQDTSE--
EH------------------------------------------------------------------
--------------------------------------------------------------------
--------------------------------
        'CCAAT_enhancer_binding_protein_delta_OS_Homo_sapiens_OX_960
6_GN_CEBPD_PE_1_SV_2'   --------------------------------------------
--------------------------------------------------------------------
--------------------------------------------------------------------
--------------------------------------------------------------------
--------------------------------------------------------------------
--------------------------------------------------------------------
---------------------------------------------MSAALFS-----------
LDGPARG-------------------APWPAEP-APFYEPGRA-GKPG----RG--AEP-
GALGEPG----------------AAAPAMYDDESAIDFSAYIDS--MAAVPTLELCHDELFADL--
FNSNHKAGGAG--PLELLPGGPARPLGP--GPA-AP--RLLKR---------
EPDWGDGDAPGSLLPAQVAA---CAQT-VVSLAAAGQPTPPTSP-EPPRSS-
P-------------------------RQT--PAPGPAREKSA---------------------
GKRG-----------PDRGSPEYRQRRERNNIAVRKSRDKAK-
RRNQEMQQKLVELSAENEKLHQRVEQLTRD---LAGLRQF-------FKQLP---SPPF----LP-
AAGTADCR------------------------------------------------------------
--------------------------------------------------------------------
-------------------------------------
        'M._musculus_gi_110347410_ref_NP_031705.3_'     ------------
--------------------------------------------------------------------
--------------------------------------------------------------------
--------------------------------------------------------------------
--------------------------------------------------------------------
--------------------------------------------------------------------
--------------------------------------------------------------------
---------MSAALFS-----------LDSPVRG-------------------TPWPTEP-
AAFYEPGRV-DKPG----RG--PEP-GDLGELG----------------
STTPAMYDDESAIDFSAYIDS--MAAVPTLELCHDELFADL--FNSNHKAAGAG--
GLELLQGGPTRPPGV--GSV-AR--GPLKR---------EPDWGDGDAPGSLLPAQVAV---CAQT-
```

VVSLAAAAQPTPPTSP-EPPRGS-P-------------------------GPS--
LAPGTVREKGA---------------------GKRG-----------
PDRGSPEYRQRRERNNIAVRKSRDKAK-RRNQEMQQKLVELSAENEKLHQRVEQLTRD---
LAGLRQF-------FKKLP---SPPF----LP-PTG-
ADCR----------------------------------------------------------------
--------------------------------------------------------------------
---------------------------------
        'X._laevis_lcl_NM_001089607.2_prot_NP_001083076.1_1'    ----
--------------------------------------------------------------------
--------------------------------------------------------------------
--------------------------------------------------------------------
--------------------------------------------------------------------
--------------------------------------------------------------------
--------------------------------------------------------------------
-----------------MSSVSMS-----------LEARCLSP-----------------
YAAWYMEP-TNFYEQRLS-GSPALCKPRGLCEEPETVVGTGGTLAEL-----------
SAAPAMYDDESAIDFSSYIDS--MASVPNLELCNDELFADL--FNSS-
KAAGERQEGDYLMSGLLAAPHCPP-GP--AK--VQLKR---------EPEWSDR---
SSSLPNQIAA---CAQT-SMSL----QPTPPTSP-EPSTSACP-------------------------
SPA--DSSASCGKDRT---------------------GKKC-----------
LDRYSPEYRQRRERNNIAVRKSRDKAK-RRNTDMQQKMLELSSENEKLHKKIELLTRD---
LSSLRHY-------FKQLPPSTTSSF----LPSLTG-
IDCR----------------------------------------------------------------
--------------------------------------------------------------------
---------------------------------
        'X._laevis_lcl_NM_001089609.1_prot_NP_001083078.1_1'    ----
--------------------------------------------------------------------
--------------------------------------------------------------------
--------------------------------------------------------------------
--------------------------------------------------------------------
--------------------------------------------------------------------
--------------------------------------------------------------------
----------------------MS-----------LEARCLSP-----------------
YAAWYMEP-TNFYEQRLS-GSPAPYKPRGMCEEPEAAVGTGGTLVEL-----------
SAAPAMYDDESAIDFSSYIDS--MASVPNLELCNDELFADL--FNSS-
KAVGERQEGDYLMSGLLAAPHCPP-GP--AK--VQLKQ---------EPEWSDSDM-
SSSLPNQIAA---CAQT-SMSL----QPTPPTSP-EPSTSACP-------------------------
SPA--ASSGSCGKDRS---------------------GKKC-----------
TDRYSPEYRQRRERNNIAVRKSRDKAK-RRNVDMQQRLLELSSENEKLHKKIELLTRD---
LSSLRHF-------FKQLPPAATGPF----LPSLTG-
IDCR----------------------------------------------------------------
--------------------------------------------------------------------
---------------------------------
        'D._rerio_lcl_NM_131887.1_prot_NP_571962.1_1'   ------------
--------------------------------------------------------------------
--------------------------------------------------------------------
--------------------------------------------------------------------
--------------------------------------------------------------------
--------------------------------------------------------------------
--------------------------------------------------------------------
-------MVTSAASMSDM-------YNLDSQCVTPP---------------CNMSWAMEP-
ANFYDNKGD-AKPG--------ENNNNNSSSTGNMMEL-----------
SNAPAIYDDESAIDFSAYIES--MSTVP-LEICNDELFADL--
FNNTVKQEKPDFYMSNTFAHKSAERHLEGFGKGSFC--APIKK---------EADWSDSEH-

```
SSSLPSQIEA---CAQT-SVNFMHTGQPTPPTTP-EPE----P-------------------------
VAH--RRPG--KEK-----------------------GKKN-----------
VDRHSPEYRQRRERNNIAVRKSRDKAK-QRNLDMQQKMIELGAENERLHKTIDQLTRE---
LSSLRNF-------FKQMP---EASF----
GSAARAAVDSR---------------------------------------------------------
--------------------------------------------------------------------
----------------------------------------
        'CCAAT_enhancer_binding_protein_epsilon_OS_Homo_sapiens_OX_9
606_GN_CEBPE_PE_1_SV_2' --------------------------------------------
--------------------------------------------------------------------
--------------------------------------------------------------------
--------------------------------------------------------------------
--------------------------------------------------------------------
--------------------------------------------------------------------
---------------------------------------------MSHGTYYECEPRGGQ----
QPLEFS------GG---------RAGP-GEL--GDMCEHEASIDLSAYIESG--
EEQLLSDLFAVKPAPEARG-------LKGPGTPAFPH--YLPPDP------
RPFAYPPHTFGPDRKALG---PGIYSSPGSYDPRAVAVKEEPRGPEGSRAASRG-----
SYNP----------------------LQYQVAH---CGQT-AMHLPP---TLAAPGQ-
PLRVLKAP-------------------------LAT-AAPPCSPLLKAPSP------AG-------
PLHKGKKA-----------VNKDSLEYRLRRERNNIAVRKSRDKAK-
RRILETQQKVLEYMAENERLRSRVEQLTQE---LDTLRNL-------FRQIP---EAAN----
LIKGVG--
GCS-----------------------------------------------------------------
--------------------------------------------------------------------
--------------------------------
        'M._musculus_gi_46369479_ref_NP_997014.1_'      ------------
--------------------------------------------------------------------
--------------------------------------------------------------------
--------------------------------------------------------------------
--------------------------------------------------------------------
--------------------------------------------------------------------
--------------------------------------------------------------------
---------MSHGTYYECEPRGGQ----QPLEFS------GG---------RAGP-GEL--
GDMCEHEASIDLSAYIESG--EEQLLSDLFAMKPTPEARS-------LKGPGAPSFPH--
YLPADP------RPFAYPSHTFGPDRKALG---
PGIYSNPGSYDPRAVAVKEEPRGPEGNRGTSRG-----SYNP----------------------
LQYQVAH---CGQT-AVHLPP---TLAAPGQ-PLRVLKAP-------------------------
VAA-AAPPCSPLLKAPSP------AG-------PSHKGKKA-----------
VNKDSLEYRLRRERNNIAVRKSRDKAK-RRIMETQQKVLEYMAENERLRNRVDQLTQE---
LDTLRNL-------FRQIP---EAAS----LIKGVG--
GCS-----------------------------------------------------------------
--------------------------------------------------------------------
--------------------------------
        'X._laevis_lcl_XM_018243908.1_prot_XP_018099397.1_1'    ----
--------------------------------------------------------------------
--------------------------------------------------------------------
--------------------------------------------------------------------
--------------------------------------------------------------------
--------------------------------------------------------------------
--------------------------------------------------------------------
-----------------MSHGSYYEYEKRG------QGSAYA------AR---------
LSGPHSELVGGNLCDPETSVDLSSYMDTG--EE-ILSDLFPLK---QDR--------LKG----
TYP---YMPPEG------LPSA-AMYGYAPS-NA-----P--ERRMGGYESGGVIVKEETRG------
```

```
THRS-----VCNT----------------------LQYQAAQ---CAQT-AMHLPS---PLEGVHP-
ALRVLKGS-------------------------ISG-MLS-
SPPLKDMAS-------------------KGKKC-----------
LSKDSLEYRLRRERNNIAVRKSRDKTK-RRNLETQQRALEYMTENEKLRNRVQQLTQE---
LDALRGV-------FRQIP---EAAA----LSKGSG--
GCS-----------------------------------------------------------------
--------------------------------------------------------------------
--------------------------------
        'X._laevis_lcl_XM_018259384.1_prot_XP_018114873.1_1'    ----
--------------------------------------------------------------------
--------------------------------------------------------------------
--------------------------------------------------------------------
--------------------------------------------------------------------
--------------------------------------------------------------------
--------------------------------------------------------------------
-----------------MSHSSYYEYEKRG------QSSAYT------AR---------
LSGPHSELVGGNLCDPETSVDLSSYMDTG--EE-ILSDLFPLK---QER--------LKG----
TYP---YIPPEG------LPSA-ALYGYAPS-NA-----P--ERRMGGYESGGVIVKEETRG------
IHRG-----VCNT----------------------LQYQVAQ---CAQT-AMHLTS---PLENVHP-
ALRVLKGS-------------------------ISG-MLS-
GSPLKDTAS-------------------KGKKG-----------
LSKDSLEYRVRRERNNIAVRKSRDKAK-RRNLETQQRALGYMAENEKLRNRVQQLTQE---
LDALRGV-------FRQIP---EAAA----MSKGPG--
GCS-----------------------------------------------------------------
--------------------------------------------------------------------
--------------------------------
        'X._laevis_lcl_NM_001086806.1_prot_NP_001080275.1_1'    ----
--------------------------------------------------------------------
--------------------------------------------------------------------
--------------------------------------------------------------------
--------------------------------------------------------------------
--------------------------------------------------------------------
--------------------------------------------------------------------
-----------------MEQANFYEVDPRPSMNIHVQ-PPHG----AYGY---------
REPPASALEHNELCENENSIDISAYIDPAAFNDEFLADLFH-SNKQEK-----------
AKGDFEYPQQQQQGPVGAAV---T-GHPLMYGC-MANYMDS-KLD--------
SGLRPLAIKQEPREEEAAASRASS-L--AALYP---------H-HAASQHSS--HLQYQVAH---
CAQT-TMHLQSG-HPTPPPTP-VPSPHHHPAH----------------------HHHHHLQT--
SSLKGISPSSSTSSSSS------ESRGKSKKW-----------
VDKNSNEYRVRRERNNIAVRKSRDKAK-MRNVETQQKVFELSSDNDKLRKRVEQLSRE---
LETLRGI-------FRQLP---ESS-----LVKAMG--
NCA-----------------------------------------------------------------
--------------------------------------------------------------------
--------------------------------
        'X._laevis_lcl_NM_001091687.1_prot_NP_001085156.1_1'    ----
--------------------------------------------------------------------
--------------------------------------------------------------------
--------------------------------------------------------------------
--------------------------------------------------------------------
--------------------------------------------------------------------
--------------------------------------------------------------------
-----------------MEQANFYEVDPRPSMNIHVQ-PPHG----VYGY---------
REPPASALEHNELCDNENSIDISAYIDPAAFNDEFLADLFH-SNKQEK-----------AKSDFEY-
QQQQQGPVGAVV---AAGHPLMYGCSMANYMDN-KLD--------
```

GGLRPLVIKQEPREVEEAGGRASS-L--AALYP---------H-HAASQHSS--HLQYQVAH---
CAQT-SMHLQPG-HPTPPPTP-VPSPHHPPAH----------------------QHHH-LHH--
SSLKGISPSSSSSSSSS----SSENRGKSKKW-----------
LDKNSNEYRVRRERNNIAVRKSRDKAK-IRNVETQQKVIELSSDNDKLRKRVEQLSRE---
LDTLRGI-------FRQLP---DSS-----LVKAMG--
NCS-----------------------------------------------------------------
--------------------------------------------------------------------
--------------------------------
'D._rerio_lcl_NM_131885.2_prot_NP_571960.1_1'   ------------
--------------------------------------------------------------------
--------------------------------------------------------------------
--------------------------------------------------------------------
--------------------------------------------------------------------
--------------------------------------------------------------------
--------------------------------------------------------------------
---------MEQANLYEVAPRPLMTSLVQ-NQQN----PYIY---------KDT-AGDLS--
EICENENSIDISAYIDPSAFNDEFLADLFHNSSKQEKLKL--------ASGDYDY----
HHGANG------APGAPQMYGC-LNGYMDSSKLEPIYD--SQARMRPVAIKQEPREEDELGDSMPP-
T--YHHSQ---------H-HAP--HLS--YLQHQIAH---CAQT-TMHLQPG-HPTPPPTP-
VPSPHH--------------------------QHSH-LPG--GSMK-IG-----------------
DRGKSKKH-----------VDKNSTEYRLRRERNNIAVRKSRDKAK-
MRNVETQQKVIELSADNDRLRKRVEHLTRE---LETLRGI-------FRQLP---DGS-----
FVKAMG--
NCA-----------------------------------------------------------------
--------------------------------------------------------------------
--------------------------------
'H._sapiens_gi_28872794_ref_NP_004355.2_'       ------------
--------------------------------------------------------------------
--------------------------------------------------------------------
--------------------------------------------------------------------
--------------------------------------------------------------------
--------------------------------------------------------------------
--------------------------------------------------------------------
---------
MESADFYEAEPRPPMSSHLQSPPHAPSSAAFGFPRGAGPAQPPAPPAAPEPLGGICEHETSIDISAYI
DPAAFNDEFLADLFQHSRQQEKAKAAVGPTGGGGGGDFDYPG-
APAGPGGAVMPGGAHGPPPGYGCAAAGYLDG-
RLEPLYERVGAPALRPLVIKQEPREEDEAKQLALAGL--
FPYQPPPPPPPPSHPHPHPPPAHLAAPHLQFQIAH---CGQT-TMHLQPG-HPTPPPTP-
VPSPHPAPAL----------------------GAAG-LPGPGSALKGLGAAHPDLRASG---GS--
GAGKAKKS-----------VDKNSNEYRVRRERNNIAVRKSRDKAK-
QRNVETQQKVLELTSDNDRLRKRVEQLSRE---LDTLRGI-------FRQLP---ESS-----
LVKAMG--
NCA-----------------------------------------------------------------
--------------------------------------------------------------------
--------------------------------
'H._sapiens_gi_566559994_ref_NP_001274364.1_'   ------------
--------------------------------------------------------------------
--------------------------------------------------------------------
--------------------------------------------------------------------
--------------------------------------------------------------------
--------------------------------------------------------------------
--------------------------------------------------------------------
-----------------------

```
MSSHLQSPPHAPSSAAFGFPRGAGPAQPPAPPAAPEPLGGICEHETSIDISAYIDPAAFNDEFLADLF
QHSRQQEKAKAAVGPTGGGGGGDFDYPG-APAGPGGAVMPGGAHGPPPGYGCAAAGYLDG-
RLEPLYERVGAPALRPLVIKQEPREEDEAKQLALAGL--
FPYQPPPPPPPSHPHPHPPPAHLAAPHLQFQIAH---CGQT-TMHLQPG-HPTPPPTP-
VPSPHPAPAL----------------------GAAG-LPGPGSALKGLGAAHPDLRASG---GS--
GAGKAKKS-----------VDKNSNEYRVRRERNNIAVRKSRDKAK-
QRNVETQQKVLELTSDNDRLRKRVEQLSRE---LDTLRGI-------FRQLP---ESS-----
LVKAMG--
NCA-----------------------------------------------------------------
--------------------------------------------------------------------
--------------------------------
        'H._sapiens_gi_566559992_ref_NP_001274353.1_'   ------------
--------------------------------------------------------------------
--------------------------------------------------------------------
--------------------------------------------------------------------
--------------------------------------------------------------------
--------------------------------------------------------------------
------------------------------------------
MRGRGRAGSPGGRRRRPAQAGGRRGSPCRENSNSPMESADFYEAEPRPPMSSHLQSPPHAPSSAAFGF
PRGAGPAQPPAPPAAPEPLGGICEHETSIDISAYIDPAAFNDEFLADLFQHSRQQEKAKAAVGPTGGG
GGGDFDYPG-APAGPGGAVMPGGAHGPPPGYGCAAAGYLDG-
RLEPLYERVGAPALRPLVIKQEPREEDEAKQLALAGL--
FPYQPPPPPPPSHPHPHPPPAHLAAPHLQFQIAH---CGQT-TMHLQPG-HPTPPPTP-
VPSPHPAPAL----------------------GAAG-LPGPGSALKGLGAAHPDLRASG---GS--
GAGKAKKS-----------VDKNSNEYRVRRERNNIAVRKSRDKAK-
QRNVETQQKVLELTSDNDRLRKRVEQLSRE---LDTLRGI-------FRQLP---ESS-----
LVKAMG--
NCA-----------------------------------------------------------------
--------------------------------------------------------------------
--------------------------------
        'H._sapiens_gi_551894998_ref_NP_001272758.1_'   ------------
--------------------------------------------------------------------
--------------------------------------------------------------------
--------------------------------------------------------------------
--------------------------------------------------------------------
--------------------------------------------------------------------
--------------------------------------------------------------------
--------------------------------------------------------------------
-------------------------------------------------------------
MPGGAHGPPPGYGCAAAGYLDG-RLEPLYERVGAPALRPLVIKQEPREEDEAKQLALAGL--
FPYQPPPPPPPSHPHPHPPPAHLAAPHLQFQIAH---CGQT-TMHLQPG-HPTPPPTP-
VPSPHPAPAL----------------------GAAG-LPGPGSALKGLGAAHPDLRASG---GS--
GAGKAKKS-----------VDKNSNEYRVRRERNNIAVRKSRDKAK-
QRNVETQQKVLELTSDNDRLRKRVEQLSRE---LDTLRGI-------FRQLP---ESS-----
LVKAMG--
NCA-----------------------------------------------------------------
--------------------------------------------------------------------
--------------------------------
        'M._musculus_gi_566559971_ref_NP_001274444.1_'  ------------
--------------------------------------------------------------------
--------------------------------------------------------------------
--------------------------------------------------------------------
--------------------------------------------------------------------
--------------------------------------------------------------------
```

--------------------------------------------------------------------
-----------------------
MSSHLQSPPHAPSNAAFGFPRGAGPAPPPAPPAAPEPLGGICEHETSIDISAYIDPAAFNDEFLADLF
QHSRQQEKAKAAAGP--AGGGGDFDYPG-APAGPGGAVMSAGAHGPPPGYGCAAAGYLDG-
RLEPLYERVGAPALRPLVIKQEPREEDEAKQLALAGL--FPYQPPPPPPP--
PHPHASPAHLAAPHLQFQIAH---CGQT-TMHLQPG-HPTPPPTP-
VPSPHAAPAL----------------------GAAG-
LPGPGSALKGLAGAHPDLRTGGGGGGSGAGAGKAKKS-----------
VDKNSNEYRVRRERNNIAVRKSRDKAK-QRNVETQQKVLELTSDNDRLRKRVEQLSRE---
LDTLRGI-------FRQLP---ESS-----LVKAMG--
NCA-----------------------------------------------------------------
--------------------------------------------------------------------
--------------------------------
'M._musculus_gi_566559986_ref_NP_001274450.1_' ------------
--------------------------------------------------------------------
--------------------------------------------------------------------
--------------------------------------------------------------------
--------------------------------------------------------------------
--------------------------------------------------------------------
--------------------------------------------------------------------
--------------------------------------------------------------------
-------------------------------------------------------------
MSAGAHGPPPGYGCAAAGYLDG-RLEPLYERVGAPALRPLVIKQEPREEDEAKQLALAGL--
FPYQPPPPPPP--PHPHASPAHLAAPHLQFQIAH---CGQT-TMHLQPG-HPTPPPTP-
VPSPHAAPAL----------------------GAAG-
LPGPGSALKGLAGAHPDLRTGGGGGGSGAGAGKAKKS-----------
VDKNSNEYRVRRERNNIAVRKSRDKAK-QRNVETQQKVLELTSDNDRLRKRVEQLSRE---
LDTLRGI-------FRQLP---ESS-----LVKAMG--
NCA-----------------------------------------------------------------
--------------------------------------------------------------------
--------------------------------
'M._musculus_gi_86198301_ref_NP_031704.2_' ------------
--------------------------------------------------------------------
--------------------------------------------------------------------
--------------------------------------------------------------------
--------------------------------------------------------------------
--------------------------------------------------------------------
--------------------------------------------------------------------
---------
MESADFYEVEPRPPMSSHLQSPPHAPSNAAFGFPRGAGPAPPPAPPAAPEPLGGICEHETSIDISAYI
DPAAFNDEFLADLFQHSRQQEKAKAAAGP--AGGGGDFDYPG-
APAGPGGAVMSAGAHGPPPGYGCAAAGYLDG-
RLEPLYERVGAPALRPLVIKQEPREEDEAKQLALAGL--FPYQPPPPPPP--
PHPHASPAHLAAPHLQFQIAH---CGQT-TMHLQPG-HPTPPPTP-
VPSPHAAPAL----------------------GAAG-
LPGPGSALKGLAGAHPDLRTGGGGGGSGAGAGKAKKS-----------
VDKNSNEYRVRRERNNIAVRKSRDKAK-QRNVETQQKVLELTSDNDRLRKRVEQLSRE---
LDTLRGI-------FRQLP---ESS-----LVKAMG--
NCA-----------------------------------------------------------------
--------------------------------------------------------------------
--------------------------------
'M._musculus_gi_566559969_ref_NP_001274443.1_' ------------
--------------------------------------------------------------------
--------------------------------------------------------------------

```
--------------------------------------------------------------------
--------------------------------------------------------------------
--------------------------------------------------------------------
----------------------------------------
MRGREPVGALGGRRRQRRHAQAGGRRGSPCRENSNSPMESADFYEVEPRPPMSSHLQSPPHAPSNAAF
GFPRGAGPAPPPAPPAAPEPLGGICEHETSIDISAYIDPAAFNDEFLADLFQHSRQQEKAKAAAGP--
AGGGGDFDYPG-APAGPGGAVMSAGAHGPPPGYGCAAAGYLDG-
RLEPLYERVGAPALRPLVIKQEPREEDEAKQLALAGL--FPYQPPPPPPP--
PHPHASPAHLAAPHLQFQIAH---CGQT-TMHLQPG-HPTPPPTP-
VPSPHAAPAL----------------------GAAG-
LPGPGSALKGLAGAHPDLRTGGGGGGSGAGAGKAKKS-----------
VDKNSNEYRVRRERNNIAVRKSRDKAK-QRNVETQQKVLELTSDNDRLRKRVEQLSRE---
LDTLRGI-------FRQLP---ESS-----LVKAMG--
NCA-----------------------------------------------------------------
--------------------------------------------------------------------
--------------------------------
        'M._musculus_gi_567316233_ref_NP_001274452.1_'  ------------
--------------------------------------------------------------------
--------------------------------------------------------------------
--------------------------------------------------------------------
--------------------------------------------------------------------
--------------------------------------------------------------------
--------------------------------------------------------------------
--------------------------------------------------------------------
--------------------------------------------------------------------
--------------------------------------------------------------------
------------------------------MHLQPG-HPTPPPTP-
VPSPHAAPAL----------------------GAAG-
LPGPGSALKGLAGAHPDLRTGGGGGGSGAGAGKAKKS-----------
VDKNSNEYRVRRERNNIAVRKSRDKAK-QRNVETQQKVLELTSDNDRLRKRVEQLSRE---
LDTLRGI-------FRQLP---ESS-----LVKAMG--
NCA-----------------------------------------------------------------
--------------------------------------------------------------------
--------------------------------
        'M._musculus_gi_6753404_ref_NP_034013.1_'       ------------
--------------------------------------------------------------------
--------------------------------------------------------------------
--------------------------------------------------------------------
--------------------------------------------------------------------
--------------------------------------------------------------------
---------------------------------------------------------------
MHRLLAWD----AACLPPPPAAFRPMEVANFYYEPDCLA-YGAKAARA------APR-APAAEP---
AIGEHERAIDFSPYLEPLAPAADFAAPAPAHH---------------DFLSDLF-
ADDYGAKPSKKPADYG-YVSLGRAGAKAAPPACF-PPPPPAALKAEPGFEP--ADCK-RADDA-
PAMAAGFP--FALRA---------------------YLGYQATP---SGSSGSLST-SSS-
SSPPGTP-SPADAKAA-------------------------PAACFAGPPAAP--
AKA-------------------KAKKT-----------VDKLSDEYKMRRERNNIAVRKSRDKAK-
MRNLETQHKVLELTAENERLQKKVEQLSRE---LSTLRNL-------FKQLP----EP-----
LLASAG--
HC------------------------------------------------------------------
--------------------------------------------------------------------
--------------------------------
        'M._musculus_gi_567757569_ref_NP_001274667.1_'  ------------
--------------------------------------------------------------------
```

--------------------------------------------------------------------
--------------------------------------------------------------------
--------------------------------------------------------------------
--------------------------------------------------------------------
--------------------------------------------------------------------
--------------------MEVANFYYEPDCLA-YGAKAARA------APR-APAAEP---
AIGEHERAIDFSPYLEPLAPAADFAAPAPAHH---------------DFLSDLF-
ADDYGAKPSKKPADYG-YVSLGRAGAKAAPPACF-PPPPPAALKAEPGFEP--ADCK-RADDA-
PAMAAGFP--FALRA---------------------YLGYQATP---SGSSGSLST-SSS-
SSPPGTP-SPADAKAA-------------------------PAACFAGPPAAP--
AKA-------------------KAKKT-----------VDKLSDEYKMRRERNNIAVRKSRDKAK-
MRNLETQHKVLELTAENERLQKKVEQLSRE---LSTLRNL-------FKQLP----EP-----
LLASAG--
HC------------------------------------------------------------------
--------------------------------------------------------------------
--------------------------------
        'M._musculus_gi_567757572_ref_NP_001274668.1_'  ------------
--------------------------------------------------------------------
--------------------------------------------------------------------
--------------------------------------------------------------------
--------------------------------------------------------------------
--------------------------------------------------------------------
--------------------------------------------------------------------
--------------------------------------------------------------------
--------------------------------------------------------------------
-----------------------------------------------MAAGFP--
FALRA---------------------YLGYQATP---SGSSGSLST-SSS-SSPPGTP-
SPADAKAA-------------------------PAACFAGPPAAP--
AKA-------------------KAKKT-----------VDKLSDEYKMRRERNNIAVRKSRDKAK-
MRNLETQHKVLELTAENERLQKKVEQLSRE---LSTLRNL-------FKQLP----EP-----
LLASAG--
HC------------------------------------------------------------------
--------------------------------------------------------------------
--------------------------------
        'H._sapiens_gi_551895123_ref_NP_001272808.1_'   ------------
--------------------------------------------------------------------
--------------------------------------------------------------------
--------------------------------------------------------------------
--------------------------------------------------------------------
--------------------------------------------------------------------
--------------------------------------------------------------------
--------------------------------------------------------------------
--------------------------------------------------------------------
-----------------------------------------------MAAGFP--
YALRA---------------------YLGYQAVP---SGSSGSLST-SSS-SSPPGTP-
SPADAKAP-------------------------
PTACYAGAAPAPSQVKS-------------------KAKKT-----------
VDKHSDEYKIRRERNNIAVRKSRDKAK-MRNLETQHKVLELTAENERLQKKVEQLSRE---
LSTLRNL-------FKQLP----EP-----LLASSG--
HC------------------------------------------------------------------
--------------------------------------------------------------------
--------------------------------
        'X._laevis_lcl_NM_001095915.1_prot_NP_001089384.1_1'    ----
--------------------------------------------------------------------

--------------------------------------------------------------------
--------------------------------------------------------------------
--------------------------------------------------------------------
--------------------------------------------------------------------
--------------------------------------------------------------------
---MHRLLQWDPAAAAACLPP---GVRSMD----YYDNDCLAGLVGKVPRR------
VPRGCPGSDS---SIGDHERAIDFSPYLEPPAGALGGGSPSAAPPG--------------
DFLSDLLGADEYKCG-RKGALEYS-PVVRGLGG-----------YPQLGETKVEPVFES--LEPY-
KGPGR-EENAMPSP--YSVRS---------------------YLTYQTVP---SGSSGNLSSASSS-
SSPPGTP-NPLESKS--------------------------G--GTSGGGY----
GKG-------------------KSKKS-----------LDKHSDEYKIRRERNNIAVRKSRDKAK-
VRNMETQHKVLELSAENERLQKRVEQLSRE---LSTLRNL-------FKQLP----EP-----
LLAATG--
RC------------------------------------------------------------------
--------------------------------------------------------------------
--------------------------------
        'X._laevis_lcl_NM_001172167.1_prot_NP_001165638.1_1'     ----
--------------------------------------------------------------------
--------------------------------------------------------------------
--------------------------------------------------------------------
--------------------------------------------------------------------
--------------------------------------------------------------------
--------------------------------------------------------------------
---MHRLPQWD--QAAACLPPPP-GIRSMD----YYDSDYLAGLVGKVPRR------
VPRGCPGSDS---SIGDHERAIDFSPYLEPPAGALGAGSPSAAPPG--------------
DFLSDLLGADEYKCG-RKGALEYS-SGWERTGG-----------VPQLGETKVEPVFES--LEPY-
KGPGR-EDNAMQSP--YSVRA---------------------YLGYQTVP---SGSSGNLSSASSS-
SSPPGTP-NPLDSKSE-------------------------GPSGASGTGYRKS-
GSG-------------------KAKKS-----------LDKQSNDYKLRRERNNIAVRKSRDKAK-
IRNMETQHKVLELSAENERLQKRVEQLSRE---LGTLRNL-------FKQVP----EP-----
LLAVTG--
RC------------------------------------------------------------------
--------------------------------------------------------------------
--------------------------------
        'D._rerio_lcl_NM_131884.2_prot_NP_571959.2_1'   ------------
--------------------------------------------------------------------
--------------------------------------------------------------------
--------------------------------------------------------------------
--------------------------------------------------------------------
--------------------------------------------------------------------
--------------------------------------------------------------------
--------------------MEVAG-
FYEGDYLAFHSTNASSSPVSDGVCKQPVNGSMTKLHDISEHEKAIDFSIYLDSLPQYQHLASQDESHR
HR-----------ALGIYSDFL-SEGNKSK-RAALQNYKNYISLTERD-----PNQL-
AYPELQETRIDAVFSPDFMGSFAKSNGRHEETPMDGPGGYDMRS---------------------
YLPYQTAP---SGSLGNISTASSSCSSPPGTP-APS-GKGR-------------------------
SP--QAGGKMT---SSG-------------------KGKKR-----------
LDKDSDEYRQRRERNNLAVRKSRDKAK-MRNLETQHKVLELAAENDRLQKRVEQLSRE---
LATLRN------------------------LLSATG--
QC------------------------------------------------------------------
--------------------------------------------------------------------
--------------------------------
        'D._melanogaster_gi_665403432_ref_NP_001286836.1_'       ----
--------------------------------------------------------------------

--------------------------------------------------------------------
--------------------------------------------------------------------
--------------------------------------------------------------------
--------------------------------------------------------------------
--------------
MLNMESPQMYADAVQTLAQLDLKKEPPLQQATIGQITLTAMSTAQQQQQQQQQQQQQQQQQQQQQQQQ
PQQQTTDANNNTSQDAALLVKQHAMHQMQQVAALGSNNNLLQKQMLQQYSTQTDLDELTTQEITLDLQ
HLIDDQFRDTETLGIFSDMVTSPGGLSATLPPSGMVSAAAKVLQQQTLRNQHGYGGRGGGGGAGGALA
YMPQPVHAT--
YNNSSDENSSVGSDSSTIKEEPIDPEYRRHLQEAASQQAAFMGNGAGLYNGYGSGANGLTGGGNPLNG
GNTTPSSNGSNGSTGSSNGSQFTNLTTANVLAHHNLP-------------------------
HLAAAAGAHNLLKQHSKLHAQQQHQQHQ-QQQQHRKHSNKH-----------
VDKGTDEYRRRRERNNIAVRKSREKAK-VRSREVEERVKSLLKEKDALIRQLGEMTNE---
LQLHKQI-------YMQLMNHANPEV----
SRVCRSFLNTNEHSL-----------------------------------------------------
--------------------------------------------------------------------
----------------------------------------
'N._vectensis_gi_156384801_ref_XP_001633321.1_' ------------
--------------------------------------------------------------------
--------------------------------------------------------------------
--------------------------------------------------------------------
--------------------------------------------------------------------
--------------------------------------------------------------------
--------------------------------------------------------------------
--------------------------------------------------------------------
--------------------------------------------------------------------
--------------------------------------------------------------------
--------------------------------------------------------------------
---------------------------------------------------M------------
DKHSEEYRQKRERNNVAVRKSRFKSK-QKFIETQSRVEELTEENERLHSRIDIITKE---
LNALRSL-------
FS------------------------------------------------------------------
--------------------------------------------------------------------
--------------------------------------------------------

uncharacterized_protein_LOC110247154__Exaiptasia_pallida_ ----
--------------------------------------------------------------------
--------------------------------------------------------------------
--------------------------------------------------------------------
--------------------------------------------------------------------
--------------------------------------------------------------------
--------------------------------------------------------------------
--------------------------------------------------------------------
--------------------------------------------------------------------
------MDVLEDLYETG-EWSAASSPCSELFDDIP-
SPASSLEDSCISDFSIDLTTYLDPAALSTSPEDLCKYLPHHNEAHPVEV-E---
SSCRYPHVTIKSEPKDAYPQKPIMSFNKPPAVVKESQQK--K----------------------
QLPS--AQLKAPSSRTSRKHS----------------NNKRM-----------
PDKHTEEYRQKRERNNIAVRKSRFKTK-QKFVDTLMKVEELTEENDRLRSRVDHLTKE---
LSALRGL-------FSNXSIFKDQALA---RALL------AQGLASS-
LGHSS---------------------------------------------------------------
--------------------------------------------------------------------
----------------------
'A._digitifera_lcl_XM_015902730.1_prot_XP_015758216.1_1'

```
        ------------------------------------------------------------
--------------------------------------------------------------------
--------------------------------------------------------------------
--------------------------------------------------------------------
--------------------------------------------------------------------
--------------------------------------------------------------------
-----------------------------------------------------------------
MTNAITRNGKKAALKCFVCHQVVLSSQPQRFGSVVENDRVSKPDHFFKKFTGQVAKIPPLKELDFPLP
IFDGLDLKQESQACVYPACNPIDVL-DLYESSTEWSAANSPSSELFEDVDLNSASDLDD--
LNDISIDFASYLDPATLSTSPEDLTKYLAPKKVSKRREVLEPYLSKVEQGYVPAKIVSK-
PYPSKPAMTY-KSTTLAKDHEQKSTT----------------------RLPSIASHLKAPKSRRG-
SSS----------------FKKVA-----------LEKGSEEYRMKRERNNIAVRKSRFKSK-
QKYVETQNKVDELEEENARLQSKVDFLTKE---LNVLRSL-------FSNTGAYKDPVVN---
AALL------
SHGISTAPHGLVASR-----------------------------------------------------
--------------------------------------------------------------------
------------------------------
        'H._vulgaris_gi_449684526_ref_XP_002164910.2_'  ------------
--------------------------------------------------------------------
--------------------------------------------------------------------
--------------------------------------------------------------------
--------------------------------------------------------------------
--------------------------------------------------------------------
--------------------------------------------------------------------
--------------------------------------------------------------------
-------MIPTAIFQGMENIKNTDIQAVPNMESDNAVKSPTDSLFSECVLDFSKYLSTADLPAA-
PGEFFDDLFHGKLSKDALQRHLPGLI-KIKKEPDSTATSRCSTPESLLPEHDFDNGDFPT-
PQFQDNIINPRYQKGGLMINQFQYIPRSSGTSENDSDDEINVEVLDFNAQRLKAALIKLEQSKGDD--
--------------------NVSHR-NELKSPRSKSR----------------------
KTM------------SKGSEEYLVKRERNNVAVRKSRTKAK-
LKHIETQMRVGELTEENDQLRNRIASLQKE---LDAMKSF-------FEYNSLNEKSSYF---
SSYP------
AMFSATKLL-----------------------------------------------------------
--------------------------------------------------------------------
------------------------------
        'H._vulgaris_gi_449684528_ref_XP_002164958.2_'  ------------
--------------------------------------------------------------------
--------------------------------------------------------------------
--------------------------------------------------------------------
--------------------------------------------------------------------
--------------------------------------------------------------------
--------------------------------------------------------------------
--------------------------------------------------------------------
-------MIPTAIFQ-------------------------------------------ADLPAA-
PGEFFDDLFHGKLSKDALQRHLPGLI-KIKKEPDSTATSRCSTPESLLPEHDFDNGDFPT-
PQFQDNIINPRYQKGGLMINQFQYIPRSSGTSENDSDDEINVEVLDFNAQRLKAALIKLEQSKGDD--
--------------------NVSHR-NELKSPRSKSR----------------------
KTM------------SKGSEEYLVKRERNNVAVRKSRTKAK-
LKHIETQMRVGELTEENDQLRNRIASLQKE---LDAMKSF-------FEYNSLNEKSSYF---
SSYP------
AMFSATKLL-----------------------------------------------------------
--------------------------------------------------------------------
------------------------------
        'T._adherens_lcl_XM_002113628.1_prot_XP_002113664.1_1'  ----
```

--------------------------------------------------------------------
--------------------------------------------------------------------
--------------------------------------------------------------------
--------------------------------------------------------------------
--------------------------------------------------------------------
--------------------------------------------------------------------
--------------------------------------------------------------------
--------------------------------------------------------------------
--------------------------------------------------------------------
--------------------------------------------------------------------
-------------------------------SNSR----------------------
KSL------------NKYSDEYRRKRERNNEAVRKSRKKTK-
LKSMETQERVMQLSMENEELKTKLSLLTKE---LSVLKSL-------
F-------------------------------------------------------------------
--------------------------------------------------------------------
--------------------------------------------------------
        'H._vulgaris_gi_449679187_ref_XP_004209260.1_'  ------------
--------------------------------------------------------------------
--------------------------------------------------------------------
--------------------------------------------------------------------
--------------------------------------------------------------------
--------------------------------------------------------------------
--------------------------------------------------------------------
--------------------------------------------------------------------
--------------------------------------------------------------------
--------------------------------------------------------------------
---------------------------MDSNEDSGEHEIQPNENWYTTNAEIA--
S----------------------QVISKYTGATSSSNKRK----------------------
QTN-----------TKPSDEEYSRKRARNNVAVKKSREKAK-
NRIVETQVRVEQLSQENEELQTKVTLLTKE---LNVLRAL-------
FTNGGFALPGELQIVSNNSNNE-
LSQNQHQNNEQSTSNGGNHNLLNKEVKMSFDTKIQPLKPMPRVFTNSQKSLLSSSQQSSDSFNPYERN
NESQLFTYNPKYTPATEYVKKETKKSNCHTILVSPDTTPQRSSSLLYQNGAEMRHTSVIQTVTSQPQQ
QQKSSLTQNSLGKFCIIQDPEKVGQVKIVPLDS
        'N._vectensis_gi_156375819_ref_XP_001630276.1_' ------------
--------------------------------------------------------------------
--------------------------------------------------------------------
--------------------------------------------------------------------
--------------------------------------------------------------------
--------------------------------------------------------------------
--------------------------------------------------------------------
--------------------------------------------------------------------
--------------------------------------------------------------------
--------------------------------------------------------------------
--------------------------------------------------------------------
-------------------------------------------------------------------
DDEYIRKRERNNEAVRKSRKKAK-QRIQETQQRVTELSKENEELRSKVTLLQKE---
LSVLRSL-------
F-------------------------------------------------------------------
--------------------------------------------------------------------
--------------------------------------------------------
        'A._digitifera_lcl_XM_015919129.1_prot_XP_015774615.1_1'
        ------------------------------------------------------------
--------------------------------------------------------------------

```
--------------------------------------------------------------------
--------------------------------------------------------------------
--------------------------------------------------------------------
--------------------------------------------------------------------
--------------------------------------------------------------------
--------------------------------------------------------------------
--------------------------------------------------------------------
--------------------------------------------------------------------
-MESA--A----------------------QNVHP-----
KPAKAVQ----------------------QSV-----------
DKEIGDDYVKKRERNNIAVRKSRMKAK-ERIEETRRRVVDLSKENEELRNKVSLLQKE---
LNVLRSL-------
FANGGISVPAEIDLQFTNCNNDNHGVTVLTAVTTSLTNGSQENGVSSSETLENGKSTHAIKKES----
--------------------------------------------------------------------
--------------------------------------------------------
        'CCAAT_enhancer_binding_protein_gamma_like__Exaiptasia_palli
da_'    ------------------------------------------------------------
--------------------------------------------------------------------
--------------------------------------------------------------------
--------------------------------------------------------------------
--------------------------------------------------------------------
--------------------------------------------------------------------
--------------------------------------------------------------------
--------------------------------------------------------------------
--------------------------------------------------------------------
------------------------------------------------------------------
MPDNESK--S----------------------DANSE-----
DPTSPID----------------------KAG------------
MDADEEYVKKRERNNIAVRKSRRKAK-QRIEETKQRVDDLTKENEDLRIKVALLQKE---
LNVLRSL-------FANGGVANGNSLP---NGVN------
VIFTTNSNSSTNSTEQNSPQIEVKTEADEND-------------------------------------
--------------------------------------------------------------------
------------------------------
        'A._californica_gi_325120973_ref_NP_001191392.1_'       ----
--------------------------------------------------------------------
--------------------------------------------------------------------
--------------------------------------------------------------------
--------------------------------------------------------------------
--------------------------------------------------------------------
--------------------------------------------------------------------
---------------------------------------
MDSLQLSELQTFFSTNKGELTSKKLFVESIDCAPLDLLDPILERVLDLSEYIPTNSSLDSHIISTHIS
PDPTDLQHQSLPQVVLDATALVHEEKYPGEGSVLPPPVVNFLESGQSPEQSDVTSYYSDDSNSATQQM
PEFSQLTSPGVDAPSTSGFQWPNLVKREDFGVRTDDFGFTPCHISAKHPHTHTSVPSSIGGD-
GASLPYVTSPTTSLPSPTSTIYSASSPGTP----------------------
GPSTPSSRGRRPHGSAAPYSTK---------------SRRRQ-----------
VEKGTQEYVDKRARNNVAVRKSRAKAK-EKQRETEGRVKSLLDQNDQLQKKVDLLSKE---
LNVLKGL-------
FLNVGASIPDEFLKLMGDS-------------------------------------------------
--------------------------------------------------------------------
--------------------------------------------------------
        'CCAAT_enhancer_binding_protein_gamma_OS_Homo_sapiens_OX_960
6_GN_CEBPG_PE_1_SV_1'   --------------------------------------------
--------------------------------------------------------------------
```

```
--------------------------------------------------------------------
--------------------------------------------------------------------
--------------------------------------------------------------------
--------------------------------------------------------------------
--------------------------------------------------------------------
--------------------------------------------------------------------
--------------------------------------------------------------------
------------------------------------MSK---
ISQQNSTPGVNGISVIHTQAH--------ASGLQ--
QVPQLVPAGPG--------------------------------
GGGKAVAPSKQS------------------KKSSP-----------
MDRNSDEYRQRRERNNMAVKKSRLKSK-QKAQDTLQRVNQLKEENERLEAKIKLLTKE---
LSVLKDL-------FLEHAHNLAD--------NVQS--ISTENTTAD-----
GDNAGQ--------------------------------------------------------------
--------------------------------------------------------------------
------------------
        'M._musculus_gi_61966683_ref_NP_034014.1_'      ------------
--------------------------------------------------------------------
--------------------------------------------------------------------
--------------------------------------------------------------------
--------------------------------------------------------------------
--------------------------------------------------------------------
--------------------------------------------------------------------
--------------------------------------------------------------------
--------------------------------------------------------------------
--------------------------------------------------------------------
MSK---LSQPATTPGVNGISVIHTQAH--------ASGLQ--
QVPQLVPAGPG--------------------------------
GGGKAVPPSKQS------------------KKSSP-----------
MDRNSDEYRQRRERNNMAVKKSRLKSK-QKAQDTLQRVNQLKEENERLEAKIKLLTKE---
LSVLKDL-------FLEHAHSLAD--------NVQP--ISTETTATN-----
SDNPGQ--------------------------------------------------------------
--------------------------------------------------------------------
------------------
        'D._rerio_lcl_NM_131886.1_prot_NP_571961.1_1'   ------------
--------------------------------------------------------------------
--------------------------------------------------------------------
--------------------------------------------------------------------
--------------------------------------------------------------------
--------------------------------------------------------------------
--------------------------------------------------------------------
--------------------------------------------------------------------
--------------------------------------------------------------------
--------------------------------------------------------------------
MSKQ--LQQKISSTDQNGVSVIQNQPHNSALNPAGAAGLQ--
QVPQLVPVNPG--------------------------------
GGGKATAPSKM-------------------KKSN------------
MDKDSDEYRQRRERNNLAVKKSRMRSK-QKAQDTQQRVNELKEENERLEAKIKLLSKE---
LSVLKDL-------FLEHAHNLAD--------NVQP--PASGGGPGDL---
CNNNSGSNSSQ---------------------------------------------------------
--------------------------------------------------------------------
-------------------
        'X._laevis_lcl_NM_001095901.1_prot_NP_001089370.1_1'    ----
--------------------------------------------------------------------
```

```
--------------------------------------------------------------------
--------------------------------------------------------------------
--------------------------------------------------------------------
--------------------------------------------------------------------
--------------------------------------------------------------------
--------------------------------------------------------------------
--------------------------------------------------------------------
--------------------------------------------------------------------
--------MDKLDQMNQSPSTAS-EGLS----DAL---------PGSP--
ATPQRVPLNPG--------------------------------
GGGKATPPSKNS------------------KKSQR-----------
LERGSEEYRQRRERNNMAVKKSRLKSK-QKAQDTLQRVNQLKEENERLEAKIKLLTKE---
LSVLKDL-------FLEHAHNLSD--------NVQP--ESS--TPGQ------
ESAG----------------------------------------------------------------
--------------------------------------------------------------------
-----------------
        'X._laevis_lcl_XM_018261138.1_prot_XP_018116627.1_1'    ----
--------------------------------------------------------------------
--------------------------------------------------------------------
--------------------------------------------------------------------
--------------------------------------------------------------------
--------------------------------------------------------------------
--------------------------------------------------------------------
--------------------------------------------------------------------
--------------------------------------------------------------------
--------------------------------------------------------------------
--------------MNQSPSSAS-EGLS----DAL---------PGSP--
ATPQLVPLNPG--------------------------------
GGGKATPPSKNS------------------KKSQR-----------
LERGSEEYRQRRERNNMAVKKSRLKSK-QKAQDTMQRVNQLKEENERLEAKIKLLTKE---
LSVLKDL-------FLEHAHNLSD--------NVQP--ESS--TPGQ------
ESAG----------------------------------------------------------------
--------------------------------------------------------------------
-----------------
        'S._purpuratus_gi_390355599_ref_XP_003728584.1_'        ----
--------------------------------------------------------------------
--------------------------------------------------------------------
--------------------------------------------------------------------
--------------------------------------------------------------------
--------------------------------------------------------------------
--------------------------------------------------------------------
--------------------------------------------------------------------
--------------------------------------------------------------------
--------------------------------------------
MASTSNPSLLPQKNIKQEIIVVFEPPAREMASSSSHSSVPQPETSQEINVVPVSPVKDMPSTSNP--
SSGHETGQENDMVPDSTGIEL----------------------
SKDDLKAGGGKVGSSSK--------------------KKSG------------
CEKDSDEYKRRRERNNEAVRKSRQKSR-QKASETEVRVTELKKENADLEQRVTLLHKE---
LELLKDL-------
FLTHANELPDPSTTFGLFNANPRLGSSSPNPALSRRIVLKTESLTVSLTCRNVPESITTTT-------
--------------------------------------------------------------------
--------------------------------------------------------
        'S._kowalevskii_gi_585693956_ref_XP_006821724.1_'       ----
--------------------------------------------------------------------
```

```
--------------------------------------------------------------------
--------------------------------------------------------------------
--------------------------------------------------------------------
--------------------------------------------------------------------
--------------------------------------------------------------------
--------------------------------------------------------------------
--------------------------------------------------------------------
--------------------------------------------------------------------
------------------------------------------------------
MAPDSTK-----------------------------------
SPHQASKT-------------------KKNK------------
PTKESEEYKKRRARNNIAVRKSRTKTK-MRTLDTLKKVNELKAENEQLEVKVKLLSKE---
LSLLKDL-------FLAHAGHMPDTS------GSNS--
NCTAIDGGTSCCAITNDAGNATIKAEA-----------------------------------------
--------------------------------------------------------------------
--------------------------------
        'A._californica_gi_524898017_ref_XP_005105463.1_'        ----
--------------------------------------------------------------------
--------------------------------------------------------------------
--------------------------------------------------------------------
--------------------------------------------------------------------
--------------------------------------------------------------------
--------------------------------------------------------------------
--------------------------------------------------------------------
--------------------------------------------------------------------
--------------------------------------------------------------------
----------------------------------------------
MPPKKTYESYESDSD--------------------------------
GDSQQSQTRGGG------------------GKRQK-----------
LDKNSDEYKKRRERNNVAVRKSREASR-QKAKDTMEKVARLREENRALEQKVTILNKE---
LGVLRDL-------FLTHASATAAAQAAT-VTKLLPKSEVEEDTASKD---
SSENSETVIKDHKYFVTQKDA-----------------------------------------------
--------------------------------------------------------------------
-------------------
        'S._kowalevskii_gi_585693879_ref_XP_006821714.1_'        ----
--------------------------------------------------------------------
--------------------------------------------------------------------
--------------------------------------------------------------------
--------------------------------------------------------------------
--------------------------------------------------------------------
--------------------------------------------------------------------
--------------------------------------------------------------------
-----------------------------------MD-
SPANFYDSEVDDTKPKSNNSVFLFDETVLADYSDIYRNEGSIDFSLYLQGTAAANSPSTTVKN--
DDIYNELDF-
LPASCNPALTITPTPTDSTLPELIEVNQIESKTSKEDTEYMSQIGENRAPPTSVKTEAQEPEKIVVTI
KTEKDASSPSVCQHTYSTS----------------------TP-
ETRNRLKSAPASKSLK------------------GKRTI-----------DK-
NSEEYRHRREKNNVAVRRSREKSK-VKQKEVQNKVSQLQDENDKLQKKVELLTKE---
LTVLKSL-------FTNVGVTPPVLSG---
SDSM----------------------------------------------------------------
--------------------------------------------------------------------
----------------------------------------
        'S._purpuratus_gi_390340887_ref_XP_003725328.1_'        ----
```

```
--------------------------------------------------------------------
--------------------------------------------------------------------
--------------------------------------------------------------------
--------------------------------------------------------------------
--------------------------------------------------------------------
--------------------------------------------------------------------
--------------------------------------------------------------------
-----------------MDSFSQILSQCHLDSAKVMDPSADNFYVCE-
EKQHFTIKQEPFDIEDSNVGLVGDFLQSEDSIDIDAYLALGRAEVQQNNHVDNTGDGILRQLEQNFAS
NHQQMLTHTPAPTPGAQPNQQPVHSVD-
QLHYATINTMPQQNTQAHVVEQAVPEYQPCWEIHSTSNSAASSPGLPTSGDSYNEE------------
----------VQIQADSGTASGSGGGSQK------------------RKRPV-----------
PTPGTHEYKQKRERNNIAVRKSREKTK-TKNKELQDKVGELQEENTGLKKRVEGLAKE---
LAVLRSL-------XXXXXXXXXXXXX---
XXXXVIRECRVRFREWRPETFSRPKIPLT---------------------------------------
--------------------------------------------------------------------
----------------------------------------
        'A._queenslandica_gi_340367947_ref_XP_003382514.1_'      ----
--------------------------------------------------------------------
--------------------------------------------------------------------
--------------------------------------------------------------------
--------------------------------------------------------------------
--------------------------------------------------------------------
--------------------------------------------------------------------
--------------------------------------------------------------------
------------------------------------------------------------------
MADKDSLYSIEQYLGLAASAKATSGNLYPFGQAPPSSVHLGFQLEEESVDFSELVKACEQKDYYSQTT
NDYYSSGYWYQQKPPGMPIRSEEETSTLVGVGTTSNPHDVTNVSANPTLPPVASSLSAPSPPP-----
---------------------------TVQDTAARTRQTT-----------------
KKQKI-----------NDKCTDDYKDKRHRNNIAVRKSRSKFR-
KRVLETEKRVQELEENNAKLKNYVALLQKE---LAVLKGL-------
FSSASNASGDERYS------------------------------------------------------
--------------------------------------------------------------------
--------------------------------------------------------

;
end;

begin trees;
        tree tree_1 = [&U] [&branchAttributeNames={"Bootstrap"}]
('S._kowalevskii_gi_585693090_ref_XP_002738815.2_':
0.21229356389999987,('S._purpuratus_gi_72006198_ref_XP_787318.1_':
0.2880140916,
(((('T._adherens_lcl_XM_002109162.1_prot_XP_002109198.1_1':
0.17931180349999964,
((hepatic_leukemia_factor_like__Exaiptasia_pallida_:
0.14783492769999995,'N._vectensis_gi_156388041_ref_XP_001634510.1_':
0.03942872130000019)[&Bootstrap=77]:
0.000002,'A._digitifera_lcl_XM_015913431.1_prot_XP_015768917.1_1':
0.18598628019999985)[&Bootstrap=100]:0.4395103368000002)
[&Bootstrap=93]:0.20256855730000023,
((((('S._kowalevskii_gi_291222488_ref_XP_002731253.1_':
0.000003,'A._californica_gi_524895551_ref_XP_005104258.1_':
0.40137491270000014)[&Bootstrap=27]:0.000002,
```

('S._purpuratus_gi_390352986_ref_XP_785519.3_':
0.1167450661000002,'A._californica_gi_524915872_ref_XP_005112723.1_
':0.15376387570000016)[&Bootstrap=23]:0.01916359589999983)
[&Bootstrap=58]:
0.1589842571000002,'A._californica_gi_524899363_ref_XP_005106114.1_'
:0.000002)[&Bootstrap=54]:
0.09957303480000013,'S._kowalevskii_gi_291234179_ref_XP_002737021.1_
':0.04916741660000046)[&Bootstrap=90]:0.21459068299999995,
((((((((((('N._vectensis_gi_156408121_ref_XP_001641705.1_':
0.12660161850000007,
(thyrotroph_embryonic_factor_like_2__Exaiptasia_pallida_:
0.000003,'S._kowalevskii_gi_585674208_ref_XP_002736397.2_':
1.288877364)[&Bootstrap=86]:0.32065762650000007)[&Bootstrap=65]:
0.18751394949999978,'A._digitifera_lcl_XM_015908039.1_prot_XP_015763
525.1_1':0.31389925019999976)[&Bootstrap=48]:0.08151918650000045,
(((('N._vectensis_gi_156336982_ref_XP_001619763.1_':
0.000002,'N._vectensis_gi_156400985_ref_XP_001639072.1_':0.000002)
[&Bootstrap=100]:0.11983378159999991,
(thyrotroph_embryonic_factor_like_1__Exaiptasia_pallida_:
0.000002,Transcription_factor_VBP__Exaiptasia_pallida_:0.000002)
[&Bootstrap=100]:0.17976590409999993)[&Bootstrap=90]:
0.17855206479999985,
(('H._vulgaris_gi_221106316_ref_XP_002167159.1_':
0.000002,'H._vulgaris_gi_221116695_ref_XP_002159994.1_':
1.0976176762)[&Bootstrap=94]:
0.6437388153999999,'T._adherens_lcl_XM_002117855.1_prot_XP_002117891
.1_1':0.9403688420999998)[&Bootstrap=22]:0.000002)[&Bootstrap=17]:
0.000002,'A._digitifera_lcl_XM_015892501.1_prot_XP_015747987.1_1':
0.18504367930000032)[&Bootstrap=46]:0.14977629869999998)
[&Bootstrap=28]:0.07005414599999948,
('A._digitifera_lcl_XM_015892500.1_prot_XP_015747986.1_1':
0.21942019070000018,'N._vectensis_gi_156408099_ref_XP_001641694.1_':
0.29291342190000025)[&Bootstrap=85]:0.25261540849999964)
[&Bootstrap=73]:0.15033047869999994,
('A._queenslandica_gi_340372715_ref_XP_003384889.1_':
0.4129702057000002,((((((A0A1D5NSQ7_DANRE:
0.000002,'D._rerio_lcl_NM_131837.1_prot_NP_571912.1_1':
0.014696716400000032)[&Bootstrap=100]:0.33928445640000016,
((((CCAAT_enhancer_binding_protein_delta_OS_Homo_sapiens_OX_9606_GN_
CEBPD_PE_1_SV_2:0.000002,'M._musculus_gi_110347410_ref_NP_031705.3_'
:0.000002)[&Bootstrap=100]:0.08112165140000016,
(('X._laevis_lcl_NM_001089607.2_prot_NP_001083076.1_1':
0.006801277500000147,'X._laevis_lcl_NM_001089609.1_prot_NP_001083078
.1_1':0.042785702299999784)[&Bootstrap=86]:
0.09553496789999993,'D._rerio_lcl_NM_131887.1_prot_NP_571962.1_1':
0.1107416898000021)[&Bootstrap=71]:0.07228188519999978)
[&Bootstrap=98]:0.14184132380000003,
(((('X._laevis_lcl_NM_001086806.1_prot_NP_001080275.1_1':
0.013044752399999915,'X._laevis_lcl_NM_001091687.1_prot_NP_001085156
.1_1':0.020741841900000058)[&Bootstrap=65]:0.017022053099999823,
((((((('H._sapiens_gi_28872794_ref_NP_004355.2_':
0.0,'M._musculus_gi_566559971_ref_NP_001274444.1_':0.0):
0.0,'M._musculus_gi_86198301_ref_NP_031704.2_':0.0):
0.0,'M._musculus_gi_566559986_ref_NP_001274450.1_':0.0):

0.0,'M._musculus_gi_567316233_ref_NP_001274452.1_':0.0):
0.0,'H._sapiens_gi_566559992_ref_NP_001274353.1_':0.0):
0.0,'H._sapiens_gi_551894998_ref_NP_001272758.1_':0.0):
0.0,'M._musculus_gi_566559969_ref_NP_001274443.1_':0.0):
0.000002,'H._sapiens_gi_566559994_ref_NP_001274364.1_':0.000002)
[&Bootstrap=100]:0.03818954290000095)[&Bootstrap=67]:
0.01387761890000055,'D._rerio_lcl_NM_131885.2_prot_NP_571960.1_1':
0.07603749919999991)[&Bootstrap=90]:0.12096283099999994,
(((('M._musculus_gi_6753404_ref_NP_034013.1_':
0.0,'M._musculus_gi_567757572_ref_NP_001274668.1_':0.0):
0.000002,'M._musculus_gi_567757569_ref_NP_001274667.1_':0.000002)
[&Bootstrap=98]:
0.03204426030000018,'H._sapiens_gi_551895123_ref_NP_001272808.1_':
0.000002)[&Bootstrap=87]:0.03412183369999999,
('X._laevis_lcl_NM_001095915.1_prot_NP_001089384.1_1':
0.000002,'X._laevis_lcl_NM_001172167.1_prot_NP_001165638.1_1':
0.07749933380000007)[&Bootstrap=79]:0.0291204209)[&Bootstrap=86]:
0.04725817600000015)[&Bootstrap=40]:
0.03052561360000057,'D._rerio_lcl_NM_131884.2_prot_NP_571959.2_1':
0.0446450801000001)[&Bootstrap=59]:0.06929677950000013)
[&Bootstrap=30]:0.01229088100000198,
((CCAAT_enhancer_binding_protein_epsilon_OS_Homo_sapiens_OX_9606_GN_
CEBPE_PE_1_SV_2:0.0171914447999999,'M._musculus_gi_46369479_ref_NP_9
97014.1_':0.03051955020000019)[&Bootstrap=54]:0.01856854489999928,
('X._laevis_lcl_XM_018243908.1_prot_XP_018099397.1_1':
0.03337192289999795,'X._laevis_lcl_XM_018259384.1_prot_XP_018114873
.1_1':0.0346946481999999)[&Bootstrap=97]:0.06725196249999987)
[&Bootstrap=87]:0.1431417129999999)[&Bootstrap=60]:
0.08445079409999989)[&Bootstrap=90]:0.21421875060000017,
(((((CCAAT_enhancer_binding_protein_gamma_OS_Homo_sapiens_OX_9606_GN
_CEBPG_PE_1_SV_1:0.000002,'M._musculus_gi_61966683_ref_NP_034014.1_'
:0.000002)[&Bootstrap=98]:0.000002,
('X._laevis_lcl_NM_001095901.1_prot_NP_001089370.1_1':
0.000002,'X._laevis_lcl_XM_018261138.1_prot_XP_018116627.1_1':
0.01523273570000166)[&Bootstrap=88]:0.03070715859999984)
[&Bootstrap=97]:
0.10282093600000008,'D._rerio_lcl_NM_131886.1_prot_NP_571961.1_1':
0.02801876889999999)[&Bootstrap=67]:
0.07208391120000002,'S._kowalevskii_gi_585693956_ref_XP_006821724.1_
':0.3767721514)[&Bootstrap=62]:0.04619829849999979,
('S._purpuratus_gi_390355599_ref_XP_003728584.1_':
0.26112501460000015,'A._californica_gi_524898017_ref_XP_005105463.1_
':0.25831472)[&Bootstrap=50]:0.186592558200001)[&Bootstrap=48]:
0.13360806189999996)[&Bootstrap=17]:0.000002,
(((('N._vectensis_gi_156384801_ref_XP_001633321.1_':
0.09336053110000009,uncharacterized_protein_LOC110247154__Exaiptasia
_pallida_:0.15325088730000003)[&Bootstrap=46]:0.02659163600000003,
('H._vulgaris_gi_449684526_ref_XP_002164910.2_':
0.000002,'H._vulgaris_gi_449684528_ref_XP_002164958.2_':0.000002)
[&Bootstrap=100]:0.4074395088)[&Bootstrap=41]:
0.08874008060000005,'A._digitifera_lcl_XM_015902730.1_prot_XP_015758
216.1_1':0.12186352440000014)[&Bootstrap=57]:0.14446929919999985,
(('T._adherens_lcl_XM_002113628.1_prot_XP_002113664.1_1':
0.2680208579999986,('H._vulgaris_gi_449679187_ref_XP_004209260.1_':

0.23936836669999995,
('N._vectensis_gi_156375819_ref_XP_001630276.1_':
0.03386181629999996,
('A._digitifera_lcl_XM_015919129.1_prot_XP_015774615.1_1':
0.19719946099999985,CCAAT_enhancer_binding_protein_gamma_like__Exaip
tasia_pallida_:0.17245514629999992)[&Bootstrap=91]:
0.11548731420000014)[&Bootstrap=85]:0.1279642741)[&Bootstrap=70]:
0.10591003919999986)[&Bootstrap=55]:
0.1344351345999999,'A._queenslandica_gi_340367947_ref_XP_003382514.1
_':0.7072061131000003)[&Bootstrap=24]:0.034294649900000085)
[&Bootstrap=33]:0.10006423919999996)[&Bootstrap=28]:0.1744933703,
(('D._melanogaster_gi_665403432_ref_NP_001286836.1_':
0.5603650778000002,'A._californica_gi_325120973_ref_NP_001191392.1_'
:0.35049996929999994)[&Bootstrap=73]:
0.1392586765999999,'S._purpuratus_gi_390340887_ref_XP_003725328.1_':
0.47488331670000017)[&Bootstrap=54]:0.06691424560000003)
[&Bootstrap=89]:
0.1600859144000002,'S._kowalevskii_gi_585693879_ref_XP_006821714.1_'
:0.000002)[&Bootstrap=100]:1.1441852279000004)[&Bootstrap=75]:
0.11225174979999952)[&Bootstrap=84]:0.20768454310000006,
((((((((('D._melanogaster_gi_24660427_ref_NP_729297.1_':
0.0,'D._melanogaster_gi_442630877_ref_NP_001261546.1_':0.0):
0.0,'D._melanogaster_gi_24660469_ref_NP_729303.1_':0.0):
0.0,'D._melanogaster_gi_442630873_ref_NP_001261544.1_':0.0):
0.0,'D._melanogaster_gi_24660458_ref_NP_729301.1_':0.0):
0.0,'D._melanogaster_gi_24660450_ref_NP_729300.1_':0.0):
0.0,'D._melanogaster_gi_24660438_ref_NP_729299.1_':0.0):
0.0,'D._melanogaster_gi_281365802_ref_NP_729302.2_':0.0):
0.0,'D._melanogaster_gi_442630875_ref_NP_001261545.1_':0.0):
0.0,'D._melanogaster_gi_281365800_ref_NP_729298.2_':0.0):
0.000002,'D._melanogaster_gi_442630879_ref_NP_001261547.1_':
0.000002)[&Bootstrap=100]:0.026417015799999888)[&Bootstrap=67]:
0.1483866652000032,'S._purpuratus_gi_193788685_ref_NP_001123286.1_'
:0.21767801090000027)[&Bootstrap=24]:0.0453134935999997,
(((('A._californica_gi_524895679_ref_XP_005104321.1_':
0.0,'A._californica_gi_524895683_ref_XP_005104323.1_':0.0):
0.000002,'A._californica_gi_524895681_ref_XP_005104322.1_':0.000002)
[&Bootstrap=100]:0.29607336390000016,
((((((('M._musculus_gi_226958409_ref_NP_705617.2_':
0.0,'M._musculus_gi_8394435_ref_NP_059072.1_':0.0):0.0,TEF_CHICK:
0.0):
0.0,Thyrotroph_embryonic_factor_OS_Homo_sapiens_OX_9606_GN_TEF_PE_1_
SV_3:0.0):0.0,'H._sapiens_gi_223972670_ref_NP_001138870.1_':0.0):
0.000002,'M._musculus_gi_568991979_ref_XP_006520806.1_':0.000002)
[&Bootstrap=100]:0.017520087399999884,
((('X._laevis_lcl_NM_001086779.1_prot_NP_001080248.1_1':
0.0,'X._laevis_lcl_XM_018256479.1_prot_XP_018111968.1_1':0.0):
0.000002,'X._laevis_lcl_XM_018256478.1_prot_XP_018111967.1_1':
0.000002)[&Bootstrap=100]:0.012623691499999978,
((('X._laevis_lcl_NM_001094595.1_prot_NP_001088064.1_1':
0.0,'X._laevis_lcl_XM_018259675.1_prot_XP_018115164.1_1':0.0):
0.0,'X._laevis_lcl_XM_018259676.1_prot_XP_018115165.1_1':0.0):
0.000002,'X._laevis_lcl_XM_018259674.1_prot_XP_018115163.1_1':
0.000002)[&Bootstrap=100]:0.019438410400000272)[&Bootstrap=81]:

```
0.053015924299999906)[&Bootstrap=67]:0.09129349239999973,
(('X._laevis_lcl_XM_018235125.1_prot_XP_018090614.1_1':
0.000002,'X._laevis_lcl_XM_018235126.1_prot_XP_018090615.1_1':
0.000002)[&Bootstrap=91]:0.030733462700000214,
('X._laevis_lcl_XM_018240540.1_prot_XP_018096029.1_1':
0.000002,'X._laevis_lcl_XM_018240541.1_prot_XP_018096030.1_1':
0.000002)[&Bootstrap=72]:0.000002)[&Bootstrap=48]:0.000002)
[&Bootstrap=35]:0.044704462500000375)[&Bootstrap=29]:
0.03667595629999987,
((('H._sapiens_gi_530412024_ref_XP_005257326.1_':
0.0,'M._musculus_gi_568973279_ref_XP_006533068.1_':0.0):
0.000002,'M._musculus_gi_568973277_ref_XP_006533067.1_':0.000002)
[&Bootstrap=97]:0.000002,((HLF_HUMAN:
0.0,'M._musculus_gi_568973283_ref_XP_006533070.1_':0.0):
0.000002,'M._musculus_gi_31982951_ref_NP_766151.1_':0.000002)
[&Bootstrap=94]:0.000002)[&Bootstrap=91]:0.000002)[&Bootstrap=24]:
0.030538839999999734)[&Bootstrap=8]:0.000002,
('D._rerio_lcl_NM_001077334.2_prot_NP_001070802.1_1':
0.04309626060000005,
('D._rerio_lcl_NM_001197059.1_prot_NP_001183988.1_1':
0.000002,'D._rerio_lcl_XM_005156334.4_prot_XP_005156391.1_1':
0.000002)[&Bootstrap=95]:0.042009797900000034)[&Bootstrap=55]:
0.016749706799999764)[&Bootstrap=38]:0.06016906830000002,
((('D._rerio_lcl_NM_131400.1_prot_NP_571475.1_1':
0.0,'D._rerio_lcl_XM_005156136.3_prot_XP_005156193.1_1':0.0):
0.000002,'D._rerio_lcl_XM_005156135.3_prot_XP_005156192.1_1':
0.000002)[&Bootstrap=100]:
0.06428118010000006,'D._rerio_lcl_NM_001020661.1_prot_NP_001018497.1
_1':0.022243962700000175)[&Bootstrap=78]:0.05323115589999983)
[&Bootstrap=42]:0.026791631800000104,
((('D._rerio_lcl_NM_001197062.1_prot_NP_001183991.1_1':
0.000002,'D._rerio_lcl_XM_005157897.4_prot_XP_005157954.1_1':
0.000002)[&Bootstrap=100]:0.000002,
(((('M._musculus_gi_170650717_ref_NP_058670.2_':
0.0,'M._musculus_gi_568947267_ref_XP_006540660.1_':0.0):
0.0,DBP_HUMAN:0.0):
0.000002,'M._musculus_gi_568947265_ref_XP_006540659.1_':0.000002)
[&Bootstrap=100]:0.04674449269999981,
('X._laevis_lcl_XM_018226265.1_prot_XP_018081754.1_1':
0.000002,'X._laevis_lcl_XM_018228082.1_prot_XP_018083571.1_1':
0.000002)[&Bootstrap=100]:0.1067225088999999)[&Bootstrap=92]:
0.11096382760000001)[&Bootstrap=52]:
0.020926261500000l,'D._rerio_lcl_NM_001197060.1_prot_NP_001183989.1_
1':0.050697023200000135)[&Bootstrap=87]:0.06761012629999996)
[&Bootstrap=77]:0.20381927320000015)[&Bootstrap=68]:
0.038676952699999934)[&Bootstrap=65]:
0.019364605499999854,'A._californica_gi_524907652_ref_XP_005108942.1
_':1.0896644650999998)[&Bootstrap=73]:
0.15025465820000017,'D._melanogaster_gi_24654082_ref_NP_611101.1_':
0.17388480920000005)[&Bootstrap=93]:0.2243268327000001)
[&Bootstrap=70]:
0.012936450299999791,'S._kowalevskii_gi_291227743_ref_XP_002733842.1
_':0.28947242699999975);
end;
```
