## Supplementary Table 1 for "Photoreceptor complexity accompanies adaptation to challenging marine environments in Anthozoa"

| Oligo sequence (5' to 3') | Name |
|---|---|

**qPCR**

| | |
|---|---|
| GCGAAGTGTTGTACCCTGGA | 6-4.1/2_qPCR_F |
| CAAACCCAGTCCTGCTTTGC | 6-4.1/2_qPCR_R |
| CTCCCAGCGATCTCCAAAGG | AnthoCRY.1_qPCR_F |
| CCAGGTGCATTGGAAACTTTGT | AnthoCRY.1_qPCR_R |
| AGCGAGCCAGAATGACAACT | AnthoCRY.2_qPCR_F |
| TCTTGTCTTAGCTGACGCATGA | AnthoCRY.2_qPCR_R |
| TGTCCCTCTCTGTCAGGTTGA | CPD_II.1_qPCR_F |
| GCACTCCAGTCAGCTCCTTT | CPD_II.1_qPCR_R |
| AGAGTGGAGAGATGATGGTCGA | CRY_II.1_qPCR_F |
| AGCCCTCCTGCCTCACTT | CRY_II.1_qPCR_R |
| CCCAGGTCAGCTTTCCCATT | DASH_qPCR_F |
| ATCCCAGAGCAAGCCATGTT | DASH_qPCR_R |
| ACACAGATTGCCCTTGTTCTCT | ASO_I-1_qPCR_F |
| CGGTCGTCGTCTTCACCCTA | ASO_I-1_qPCR_R |
| CCGCCATTTCTATCGACCGT | ASO_I-2_qPCR_F |
| GCTCTTGTCCGCGTAATCCT | ASO_I-2_qPCR_R |
| ACCCAAGATTTCGAGCAGATGT | ASO_II-1_qPCR_F |
| CCTTCTCCACGCGGTCTAAG | ASO_II-1_qPCR_R |
| GAGTGCTCAAGCGATGTCCT | ASO_II-10_qPCR_F |
| GCTGTTTAATTCCAGTGACCTGT | ASO_II-10_qPCR_R |
| ACGACGATTTCAGTCGCCAA | ASO_II-11_qPCR_F |
| TGGGGGAATGGATAAAACGAGC | ASO_II-11_qPCR_R |
| CAAGTACTCTGTACAGCCCGTT | ASO_II-12_qPCR_F |
| GACACTTGCTCGAAACCTTGGA | ASO_II-12_qPCR_R |
| GCACTGGTAAATCCAAGGTTTCG | ASO_II-2_qPCR_F |
| ATCTTCTCGCGGACGAAGTG | ASO_II-2_qPCR_R |
| GTAGAAATGTGGCCGACGCT | ASO_II-3_qPCR_F |
| GTCTCGACGGAATCTTGGGT | ASO_II-3_qPCR_R |
| GTTTGGCGGGAGGAATTTGC | ASO_II-4_qPCR_F |
| ACCACAATTTTTCGAAATCTGCCA | ASO_II-4_qPCR_R |
| GAGAGGCGTACAGGTCCAAA | ASO_II-5_qPCR_F |
| CCTCGTAAACCAGTGTGCCT | ASO_II-5_qPCR_R |
| GACACCAACCACGGCCTTTA | ASO_II-6_qPCR_F |
| ACTTTGACCATAACCCTAAAGCGA | ASO_II-6_qPCR_R |
| CACAGGATAAGCGTCGTCAGT | ASO_II-7_qPCR_F |
| CGCAGTACGGAGACCAACTT | ASO_II-7_qPCR_R |
| GCTCCTGACTGGACATCGAC | ASO_II-8_qPCR_F |
| TTCTTCAGGGTCAACCTATACAGC | ASO_II-8_qPCR_R |
| GGGAACAGCAAGGGTCAGTT | ASO_II-9_qPCR_F |
| GCAAAAGGGCCCTGTAGATTTG | ASO_II-9_qPCR_R |
| AAACCAGGGTTTTAAAGATCGAAAC | Cnidopsin-1_qPCR_F |
| TGACGGCAATAACAAGACCA | Cnidopsin-1_qPCR_R |
| CACACTCGAAAAAGGCCACA | Cnidopsin-2_qPCR_F |
| TCTTAGGCCTTAAGTCGTCGT | Cnidopsin-2_qPCR_R |
| GAACGTACCTGGAGGTCAGC | Cnidopsin-3_qPCR_F |
| TCGTTTTGTTGGGAGCGACT | Cnidopsin-3_qPCR_R |
| TGCCAGGAATTGCTGAACCAT | Cnidopsin-4_qPCR_F |
| GTGTTGAGTACGAACGACCCT | Cnidopsin-4_qPCR_R |

**PHR region duplication**

| | |
|---|---|
| CTCCCAGCGATCTCCAAAGG | AnthoCRY.1_E7-E8_F |
| GTGGTCCAAGCTTGCGAATG | AnthoCRY.1_E7-E8_R |
